# Double-stranded RNA drives SARS-CoV-2 nucleocapsid protein to undergo phase separation at specific temperatures

**DOI:** 10.1101/2021.06.14.448452

**Authors:** Christine A. Roden, Yifan Dai, Ian Seim, Myungwoon Lee, Rachel Sealfon, Grace A. McLaughlin, Mark A. Boerneke, Christiane Iserman, Samuel A. Wey, Joanne L. Ekena, Olga G. Troyanskaya, Kevin M. Weeks, Lingchong You, Ashutosh Chilkoti, Amy S. Gladfelter

**Affiliations:** Department of Biology, University of North Carolina at Chapel Hill, Chapel Hill, NC, USA; Lineberger Comprehensive Cancer Center, University of North Carolina at Chapel Hill, Chapel Hill, NC, USA; Department of Biomedical Engineering, Duke University, Durham, NC 27708; Curriculum in Bioinformatics and Computational Biology, University of North Carolina at Chapel Hill, Chapel Hill, USA; Department of Applied Physical Sciences, University of North Carolina at Chapel Hill, Chapel Hill, USA; Laboratory of Chemical Physics, National Institute of Diabetes and Digestive and Kidney Diseases, National Institutes of Health, Bethesda, MD 20892-0520.; Flatiron Institute, Simons Foundation, New York, NY, USA; Department of Chemistry, University of North Carolina at Chapel Hill, Chapel Hill, NC, USA; Department of Computer Science, Princeton University, Princeton, NJ USA; Lewis-Sigler Institute for Integrative Genomics, Princeton University, Princeton, NJ USA; Center for Genomic and Computational Biology, Duke University, Durham, NC 27708; Department of Molecular Genetics and Microbiology, Duke University School of Medicine, Durham, NC27708

**Keywords:** SARS-CoV-2, nucleocapsid, liquid-liquid phase separation (LLPS), double-stranded RNA (dsRNA), RNA stickers and spacers, lower critical solution temperature (LCST), translational repression, viral genome packaging, RNP formation.

## Abstract

Betacoronavirus SARS-CoV-2 infections caused the global Covid-19 pandemic. The nucleocapsid protein (N-protein) is required for multiple steps in the betacoronavirus replication cycle. SARS-CoV-2-N-protein is known to undergo liquid-liquid phase separation (LLPS) with specific RNAs at particular temperatures to form condensates. We show that N-protein recognizes at least two separate and distinct RNA motifs, both of which require double-stranded RNA (dsRNA) for LLPS. These motifs are separately recognized by N-protein’s two RNA binding domains (RBDs). Addition of dsRNA accelerates and modifies N-protein LLPS in vitro and in cells and controls the temperature condensates form. The abundance of dsRNA tunes N-protein-mediated translational repression and may confer a switch from translation to genome packaging. Thus, N-protein’s two RBDs interact with separate dsRNA motifs, and these interactions impart distinct droplet properties that can support multiple viral functions. These experiments demonstrate a paradigm of how RNA structure can control the properties of biomolecular condensates.

## Introduction

Liquid-liquid phase separation (LLPS) is a mechanism of macromolecular self-assembly that results in the formation of micron-scale droplets which are thought to contribute to numerous cellular functions (Alberti et al., 2019; Boeynaems et al., 2018; Hyman et al., 2014). While many of the mechanisms for protein-based condensation into droplets are known the rules for partitioning specific nucleic acids are largely undefined. A model of “stickers and spacers” describes many phase separating and percolation systems in which “stickers” represent sites of interactions amongst polymers and “spacers” are the intervening sequences between the association sites (Choi et al., 2020; Rubinstein and Semenov, 1998; Semenov and Rubinstein, 1998). The grammar of protein-protein interaction “stickers” amongst disordered proteins and oligomerization domains is beginning to be established (Bremer et al., 2021; Choi et al., 2019, 2020; Martin et al., 2020; Nott et al., 2015; Vernon et al., 2018; Wang et al., 2018, 2021). How “stickers” are encoded for RNA-protein or RNA-RNA interactions to promote condensates of specific identity and properties is far more mysterious (Roden and Gladfelter, 2021).

Viruses present an opportunity to dissect LLPS-promoting interactions between proteins and nucleic acids because of their limited proteome that must engage with specific viral nucleic acids (i.e., viral genome). Indeed, proteins and nucleic acids from many different viruses have now been shown to undergo LLPS (Brocca et al., 2020; Guseva et al., 2020; Heinrich et al., 2018; Nikolic et al., 2017; Rincheval et al., 2017). Importantly, viral model systems involving one LLPS promoting protein and one genomic nucleic acid (such as RNA), can reveal new principals for nucleic acid sequence- and structure-encoded LLPS. This is because chemical complexity is reduced by having a single protein and specific nucleic acids for viral droplets rather than the complex mixtures that are present in other cellular droplets. We predict that viruses store information in their nucleic acid sequence and RNA structure to encode different LLPS dependent functions to achieve biochemical complexity. In this study, we manipulate RNA sequence and structure to decode RNA features that specify LLPS for SARS-CoV-2 nucleocapsid (N-protein) and genomic RNA.

Although, the global COVID-19 pandemic motivated many studies of N-protein LLPS, the specific role(s) of LLPS in the viral replication cycle is still an open problem. The nucleocapsid protein (N-protein) is required for multiple viral functions (McBride et al., 2014). N-protein has many features associated with proteins that undergo LLPS including two RNA binding domains (RBD1 and RBD2) and additional intrinsically disordered motifs (Chang et al., 2014). Notably, N-protein displays lower critical solution temperature behavior (LCST), and RNA tunes the temperature at which N-protein undergoes LLPS (Iserman et al., 2020). N-protein undergoes LLPS both in cells and cell-free (Carlson et al., 2020; Cascarina and Ross, 2020; Chen et al., 2020; Cubuk et al., 2021; Iserman et al., 2020; Jack et al., 2020; Lu et al., 2021; Perdikari et al., 2020; Savastano et al., 2020; Wang et al., 2021; Zhao et al., 2021b) with fragments of the viral RNA genome (Carlson et al., 2020; Iserman et al., 2020; Lu et al., 2021). N-protein LLPS is also dependent on salt (Jack et al., 2020; Lu et al., 2021; Perdikari et al., 2020), pH (Perdikari et al., 2020), and RNA sequence (Carlson et al., 2020; Iserman et al., 2020; Jack et al., 2020; Lu et al., 2021). Although RNA is required to induce N-protein LLPS at physiological temperatures and ion conditions, the RNA sequence and structural preferences that govern N-protein interactions with RNA are unknown. Remarkably, RNAs of the same length but different sequence and structure do not equally drive N-protein LLPS (Iserman et al., 2020). This specificity indicates that N-protein LLPS is encoded by sequence- and structure-specific interactions with RNA and raises the possibility that during infection, distinct N-protein functions occur in molecularly distinct droplets.

N-protein phase separation shows remarkable specificity for RNA sequence but the mechanism for N-protein’s recognition of RNA is unknown. We previously showed that the first 1000 nucleotides of the SARS-CoV-2 genome (termed 5′end RNA) reliably drive N-protein LLPS. In contrast, another RNA sequence of identical length surrounding the frameshifting element (FS) promoted solubilization of N-protein (Iserman et al., 2020). A clue to these opposing effects came from the observation that these two RNAs exhibited differential crosslinking patterns with N-protein. Crosslinking between the 5′end RNA and N-protein preferentially occurred in specific single-stranded areas adjacent to structured elements. In contrast, crosslinking was uniformly distributed in the solubilizing FS sequence. We speculated that the differential crosslinking between LLPS-promoting and solubilizing RNA could be used as a tool to identify N-protein preferences for particular RNA sequences and give insights into how different modes of protein-RNA interactions influence LLPS. Thus, we sought to uncover how N-protein recognizes RNA to lead to LLPS.

We show that the two RNA-binding domains in N-protein interact with distinct RNA-sequence and structure elements. This indicates N-protein has at least two distinct types of protein-RNA interaction “stickers” that could provide multivalency for LLPS. RBD1 recognizes transcription-regulating sequence (TRS) and similar sequences in an RNA structure dependent manner. RBD2 specifically interacts with dsRNA and these interactions are critical for specifying the LLPS temperature. The patterning of N-protein RBD1 and 2 binding “stickers” in the RNA sequence regulates LLPS to specify condensation rate, RNA translation efficiency and genome condensation. Our work expands the understanding of RNA features which promote N-protein binding and provides to our knowledge, the first evidence of dsRNA/RBD interactions in regulating LLPS and LCST behavior. Importantly, we identify how combinations of the two dsRNA stickers can pattern protein-RNA interactions to regulate LLPS with important implications for betacoronavirus replication.

## Results

We sought to determine which RNA features promote N-protein LLPS using an in vitro LLPS assay. Previously, we identified two regions within the first 1000 nucleotides (nt) or 5′end of SARS-CoV-2 which preferentially crosslinked with recombinant N-protein at protein concentrations below the critical concentration for phase separation, which we termed principal sites. These principal sites are in single stranded sequences between two strongly structured (**Fig. 1A**) and conserved (**Fig. 1B**) stem-loops. Our goal here is to understand which features of this RNA sequence are the interaction sites of N-protein relevant for driving LLPS. We hypothesize that these principal sites either act as “stickers” that drive co-phase separation with N-protein or are “spacers” adjacent to the functional stickers in the structured elements.

**Fig 1.**
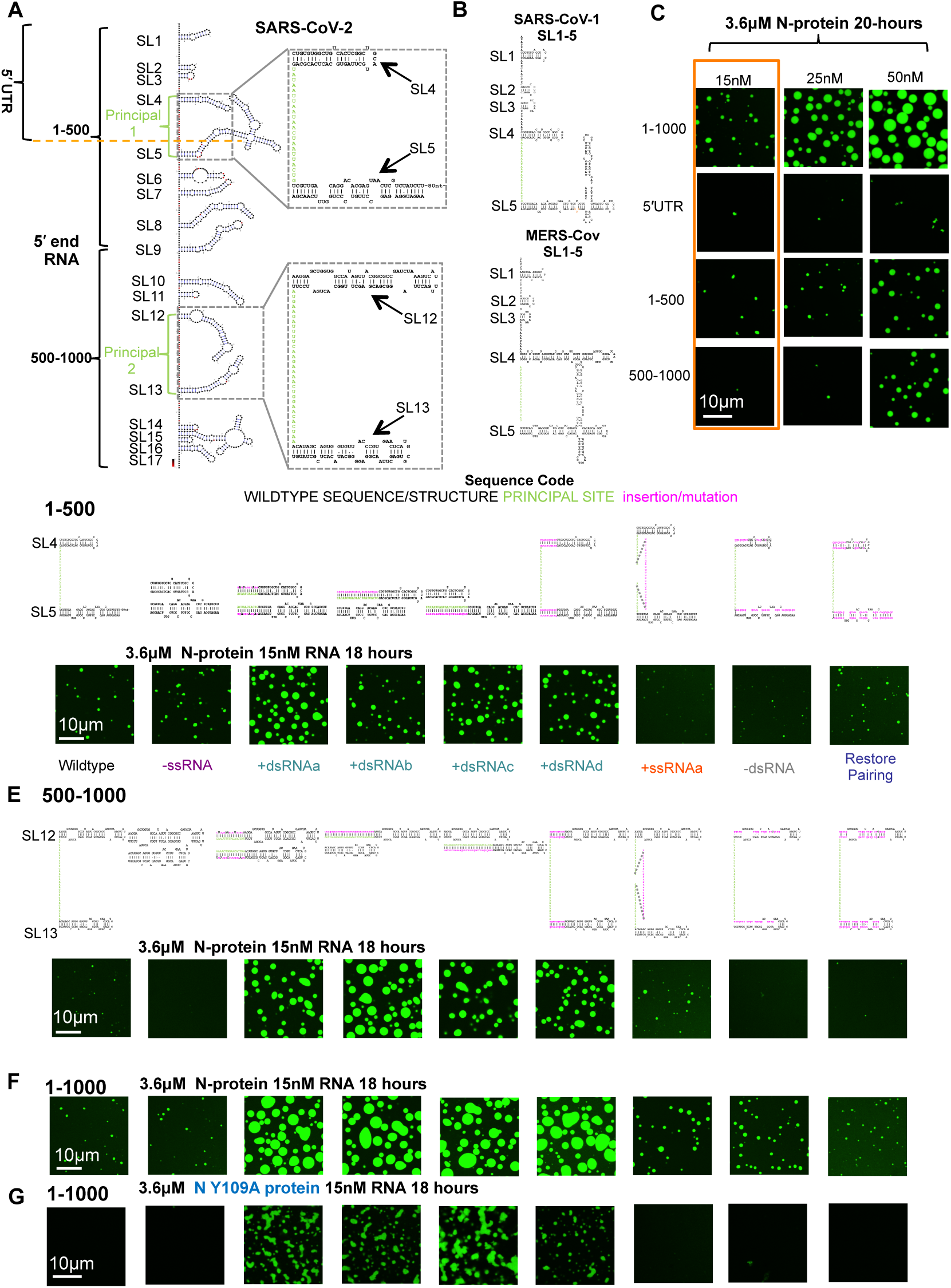
dsRNA-driven LLPS is independent of RBD1. **(A)** SHAPE based structure model of the first 1000 nucleotides of the SARS-CoV-2 genome. Light green letters indicate locations of preferential N-protein crosslinking (principal sites). Brackets indicate the fragments; 5′UTR, 1-500nt and 500-1000nt. Stem-loops are numbered (SL). Inset indicates locations of structure manipulations for the rest of the figure. Specifically, mutations altered the region containing SL4 and 5 of principal 1 and/or SL12 and 13 principal 2. **(B)** Comparison of SL1-5 of SARS-CoV-1 and MERS-CoV(Sun et al., 2021). **(C)** Representative images from LLPS experiments with 3.6μM recombinant N-protein (green) and the corresponding RNA sequence 1-1000, 5′UTR, 1-500, and 500-1000 for 15, 25, and 50nM RNA. Orange box indicates selected condition for (Fig. 1D-F). **(D)** Mutation series in the 1-500 context depicting the predicted structure of mutants directed against SL4 and 5 and the intervening single stranded sequence of principal site 1 (light green letters). N-protein is depicted in green. Mutation classes are as follows -ssRNA (purple), +dsRNA (teal), +ssRNA (orange). -dsRNA (grey), Restore pairing (blue). **(E)** The equivalent mutation series (as in Fig. 1D) for 500-1000 context (principal site 2 in light green letters) depicting the predicted structure of mutants directed against SL12 and 13 or the intervening single stranded sequence of principal site 2. **(F)** Combination of mutations from **1D** and **E** in the context of 1-1000. N-protein is depicted in green. **(DEF)** Deletion of the single stranded regions of the principal sites do not significantly impact LLPS (-ssRNA). Addition of dsRNA (teal) (+dsRNAa-d) enhances N-protein LLPS. Addition of single stranded RNA (+ssRNAa orange) coding for HA tag in the center of the principal sites leads to a mild enhancement of LLPS. Unpairing principal site adjacent stem-loops (grey -dsRNA) on the 5′ side reduces LLPS. Restoration of wildtype RNA structure (blue Restore pairing) but with a different sequence restores LLPS to wildtype levels. **(G)** Only those mutations who lead to an addition of dsRNA (+dsRNAa-d), retain the ability to induce phase separation following Y109A mutation and destruction of N-protein RBD1. For all images, scale bar indicates 10μm all experiments show representative images from at least 3 replicates and 2 independent batches of RNA.

We wanted to be able test each principal site independently and in combination in conditions which allowed us to see droplet size and/or morphology change following mutation. Principal site 1 is located in the 5′UTR (nt:1-267 above orange dashed line) **(Fig. 1A**). Given the observation that 5′UTR and smaller fragments can induce N-protein phase separation (Carlson et al., 2020; Lu et al., 2021), we first asked what segments of the first 1000 nucleotides (nt) of the genome (5′end) were sufficient to promote LLPS? To this end, we tested 1-1000nt, 1-267nt (the 5′UTR), 1-500nt, and 500-1000nt fragments at either 15, 25, or 50nM RNA and 3.6μM protein. All tested fragments could drive N-protein LLPS, however some fragments drove LLPS more readily (1-500 or 1-1000) (**Fig. 1C**). We selected 3.6μM N-protein and 15 nM RNA for subsequent experiments (orange box), which resulted in medium sized droplets for 1-1000nt and 1-500nt. Medium sized droplets allowed us to see reduction or enhancement of LLPS following mutation.

We predict if N-protein recognizes ssRNA, altering the ssRNA content between stem-loops (increasing or decreasing) should alter N-protein binding to principal sites and in turn LLPS. Alternatively, if adjacent dsRNA mediates N-protein recognition of principal sites, we predict that changing the length of stem-loops will alter N-protein binding and LLPS. Thus, we designed a series of mutations to independently disrupt single stranded and double-stranded RNA in or adjacent to the principal sites.

### dsRNA promotes N-protein LLPS

We disrupted principal site 1 in 1-500nt (**Fig. 1D**), principal site 2 in 500-1000nt (**Fig. 1E**), or both principal sites in 1-1000nt (**Fig. 1F**). We first wanted to examine whether dsRNA or ssRNA was important for N-protein crosslinking to principal sites. To test the importance of the single-stranded, principal site sequence alone, we first deleted the single stranded sequence (-ssRNA). We observed that in any of the tested sequence contexts (**Fig. 1D-F**) deletion of the principal site did not significantly alter LLPS relative to wild-type. This shows that the ssRNA is not required for N-protein LLPS. Instead, N-protein binding to dsRNA may drive LLPS.

We next sought to address the role of the conserved structured RNA located adjacent to the principal sites. To do this, we converted the single-stranded, principal site sequence to dsRNA (preserving the total RNA length, by recoding the sequence 5’ and 3′ to the stem-loops to pair with the principal site (+dsRNAa). Strikingly, this type of mutation resulted in enhanced LLPS in all three sequence contexts (**Fig. 1D-F**), with much larger droplets forming more quickly in identical protein and RNA concentrations. We next sought to induce the formation of additional double-stranded RNA in a different way. Thus, we converted the single stranded principal site region to double-stranded RNA by forcing the single stranded region to base pair by adding complementary sequence either 5′ of the first hairpin (+dsRNAb) or 3′ of the second hairpin (+dsRNAc) which flanked the single stranded principal sites. In all three sequence contexts, this type of mutation again enhanced LLPS (**Fig.1 D-F**).

To examine if dsRNA addition could be additive, we also tested individually +dsRNAb and +dsRNAc on a single principal site (either 1 or 2) in the context of 1-1000nt. We observed that mutations which affect principal site 2 were better able to enhance LLPS compared to those which effect principal site 1 **(Fig. S1A**). We think this is likely due to dsRNA length differences. For example, mutated principal site 1 adding 44nt of dsRNA (22nt of additional RNA sequence) was less good at driving LLPS than principal 2 adding 62nt of dsRNA (31nt of additional RNA sequence) **(Fig. S1A**). Further, the combination of the mutations did not enhance LLPS much more than those only altering principal site 2 alone which indicates there may be a threshold to the enhancement **(Fig. S1A**). These data suggest that increasing dsRNA content, up to a certain threshold, can drive enhanced N-protein LLPS.

So far, all tested mutations which enhanced structure also destroyed the single stranded principal site by converting it to dsRNA. Therefore, we next asked whether addition of 10 paired nucleotides (20nt per stem-loop) of dsRNA at the base of the principal site flanking stem-loops would also promote LLPS (+dsRNAd) as this mutant would preserve the ssRNA of the principal site while creating additional structure. We observed that these extended stem-loops indeed also enhanced LLPS relative to wildtype in all three sequence contexts (**Fig. 1D-F**). A caveat to interpreting these results however is that 3/4 classes of +dsRNA (+dsRNAb,c,d) mutant RNA increases the length by 22-80 nt) and RNA length has been shown to modulate the ability of N-protein to undergo LLPS (Iserman et al., 2020).

We sought to disentangle the effects of length addition to the dsRNA constructs by inserting single stranded sequence (+ssRNAa) (27nt coding for Hemagglutinin (HA)) in the center of the single stranded principal sites. We observed in all three sequence contexts, addition of HA RNA sequence resulted in negligible enhancement of LLPS relative to wildtype (**Fig. 1D-F**). Two different ssRNA sequences were independently inserted into principal site 2 or the 3′ end. These RNA length controls all resulted in negligible levels of enhancement (**Fig.S1B**). Taken together, these results suggest that dsRNA addition enhances N-protein LLPS more than ssRNA addition and the enhancement by additional dsRNA cannot be explained simply by increased RNA length.

Next, we sought to determine whether the sequence and/or structure of the stem-loops flanking the principal sites were important. Therefore, we unpaired the principal site flanking stem-loops by making mutations (-dsRNA) on the 5′ side. We observed that in all three contexts -dsRNA resulted in a reduction (1-500) or loss (500-1000, 1-1000) of LLPS relative to wildtype (**Fig. 1D-F**). To rescue the -dsRNA mutant we made compensatory mutations on the 3′ side of the stem-loop to restore the structure (Restore pairing). The Restore pairing mutant resembled wildtype levels of LLPS in all sequence contexts. Thus, we concluded that unpairing of the principal stem-loops generally reduces LLPS and the specific primary sequence of the stem-loops does not play a significant role in LLPS.

To assess if addition of dsRNA was sensitive to the relative stoichiometry of RNA and protein, we tested wildtype and +dsRNAa in the context of 1-1000 in a small phase diagram. +dsRNAa was chosen as it is the same exact length as wildtype but drove more LLPS at 3.6μM N-protein and 15nM RNA. We observed that relative to wildtype **(Fig. S1C**), +dsRNAa (**Fig. S1D and E**) consistently drove more LLPS at 3μM N-protein (**Fig. S1C-D**) indicating this enhancement is reproducible in multiple regimes. However, differences were observed at 1μM N-protein with only some conditions driving more LLPS, indicating a shifted phase boundary for the mutant. (**Fig. S1C-D).** We further confirmed that N-protein recruitment to droplets was higher for +dsRNAa by measuring the absorbance of the diffuse phase at 280nm (A280) following the phase separation assay (**Fig. S1E**). In all three tested RNA concentrations at 3μM N-protein mutant RNA addition resulted in significantly higher A280 signal indicative of higher levels of droplet protein recruitment (**Fig. S1E**).

### dsRNA-driven LLPS is independent of RBD1

We next determined which RNA binding domain of N-protein mediates the dsRNA-based LLPS enhancement. N-protein has two distinct RNA-binding domains; RBD1 is structured (Kang et al., 2020) and RBD2 is a lysine-rich IDR (Zinzula et al., 2021). The single point mutant Y109A in RBD1 blocked LLPS with 5′end RNA (1-1000) (Iserman et al., 2020) and resulted in a 2000-fold reduction in affinity for RNA (Kang et al., 2020). Y109A mutant N-protein was incubated with the panel of mutant RNAs in the context of 1-1000. Only those mutations which resulted in more dsRNA could induce LLPS (**Fig. 1G**). Notably, the droplets that form with these more structured RNAs and Y109A are smaller and flocculated (different morphology) suggesting key aspects of the material properties of droplets are lost with the loss of RBD1 activity. Thus, +dsRNA can drive LLPS independent of a functional RBD1 N-protein suggesting +dsRNA requires RBD2.

We sought to test whether the LLPS-promoting mutations in the RNA sequences were specific to N-protein or generalizable to any RNA-driven phase separating system. To this end, we tested all mutations in the 1-1000 context with recombinant Whi3 protein. Whi3 has previously been shown to undergo sequence-specific RNA-dependent LLPS (Langdon et al., 2018; Zhang et al., 2015). We observed no obvious difference between any of the mutant RNAs and the wildtype 1-1000nt sequence with condensing Whi3 protein (**Fig. S1F**). This indicates that the mutations are specifically acting through alteration of N-protein/RNA LLPS and not a general, non-specific RNA:protein interaction or trans RNA:RNA interaction. Taken together, addition of dsRNA enhances LLPS of N-protein specifically, and this enhancement is independent of RBD1 and RNA primary sequence.

### RNA structure mutants accelerate droplet formation in cells and in solution

Next, we sought to confirm our observations regarding RNA sequence/structure-mediated N-protein LLPS in cells to see if the sequences behave similarly in the more complex and crowded cellular environment. To this end, we first needed to control for the reported translational repressive effects (Tidu et al., 2021; Yuan et al., 2020) of non-structural protein 1 (NSP1) which was encoded in our 5′end 1-1000 fragment. Thus, we designed a mutation in the start codon of NSP1 (Start Mutant) which would preserve the structure of SL5 but block NSP1 translation (**Fig. S2A**). We then confirmed that the Start Mutant yielded similar levels of LLPS as wild-type (**Fig. S2B**). It was unnecessary to also mutate the NSP1 start codon of our structure mutant of the same length (+dsRNAa) as this mutation also resulted in premature stop codons in NSP1 protein (**Fig. S2C**). Thus, we cloned wildtype 1-1000, Start Mutant, and +dsRNAa into a mammalian expression vector and co-transfected these plasmids with a plasmid driving N-protein: GFP in HEK293T cells (**Fig. S2D**).

To determine if dsRNA addition altered LLPS in cells, we imaged co-transfected cells. We observed that at early timepoints (24 hours) +dsRNAa resulted in a significant increase in the number of puncta (4-5 per cell) per square micron in cells compared to Wildtype or Start Mutant control (2-3 per cell) (**Fig.S2G**). However, this difference was reduced at 48 hours. Further, there was no significant difference in the mean fluorescence of N: GFP between the compared cells (**Fig. S2G**) so differences could not be explained by N-protein expression levels (Iserman et al., 2020). Collectively, these results suggest that dsRNA addition accelerates N-protein LLPS in cells.

The apparent acceleration in droplet formation time, prompted us to examine differences in timing with the in vitro system for mutants in all 3 sequence contexts (shorter incubation time (2 hours) than shown **in Figure 1 D-F** (18 hours)). Consistent with the structure mutants accelerating N-protein LLPS in cells, the mutants which result in more LLPS at 18 hours had vastly more pronounced differences at 2 hours indicating that these structure mutants also accelerate droplet formation cell free (**Fig. S2H**). Similar results as for 1-1000 **(Fig. S2H)** were obtained for both the 1-500 and 500-1000 context (**Fig. S2I and J)**. Collectively, these data suggest that in cells and cell free addition of dsRNA accelerates N-protein phase separation with target RNAs.

### TRS sequence/structure motif promotes N-protein LLPS

The data thus far show that a primary driver of LLPS is dsRNA/RBD2 interactions but there are several lines of evidence that suggest additional interactions are mediated by RBD1. First, Y109A mutant N-protein does not undergo LLPS with wildtype 5′end sequence. Second, the Y109A+dsRNA droplets have altered morphology suggesting some interaction between N and RNA has been lost in the absence of RBD1 activity. Thus, we sought to identify what RNA sequence features are favored by RBD1.

Given the transcriptional regulatory sequence (TRS/SL3) is the reported binding site of RBD1 in MHV (Grossoehme et al., 2009), and the 1-500 fragment which contains the TRS was better able to drive LLPS than the 500-1000nt fragment (**Fig. 1C**), we reasoned the TRS may be the preferred binding site of RBD1. Thus, we sought to characterize the importance of the TRS in N-protein LLPS (**Fig. 2**).

**Fig. 2.**
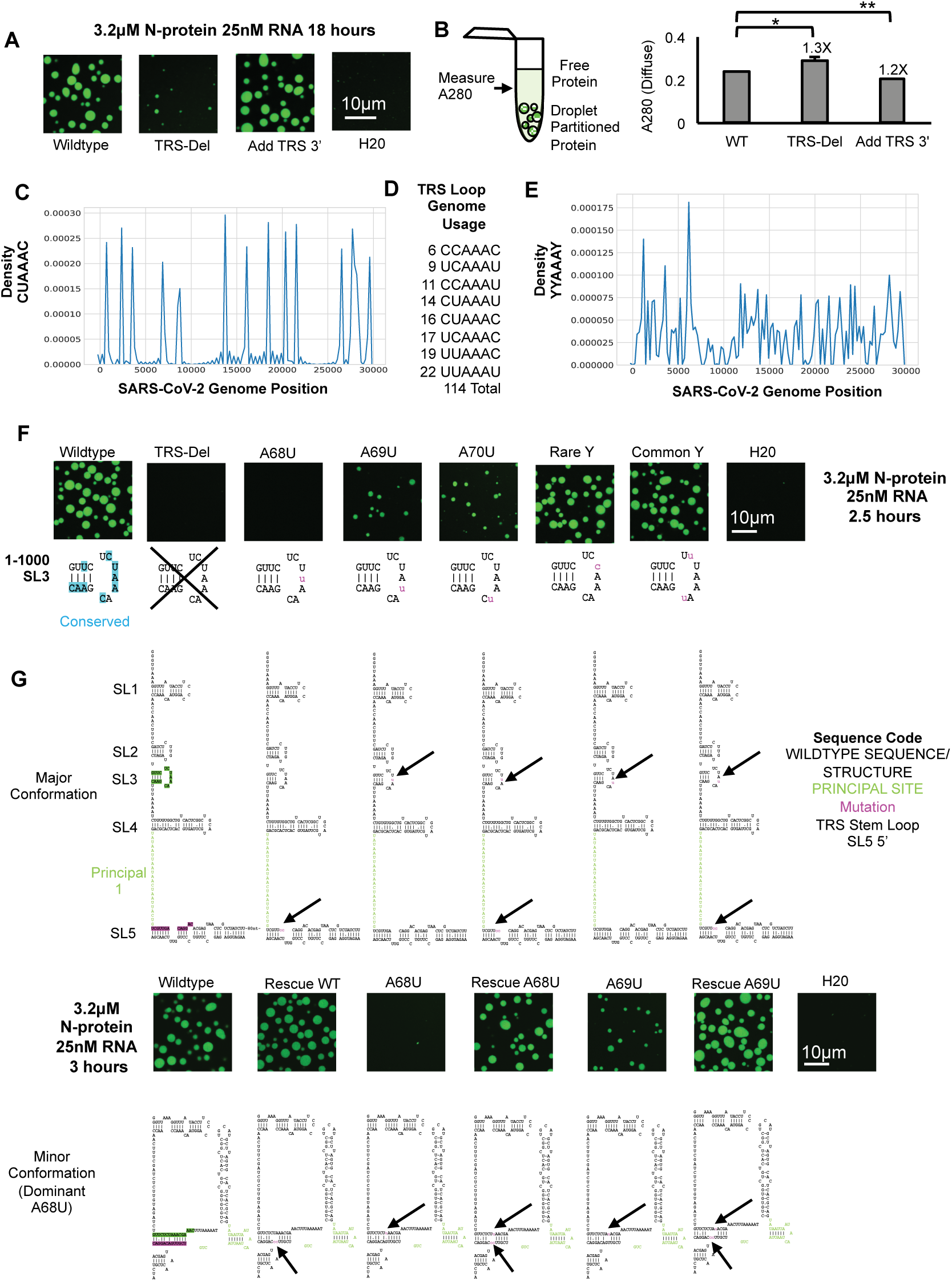
TRS sequence/structure motif promotes N-protein LLPS. **(A)** 3.2μM N-protein (green) and 25nM RNA following 18 hours of incubation for wildtype 1-1000, a mutation which deletes the entire TRS-stem-loop (TRS-del), or a mutation which appends an additional TRS to the 3′ end (Add TRS 3′). **(B)** Add TRS 3′ has lower A280 measurements then wildtype indicative of less protein in solution and more LLPS whereas TRS-del is the opposite. Error bars mark standard deviation for the three replicates and * indicate significance students T test (*** <0.001, ** <0.01, ns not significant) with brackets showing comparison for the indicated statistical test. **(C)** SARS-CoV-2 genome positions for TRS-loop sequence CUAAAC with Y axis indicating the density of that sequence per position in the genome (x axis). **(D)** Count of TRS-loop-like sequences (YYAAAY) in the genome where Y is equivalent to C or U. **(E)** Distribution of YYAAAY motif throughout the SARS-CoV-2 genome. **(F)** 3.2μM N-protein (green) and 25nM RNA following 2.5 hours of incubation for wildtype 1-1000, a mutation which deletes the entire TRS-stem-loop (TRS-del), A68U mutation, A69U mutation, A70U mutation and mutations which alter the sequence of the A flanking pyrimidines (Y’s) (C’s and U’s) to the most rare and common YYAAAY in the SARS-CoV-2 genome (see Fig. 2D). Deletion of TRS-loop or alteration of the AAA of the loop but not the Y’s leads to a reduction in LLPS. **(E)** TRS-loop mutations (A68 and 69U) exhibit effects primarily through structural rearrangement. Wildtype 1-1000 RNA is predicted to exist in at least two possible confirmations. The major structure which supports the TRS stem loop structure (SL3) and a minor confirmation which is conferred by interaction between the apical loop of the TRS stem-loop and the base of SL5. A68U/A69U are predicted to favor the minor conformation. Rescue mutation partially suppresses A68U and completely suppress A69U compared to Rescue Wildtype. 3.2μM N-protein (green) and 25nM RNA following 3 hours of incubation. For all images scale bar indicates 10μm all experiments show representative images from at least 3 replicates and 2 independent batches of RNA. H20 is water only no RNA added control. For RNA structure predictions principal site 1 is depicted in green letters and mutations are depicted in magenta lower-case letters.

To test if the presence of the TRS was required for LLPS, in the context of 1-1000nt, we deleted the entire TRS stem-loop (TRS-del) or added an additional TRS motifs to the 3′ end (Add TRS-3′). These mutations effectively increased or decreased the putative RBD1 binding site by one (**Fig. 2A**). TRS-del almost completely blocked LLPS (**Fig. 2A**) and reduced N-protein recruitment to droplets (more N-protein in the diffuse phase) (**Fig. 2B**). These results are analogous to how in the absence of RBD1 activity (Y109A mutation), 5′end RNA 1-1000 could no longer drive N-protein LLPS (Iserman et al., 2020)**. Fig. 1G**). Additionally, crosslinking experiments reveal reduced binding of Y109A N-protein at regions adjacent to the TRS-Loop (**Fig. S3A and B** (Iserman et al., 2020). Conversely, the addition of a TRS-loop (Add TRS-3′) resulted in slightly larger droplets than wildtype and enhanced N-protein recruitment to droplets (**Fig. 2B**). We conclude from these studies that the presence of TRS-loop facilitates N-protein LLPS with 5′end RNA.

We next wondered if N-protein could also bind sequences which were similar to the TRS-loop. This is because N-protein can drive LLPS with other genomic RNA sequences (Carlson et al., 2020; Iserman et al., 2020; Jack et al., 2020; Lu et al., 2021) We hypothesized that the most favored RBD1 binding site was YYAAAY (Y= C or U) which is similar to the TRS-loop sequence. In accordance with this hypothesis, crosslinking of N-protein is reduced in the region adjacent to the TRS-Loop for the RBD1 Y109A mutant N-protein (**Fig. S3A and B** (Iserman et al., 2020). The TRS-loop sequence, CUAAAC, occurs 16 times across the genome (**Fig. 2C**), and a chemically similar sequence YYAAAY (Y= C or U), a TRS-loop-like sequence, occurs 114 times across the genome **(Fig. 2D)**. Binding to this sequence is suggested to occur in MHV N-protein experiments (Grossoehme et al., 2009). Thus, we tested TRS-del, or mutated the individual As in the TRS-Loop sequence, CUAAAC, (A68U, A69U, A70U named so for the corresponding nucleotides in MHV). We also mutated the Cs and Us in the TRS-Loop sequence based on their occurrence in the SARS-CoV-2 genome. A68U, like TRS-del, completely blocked LLPS, and A69U and A70U resulted in a decrease relative to wildtype (**Fig. 2F**). Mutation of sequences to the rare (low frequency in the genome) or common Y (high frequency) sequence had negligible effects on N-protein LLPS (**Fig. 2F**). We conclude that while the A identity is critical for N-protein LLPS, the identity of the flanking Y sequence (C or U) is not. These results suggest that RBD1 may interact with TRS-loop-like sequences (YYAAAY) across the genome. Thus, individual mutations of specific, conserved nucleotides in the TRS-Loop sequence also resulted in reduced LLPS.

We next asked if the reduction of LLPS which occurred following primary sequence mutation was due to RNA structure-dependent alterations. This is because when we were designing A mutations, we noticed that the A68U/A69U mutations were predicted to favor a different structure than wildtype. (**Fig. 2G**). This predicted alternative structural arrangement pairs the TRS stem-loop (SL3, green) with SL5 (magenta). Thus, the A to U mutants’ impact on LLPS may occur via a sequence change, structure change or both. We reasoned we could make an allele that disentangles these confounding features by mutating nucleotides in SL5. This mutant would restore the wildtype structure while preserving the single nucleotide changes in the TRS-loop (Rescue mutants, **Fig. 2G**). Rescue wildtype RNA resulted in a mild enhancement of N-protein LLPS in comparison to the wildtype. In contrast, Rescue A68U almost completely restored the LLPS and Rescue A69U completely restored the LLPS. We conclude that the identity of A68 and A70 are important for N-protein RBD-1 binding. In contrast to MHV N-protein preferences, A68 and A69U further suppress LLPS by causing a structural rearrangement. Thus, the sequence and structure of the TRS stem-loop is required for N-protein binding and LLPS.

### RNA sequence and structure encode N-protein LCST behavior via RBD2

N-protein can phase separate in the absence of RNA at high temperatures but, RNA addition lowers LLPS temperature. Thus N-protein displays lower critical solution temperature (LCST) behavior with respect to RNA (Iserman et al., 2020). Notably, the addition of RNA, lowers the LCST to physiological temperatures. It is unclear if and how RNA sequence and structure dictates the temperature at which LLPS occurs. Given our observations from **Figures 1 and 2**, where we observed N-protein LLPS is controlled by RNA sequence/structure we wondered if LCST temperature was also controlled by RNA sequence/structure. To our knowledge, nothing is known about the role of RNA sequence in influencing LCST behavior in LLPS systems.

To test if LCST temperature is encoded by RNA sequence we first tested if N-protein condensation temperature changed as a result of the co-condensing RNA. Thus, we tested 3 different RNA sequences. To this end, we used a temperature-dependent ultraviolet-visible spectroscopy assay to map the saturation temperature, read out as turbidity with turbidity being a proxy for LLPS. We examined the following conditions: N-protein alone, N-protein + an RNA which does not drive LLPS (Frameshifting region RNA (FS)(Iserman et al., 2020), N-protein + 5′end RNA (1-1000nt) which drove LLPS, or N-protein + Nucleocapsid RNA (drives LLPS but is a longer sequence then 5′end). We used these different sequences to examine if all RNA sequences influence N-protein LCST to the same degree.

We observed that N-protein+FS (which does not drive LLPS) and N-protein alone underwent phase separation at the same high temperature (**Fig. 3A**) of ∼46°C. In contrast, the two LLPS-promoting RNAs both lowered temperature with Nucleocapsid RNA conferring a lower temperature then 5′end. The turbidity curves differ in shape depending on the specific RNAs such that LLPS-promoting RNAs display a more gradual turbidity increase. This may be due to heterotypic RNA-protein interactions. Thus, while different RNAs promoted different LCST behavior, this could be due to sequence and/or length-dependent effects.

**Fig. 3.**
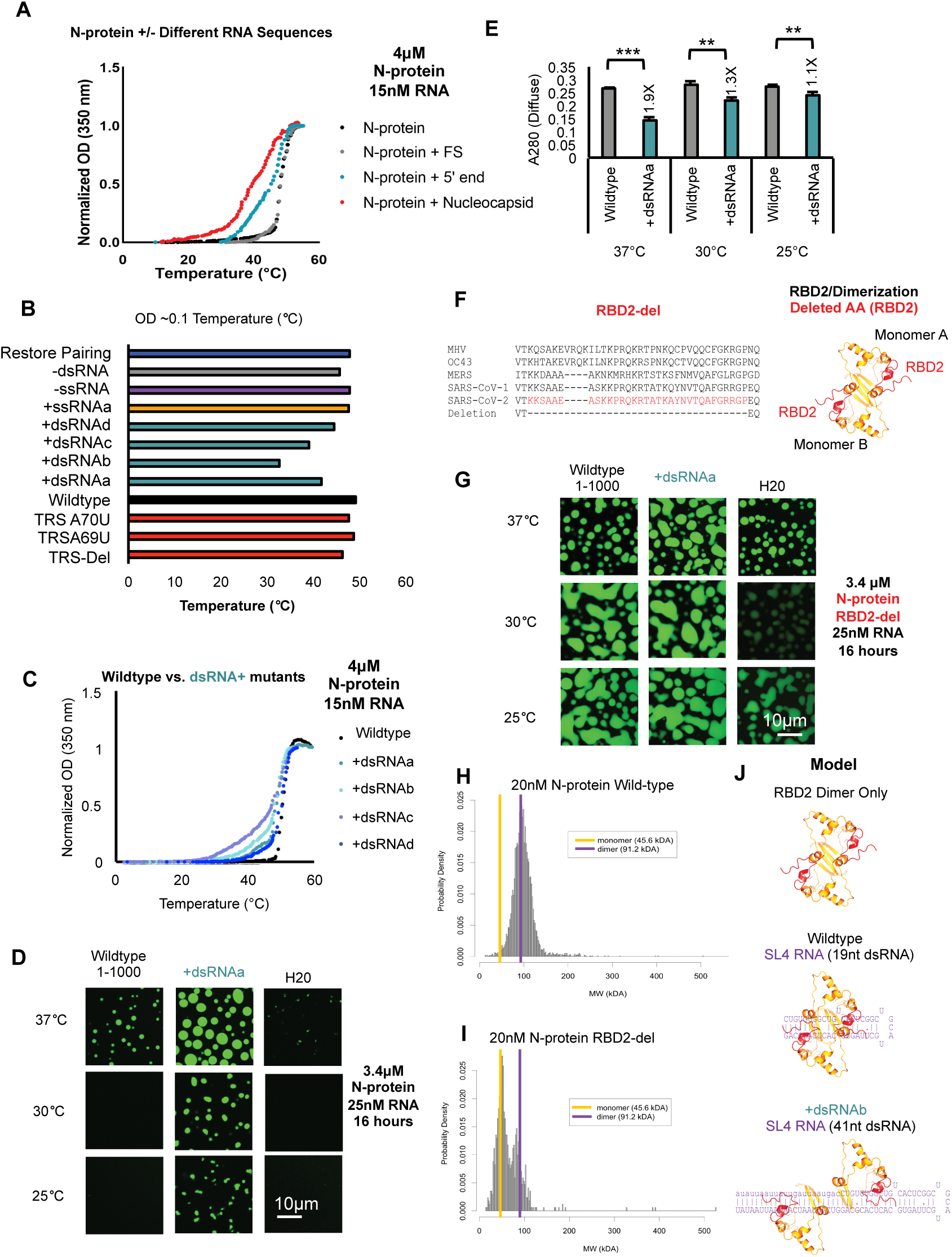
RNA sequence and structure encodes N-protein LCST behavior via RBD2. **(A)** Temperature dependent turbidity tests of N-protein alone (Black), N-protein with Frameshifting region RNA (FS) (Gray), and N-protein with 5′end RNA (1-1000nt) (Blue) and N-protein with Nucleocapsid RNA (Red). Addition of droplet forming RNAs, 5′end 1-1000nt and Nucleocapsid RNA to N-protein, lowers the transition temperature but solubilizing RNA (FS) does not**. (B)** Transition temperature comparison (repeat of the experiment shown in **(A)** of wildtype 5′end or 11 mutants in the context of 1-1000nt. Bar length indicates the temperature in °Celsius at which the turbidity of the solution reaches ∼0.1. Only those mutants which alter the dsRNA content (teal +dsRNA), lower the temperature at which OD reaches ∼0.1 indicative of increased solution turbidity. **(C)** Temperature dependent turbidity tests for N-protein plus wildtype 5′end RNA as well as the 4 more structured mutants (+dsRNA) which lower the transition temperature. **(D)** Validation of the turbidity assay using droplet imaging (Fig. 3B **and C**). 3.4μM Wildtype N-protein was mixed with either 25nM of wildtype 5′end 1-1000 RNA, +dsRNAa (RBD1 independent Fig. 1G) or water only added control (H20) and incubated at the indicated temperature 37, 30, or 25°C for a period of 20 hours prior to imaging. Consistent with previous results, +dsRNAa increases droplet size relative to wildtype at 37°C (Fig. 1F) & induces LLPS at lower temperatures. **(E)** A280 measurement of remaining N-protein in the diffuse phase for Fig. 3D. At all temperatures +dsRNAa lowers A280 measurements relative to wildtype. Error bars mark standard deviation for the three replicates and * indicate significance students T test (*** p<0.001, ** p<0.01, *p<0.05, ns not significant) with brackets showing comparison for the indicated statistical test. **(F)** Protein sequence conservation of N-protein RBD2 and structure model of the RBD2 dimerization domain for SARS-CoV-2 (red sequences/ red ribbon) indicate the location of the deletion in the primary sequence tested in **(G)**. **(G)** RBD2/Dimerization domain is required for proper N-protein LCST behavior at indicated temperature range. 3.4μM of N-protein RBD-del (green) was mixed with 25nM of either wildtype 1-1000, +dsRNAa, or water only control and incubated at the indicated temperatures for 16 hours. Droplet formation was observed in all conditions although RNA dependence was more evident at lower protein concentrations (**Fig. S4 G-H**). **(H and I)** Mass photometry histograms showing the molecular weight (MW) distribution of detected particles for wild-type N-protein **(H)** or RBD-del N-protein **(I). (H)** Wildtype N-protein is a stable dimer in solution (250mM NaCl pH 7.5 20mM phosphate buffer 20nM N-protein) but RBD2-del is mostly a monomer **(I). (J)** Model of N-protein RBD2/Dimerization domain interactions with dsRNA. Binding of the two RBD2s of the two monomers of N-protein to dsRNA facilitates dimerization dissociation with temperature facilitating dissociation for shorter stem-loops. For all images scale bar indicates 10μm all experiments show representative images from at least 3 replicates.

We wanted to identify the RNA and protein features which were responsible for conferring N-protein LCST independent of RNA length (Nucleocapsid RNA is longer then 5′end). To disentangle the effects of RNA length and RNA sequence we tested the 1-1000nt mutant RNAs (similar or identical lengths only very slightly different sequences and structure) with N-protein (**Fig. 3B**). Notably, only those mutations which resulted in more secondary structure (+dsRNAa-d) significantly lowered the LCST **(Fig. 3B and C**). Loss of the putative RBD1 binding site (TRS) had no significant impact on temperature. Additionally, raising the temperature resulted in mildly enhanced N-protein binding to 5′UTR RNA in RBD1 deficient (Y109A mutant) protein via EMSA **(Fig. S4A and B)** suggesting N-protein’s RNA binding activity at higher temperatures is independent of RBD1.

To confirm that RBD1 activity did not alter LLPS temperature, we additionally tested the Add TRS 3′ mutation which creates an additional RBD1 binding site. Add TRS 3′ did not lower the LCST (**Fig. S4C and D**) via either a microscopy assay or diffuse phase measurement. These results suggested temperature sensitivity was conferred by RBD2/dsRNA interactions rather than generally enhanced binding to RNA (through RBD1 for example).

We confirmed the temperature-dependent turbidity results reflected the formation of droplets by examining assemblies under the microscope. +dsRNAa (one of the add more structure mutants with the same length as 1-1000 wildtype **Fig. 3B**) lowered N-protein LLPS temperature to 25°C (**Fig. 3D**). We confirmed that the diffuse phase measurement was perfectly anti-correlated with the imaging of the droplets at all temperatures (**Fig. 3E**). This further suggests that dsRNA-dependent interactions can tune the LCST.

Given that the more structured mutants do not require RBD1 activity to undergo LLPS (**Fig. 1G**) but all other tested RNAs do, we reasoned that the LCST behavior of N-protein could be specified by RBD2. To assess this possibility, we purified N-protein with a deletion in RBD2 (red amino acids) (**Fig. 3F**) which was predicted to preserve the conserved dimerization interface, (**Fig. S4E).** We observed that N-protein RBD-2-Del’s LCST behavior was significantly altered with both wildtype and mutant RNA (**Fig. 3G**) and could undergo LLPS at all tested temperatures even without additional RNA. Further, similar levels of protein in the diffuse phase (A280 signal) were detected following the LLPS assay (**Fig. S4F**). Reducing the N-protein and RNA concentration showed some degree of RNA dependence for the RBD-2-Del mutant (**Fig. S4G and H**). These data support that RBD2 interactions encode the temperature threshold for LLPS in this system.

We were surprised, however, to see that RBD2 deletion leads to increased LLPS. Based on the literature of SARS-CoV-1 N-protein RBD2 crystal structure (Chen et al., 2007), we hypothesized that RBD2-del region may stabilize the formation of higher order oligomers of N-protein. To address if RBD2-del mutation was destabilizing the formation of N-protein dimers (the reported oligomerization state of N-protein in the absence of nucleic acid (Zeng et al., 2020; Zhao et al., 2021a) we performed mass photometry. We observed that, consistent with previous studies, wildtype N-protein forms a dimer (**Fig. 3H**) whereas RBD-2 del is mostly a monomer (**Fig. 3I**). We conclude from this that the RBD2-del mutation destabilizes the N-protein dimer which may lead to reduced temperature and less RNA dependence of LLPS. dsRNA addition can similarly perform the dimer destabilization (splitting) but only for wild-type N-protein (**Fig. 3J**). Collectively, these data suggest that while RBD1 is required for 5′end to phase separate, RBD2 is required for LCST behavior at physiologically-relevant temperature and salt, and RBD2 encodes LCST behavior through preferentially binding dsRNA.

### N-protein binding and LLPS represses translation

Given the +dsRNA structure mutant promoted puncta formation in cells **(Fig. S2)** and the SARS-CoV-2 genome is enriched in dsRNA, even in protein coding sequences (Huston et al., 2021; Lan et al., 2021; Sun et al., 2021), we next asked if N-protein binding and LLPS could regulate target RNA translation. We reasoned the increase in LLPS due to dsRNA addition may be antagonistic to translation as some condensates can repress translation (Kim et al., 2019; Tsang et al., 2019). Thus, an understanding of how N-protein regulates translation would be informative for the viral life cycle.

To address N-protein mediated protein translational regulation encoded by dsRNA, we sought to replicate our dsRNA/ssRNA addition experiments (**Figure 1**) in the context of the 5′UTR (**Fig. 4A**) by altering stem-loop 4 (SL4). We observed that only the +dsRNAb (which results in 22nt additional sequence and 44nt of additional dsRNA) drove significant additional LLPS relative to wildtype (**Fig. 4A**). +dsRNAd which adds 10 paired nucleotides to the base of SL4 (20 additional nucleotides total) also resulted in minor enhancement. All other non +dsRNA mutants had negligible effects. Thus, length dependent addition of dsRNA to the 5′UTR should be sufficient to enhance LLPS independent of the coding sequence when appended in cis.

**Fig. 4.**
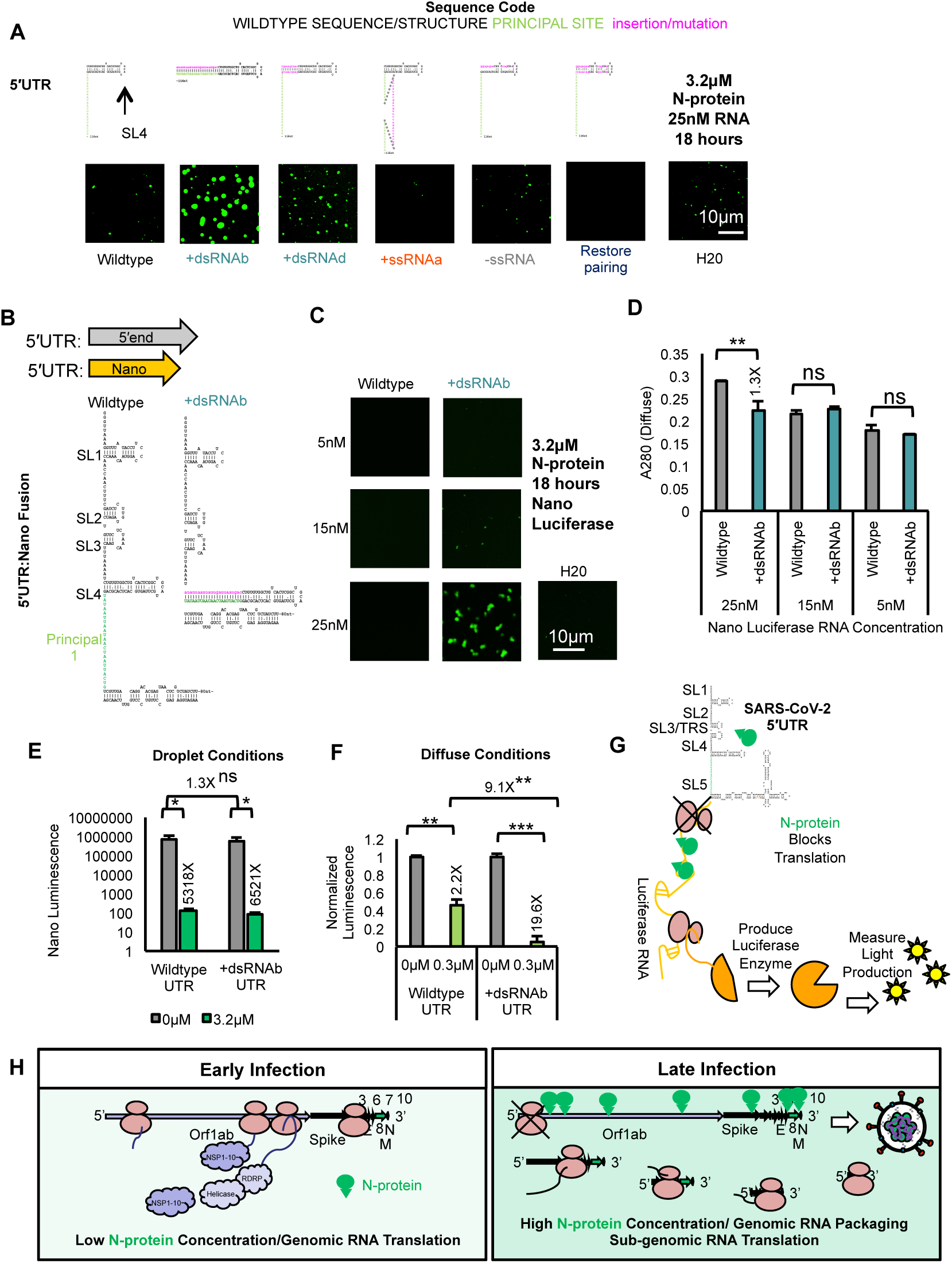
N-protein binding and LLPS represses translation. **(A)** Only +dsRNA (teal) mutants enhance LLPS in the context of the 5′UTR fragment. All other mutations do not significantly alter LLPS. 3.2uM N-protein (green) 25nM RNA 18 hours of incubation. H20 is water only control. **(B)** Design of luciferase fusion to the 5′UTR of SARS-CoV-2 constructs. Wildtype or a more structured 5′UTR (+dsRNAb) was fused to nano luciferase (orange). Arrows show the approximate length of nano luciferase as compared to 5′end 1-1000 RNA**. (C)** Only +dsRNAb UTR: Nano Luciferase undergoes LLPS at the highest tested RNA concentration (25nM/3.2μM N-protein (green)) **(D)** A280 absorbance of the remaining protein in the diffuse phase from Fig. 4C. Error bars mark standard deviation for the three replicates and * indicate significance students T test (** <0.01, ns not significant) with brackets showing comparison for the indicated statistical test. **(E)** In vitro translation assay results for nano luciferase wildtype or more structured fusion constructs. 20-minute incubation with 3.2μM N-protein prior to in vitro translation is sufficient to completely repress translation of nano luciferase. Error bars mark standard deviation for the three replicates and * indicate significance students T test (** <0.01, ns not significant) with brackets showing comparison for the indicated statistical test. **(F)** Presence of N-protein LLPS promoting RNA structures is associated with reduced translation in diffuse phase conditions. Normalized luminescence for nano luciferase constructs (no protein control fluorescent signal is set to 1). Nano luciferase +dsRNAb has a much greater reduction in normalized signal as compared to wildtype. **(G)** Model for N-protein mediated repression of translation via RNA affinity in the diffuse phase. Pink spheres are ribosomes. Green spheres are N-protein. Luciferase RNA (orange line) is translated to produce luciferase enzyme (orange Pac-man) to measure light production. **(H)** Model for N-protein mediated repression of translation via RNA affinity in infection. High affinity sites in the 5’UTR are preferentially occupied by N-protein in early infection to shut down orf1ab translation and switch to packaging. Late-stage infection translation occurs preferentially in sub-genomic RNA.

To ask if 5′UTR or +dsRNA UTR could differentially regulate translation in droplets we fused either the wildtype 5′UTR or a more structured mutant (+dsRNAb) to nano luciferase (**Fig. 4B**). To determine if 5′UTR structure affects LLPS for the fusions, we mixed 3.2μM N-protein with 25, 15, or 5 nM RNA. At the highest tested RNA concentration, 25nM, only the more structured mutant UTR resulted in LLPS (**Fig. 4C**). Similarly, only 25nM RNA condition had a statistically significant difference in A280 absorbance in the diffuse phase (**Fig. 4D**). Thus, consistent with results above, addition of dsRNA facilitates LLPS of nano luciferase fusion RNA. We next asked how LLPS conditions (3.2μM N-protein 25nM RNA) impact translation? To this end, we performed an in vitro translation assay +/-3.2μM N-protein. We observed that addition of 3.2μM N-protein almost completely blocked the translation of RNAs (**Fig. 4E**) Collectively, these results suggest that N-protein droplet conditions block translation.

We next asked if translation inhibition depended on LLPS or N-protein binding in the diffuse phase? To this end, we repeated our in vitro translation assay this time with 0.3μM of N-protein (**Fig. 4F** over 10-fold less N-protein than in **Fig. 4E**). In these conditions, translation of the wild-type UTR was moderately but significantly repressed by 0.3μM N-protein addition, and the translation of the +dsRNA UTR mutant was almost completely repressed translation (9.1X further reduction in translation compared to wildtype). This is consistent with phase behavior and N-protein affinity differences for these two RNAs (**Fig. 4C** and **D**). Collectively, these data suggest that droplet promoting conditions completely block translation, diffuse state-promoting conditions partially block translation. Translational block is dependent on relative N-protein affinity for the translating RNA (**Fig. 4G**).

We hypothesize that N-protein binding to the genome (particularly the 5′UTR) may act to halt protein translation and promote packaging in later stages on infection (**Fig. 4H**). In support of this idea, N-protein RNA is low at early stages of infection and gradually increases (via generation of sub-genomic N-protein RNA at late stages of infection (Kim et al., 2020). Thus, as time increases translatable genome should go down, thereby promoting a switch from translation to packaging in late-stage infection.

### RNA sequence/ structure may encode N-protein genome interactions to pattern RNP formation in virions

Given the key central role of N-protein in genome packaging, we next asked how what we have learned thus far about different types of N-protein/RNA interactions may impact packaging. Specifically, we postulated that the 5′end of the genome should have different patterning and/or affinity for N-protein binding sites than the genome center. We reason that this is because the non-structural genes in orf1ab need to be efficiently translated and LLPS of N-protein clearly can repress translation (**Fig. 4H**). Further, sequence property differences between the ends and the center of the genome are supported by sequence analysis (Iserman et al., 2020). Given the surfeit of in cell and cell free genome structural data for SARS-CoV-2 and related coronaviruses that is now available (Huston et al., 2021; Lan et al., 2021; Sun et al., 2021), as well as the published genome sequence data (NC_045512.2) we next asked how the two “dsRNA stickers” we identified as important for N-protein LLPS (RBD1 binding structured YYAAAY and RBD2 binding dsRNA) (**Fig. 5A**) are distributed throughout the genome?

**Fig. 5.**
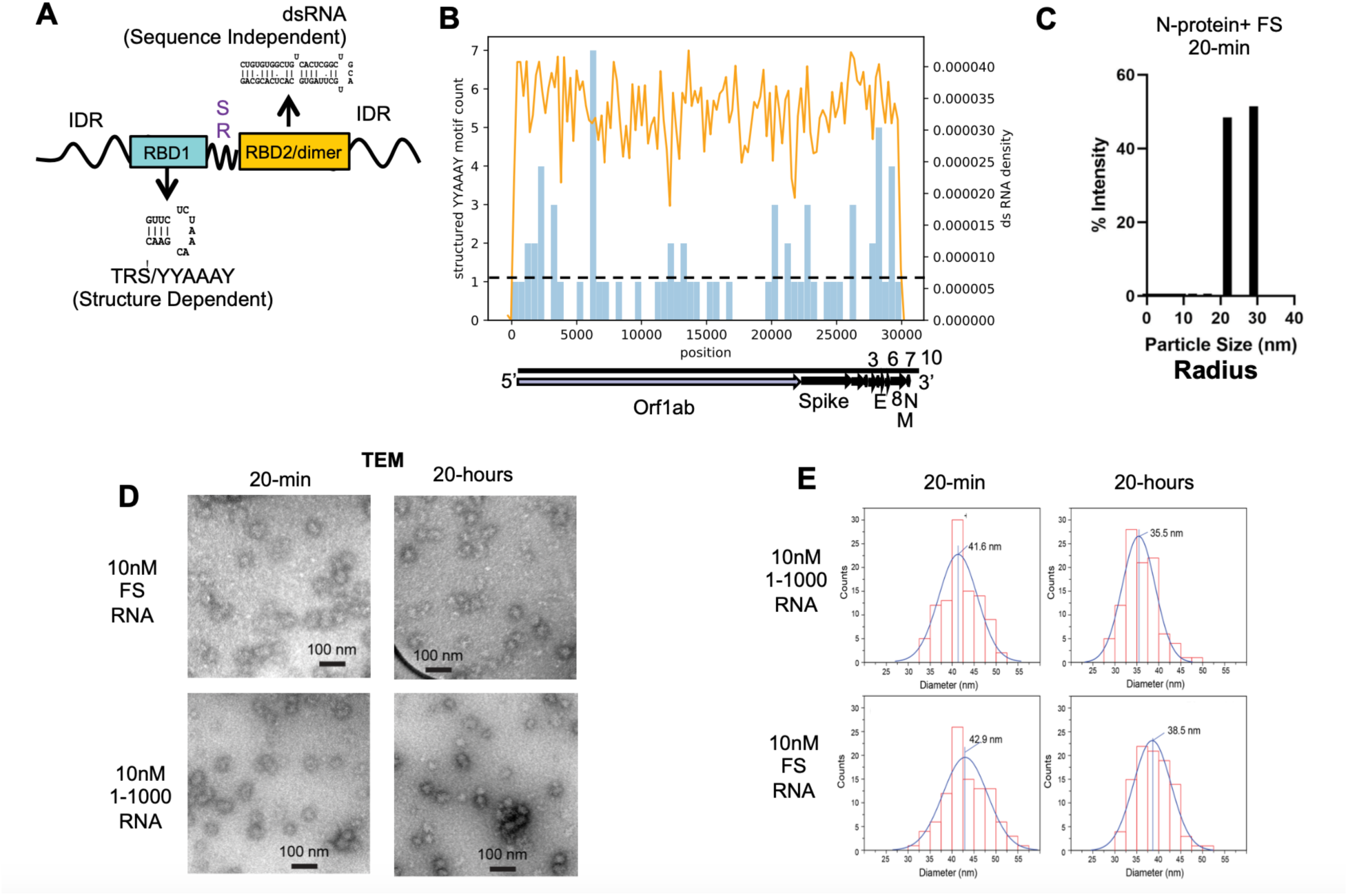
RNA sequence/ structure encodes N-protein genome interactions. **(A)** Model of RNA sequence preferences of SARS-CoV-2 N-protein RNA binding domains 1 (blue) and 2 (orange). RBD1 (teal box) binds TRS-like (YYAAAY) sequences in a structure dependent manner. RBD2/dimerization domain (orange box) binds dsRNA in a sequence independent manner. **(B)** Count of the structured YYAAAY motif (blue) and density of dsRNA (orange) across the SARS-CoV-2 genome. **(C)** Dynamic light scattering of 10nM FS RNA and 4μM protein. Following 20-minutes of incubation results in particles of 21.851 or 28.986nm radius (43.7-57.972nM in diameter). **(D)** Representative TEM images of small clusters which form from a mixture of 4μM N-protein and either 10nM FS or 10nM 1-1000 5′end when incubated for 20-minutes or 20-hours at room temperature. Scale bar is 100nM. **(E)** Quantification of small clusters as depicted in panel **5D**. for 1-1000 5′end, or FS. Clusters shrink by ∼15% following 20-hours of incubation

To determine the patterning of our two identified N-protein “stickers”, we obtained published in cell SARS-CoV-2 DMS-MaP based RNA structure models (Lan et al., 2021) to obtain locations of dsRNA. We also collected the density of YYAAAY from NC_045512.2 reference SARS-CoV-2 genome. Given our observation that structure of YYAAAY is important for binding (**Figure 2**) we restricted the YYAAAY motif to only those which overlap with structured RNA (80 structured YYAAAY motifs) (**Fig. 5B**). We observed that both features were relatively uniformly distributed across the genome with enrichment of structured YYAAAY motifs (above black dashed line) at the 5′ and 3′ ends of the genome, perhaps to promote genome circularization (Seim et al., 2021).

While the enrichment of LLPS-driving sequences at the 5′ and 3′ ends could suggest that LLPS may promote genome circularization (Iserman et al., 2020; Seim et al., 2021; Ziv et al., 2020) the final packaged genome is clearly not a single droplet. Based on high-resolution cryo-EM tomography, the genome of SARS CoV-2 is arranged inside virions in a so-called “birds-nest” arrangement with “eggs” made of RNP complexes that are ∼14-20nm (Klein et al., 2020; Yao et al., 2020). We previously observed that RNA derived from the center of the SARS-CoV-2 genome including RNA encoding the Frameshifting-region (FS) promoted N-protein solubilization at the microscopic level (Iserman et al., 2020). We reasoned that the solubilizing effect of FS RNA may be conferred by the formation of diffraction limited clusters that may be distinct from LLPS or are arrested from coarsening into macroscopic droplets. If indeed small RNP-scale particles form in this cell free system this would indicate that N-protein binding to RNA, as dictated by RNA sequence, was sufficient to condense RNA independent of cellular machinery.

To address if N-protein mediated condensation is sufficient to compact RNA to RNP-size assemblies’ cell free, we first asked what size particles form from FS RNA (1000nt in length)? We examined FS RNA as this RNA does not drive LLPS at 4μM N-protein 10-15nM RNA (**Fig. 3A**). To this end, we measured the particles formed from 10nM FS RNA and 4μM N-protein by dynamic light scattering. We chose 400X protein to RNA as this would be reminiscent of late-stage infection and packaging (Kim et al., 2020). We observed that following 20-minute incubation time at room temperature FS RNA forms homogenously sized clusters 43.7-57.9 nM in diameter (**Fig. 5C**) suggesting RNA cluster generation can occur cell free.

To directly visualize cluster formation a second way, we used TEM. Indeed, after 20-min of incubation a relatively monodispersed population of symmetric, circular assemblies form that are centered on 42.9 nm diameter (**Fig. 5D**). To assess if these formations were specific to FS RNA, we also examined N-protein 1-1000 RNA in conditions that do not support phase separation (room temperature). These formed similarly shaped and sized particles as the FS RNA (**Fig. 5D**). The assemblies formed with both RNAs are more than double the size of the reported RNP (∼14-20nM) diameter (Klein et al., 2020; Yao et al., 2020).

We wondered what caused the >2-fold size discrepancy between these RNP assemblies and the RNPs seen in virions? It is established that some droplets age into gel-like or glass-like states that can be associated with compaction (Jawerth et al., 2020), we therefore asked how the clusters change with time. Indeed, at the 20-hour time point smaller, more similar sized clusters for both RNAs were formed (**Fig. 5D**) indicating the clusters are shrinking by ∼15% over time, independent of RNA sequence (**Fig. 5E**). Some larger, rarer clusters were detected at 20-hours for both RNAs (**Fig. S5A and B**). Thus, N-protein and 1kB gRNA form monodispersed clusters cell free that compact over time. Both RNA and protein are required to form clusters (**Fig. S5C**). The similar size distribution of 5′end and FS fragments may result from the similar length (1kb) and overall affinity for RBD1 (the temperature insensitive RNA binding domain).Therefore, condensation differences between 5′end and FS require temperature-sensitive RBD2 interactions.

We postulated that FS interactions with N-protein may be heavily dependent on RBD1 rather than RBD2. In support of this hypothesis, the FS RNA contains 7 YYAAAY motifs 5 of which are structured and FS does not engage with RBD2 in a way that alters LLPS temperature (**Fig. 3A**). To confirm FS N-protein interactions are strongly RBD1 dependent, we performed RNP-map on FS with wildtype and Y109A mutant (RBD1 deficient) N-protein (**Fig. S5D and E**). We observed that the majority of the N-protein crosslinking peaks in FS were absent following incubation with Y109A mutation. Some Y109A-independent crosslinking was detected (purple boxes) and this tended to be adjacent to structured RNA. Thus, FS/N-protein interactions are primarily driven by RBD1 **(Fig. S5D and** E) whereas 5′end N-protein interactions are driven by both RBD1 and 2 **(Fig. S3A and B).** RBD1 binding site patterning conferred by structured YYAAAY motifs may be required for RNP-sized cluster generation.

## Discussion

In this paper, we elucidate the RNA sequence and structure preferences of SARS-CoV-2 N-protein to understand how these features lead to condensate properties relevant to viral processes in cells. We show that **1)** RBD1 prefers TRS-like sequences in an RNA structure dependent manner. **2)** RBD2 prefers dsRNA in a sequence-independent manner. **3**) RBD2 dsRNA interactions encode N-protein LCST behavior (**Fig. 6A**)**. 4)** RNA sequence/structure features specify N-protein interactions to regulate puncta formation rate, translation, and RNA cluster size cell free (RNP size) (**Fig. 6B**). To our knowledge, this is the first example of a role of dsRNA in encoding “stickers” and specific droplet properties.

**Fig. 6.**
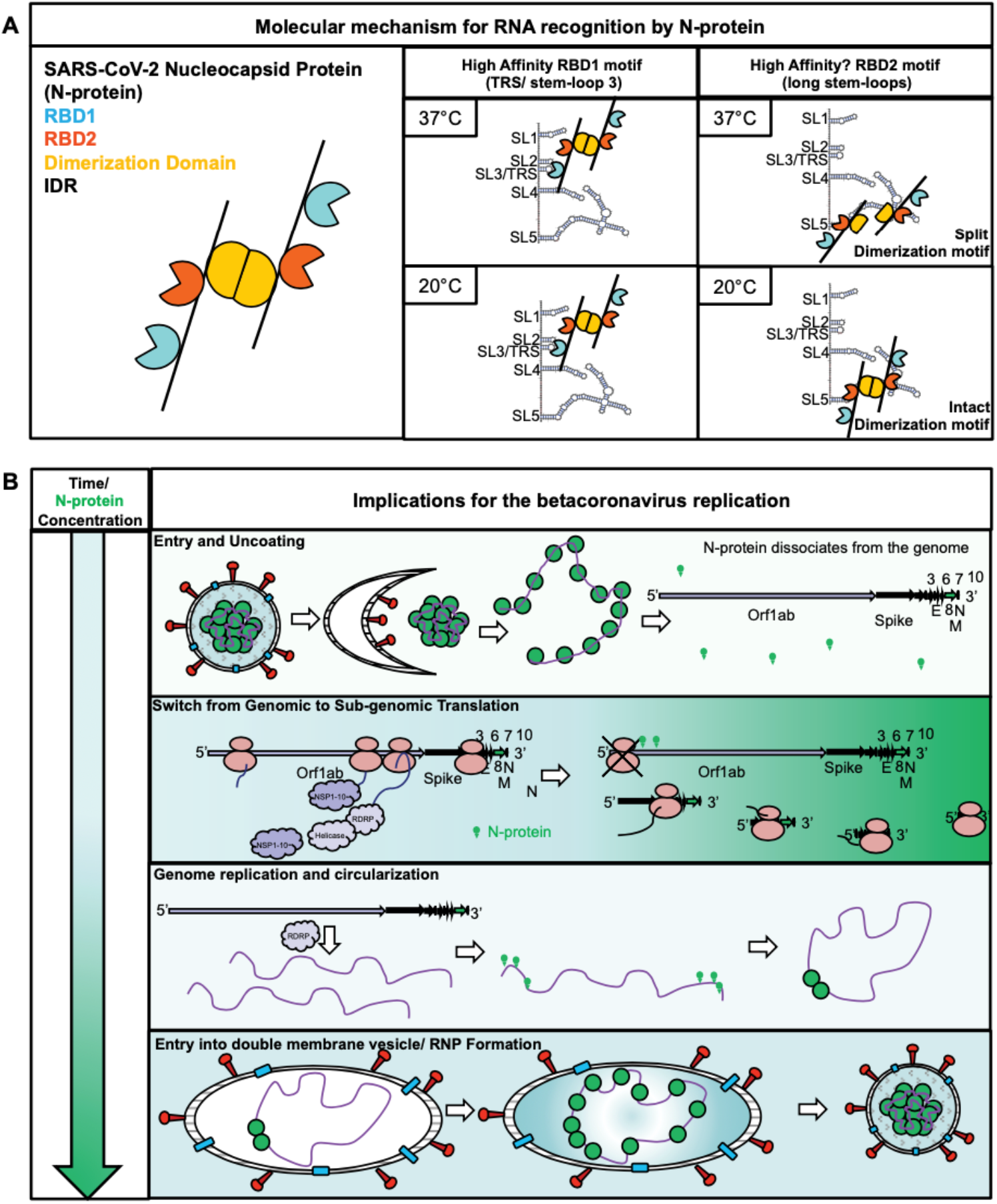
Model. **(A)** N-protein’s two RNA binding domains prefer two dsRNA dependent RNA stickers. RBD1 (teal) binds TRS stem-loop (and similar sequences) with high affinity in a temperature independent manner. RBD2 (dark orange) binds long stem-loops with temperature facilitating dimer (light orange) dissociation and LLPS. **(B)** Time dependent accumulation of N-protein specifies N-protein’s multiple roles in betacoronavirus by tuned patterned affinity for the two dsRNA dependent RNA stickers in **(A)**. High affinity sites (genome ends) are occupied preferentially early in infection when N-protein concentrations are low. Low affinity sites (genome center) are occupied late in infection when N-protein concentrations are high. Occupation of high affinity sites at genome ends promotes the switch from genome translation to circularization and ultimately packaging.

### N-protein accumulation regulates infection by tuned/patterned affinity for dsRNA

Our data (**Fig. 6B**) suggests a mechanism by which N-protein can perform multiple distinct functions over time in the same cytoplasm depending on N-protein concentration. Following viral entry, N-protein concentration is low. The low protein concentration allows for N-protein to dissociate from the condensed genome and for the initiation of translation. As infection progresses, N-protein’s (and other structural proteins) accumulation is driven by production from sub-genomic transcripts (Finkel et al., 2021; Kim et al., 2020). The accumulation of N-protein initiates a switch from translation to packaging, shutting down non-structural protein production while sparing the sub-genomic RNAs (which lack the most structured stem-loops 5 and on) (Kim et al., 2020; Sun et al., 2021). The structure of our RBD1 motif, the TRS, on the sub-genomic RNA is also predicted to regulate translation in a structure dependent manner with unstructured TRS present on the highly translated N-protein sub-genomic RNA (Finkel et al., 2021; Sun et al., 2021).The enrichment of high affinity N-protein binding sites at genome ends may allow for LLPS-mediated circularization to promote single genome packaging (Iserman et al., 2020; Seim et al., 2021). Finally, within double membrane vesicles, RNPs form as additional N-protein accumulates over time, with high concentration driving N-protein recruitment to low affinity sites in the genome center. The condensed genome ultimately matures into virions. We have identified unique dsRNA encoded “stickers” for N-protein conferred by the two RNA binding domains. The patterning and quality of the two N-protein dsRNA stickers can confer N-protein’s multiple functions through concentration dependent binding and LLPS. Thus, biochemical complexity needed for viral replication can be achieved with minimal components.

### RBD2 prefers dsRNA in a sequence independent manner

Addition of dsRNA, independent of sequence (**Fig. 1D-F, Fig. S1A**), resulted in more LLPS at all tested temperatures (**Fig. 3B**) and conditions (**Fig. S1C**). Reduction or addition of short ssRNA (comparable lengths to dsRNA mutants) sequences resulted in negligible enhancement of LLPS (**Fig. 1D-F, Fig. S1B**). Unpairing dsRNA generally reduced LLPS (**Fig. 1D-F**). There is likely an absolute length preference for RBD2 binding (Fig. S1A) which is consistent with 5′UTR stem-loop length altering experiments leading to viral plaque reduction (Raman and Brian, 2005; Yang et al., 2011). The lack of observed primary sequence specificity to N-protein RBD2 dsRNA binding **(Fig1. D-F)** may explain why previous stem-loop swap experiments, switching stem-loops from one betacoronavirus for another, generated functional virus (Guan et al., 2011; Kang et al., 2006a, 2006b). Although dsRNA length is important for RBD2 engagement, the specific sequence of the stem-loops is not critical (**Fig. 1D-F**) suggesting that N-protein may be able to engage with the entirety of the highly structured genome of SARS-CoV-2 (Huston et al., 2021; Lan et al., 2021; Sun et al., 2021). The lack of dsRNA sequence specificity is also suggested by the nature of the RBD2 motif, a disordered lysine rich IDR **(Fig. 3F, Fig. S4C),** which is unlikely to have primary RNA sequence specificity. Therefore, these data suggest that the sequence of the stem-loop does not matter for viral production and only minor differences in length are tolerated.

Importantly, our results suggest excessive differences in the length of the stem-loops appears to alter temperature encoding behavior with ∼20-24nt of dsRNA (present in SL5, SL12, and SL13 (**Fig. 1A**) encoding LLPS at 37°C and additional dsRNA (10nt+ - 80nt+) lowering the temperature to as low as 25°C (**Fig. 3B** and **C**). The absolute length of the stem-loop must be under a degree of selective pressure. Importantly, the most structured stem-loops (5, 12 and 13) are highly conserved (**Fig. 1B** (Yang and Leibowitz, 2015)) suggesting that stem-loop length mediated regulation is a universal feature for proper viral production, but subtle differences exist in the stem-loop length between individual viruses. This matches with experiments in MHV virus where altering the pairing of stem-loop 1 reduced the efficiency of viral production (Li et al., 2008). This suggests that stem-loop length may be co-evolving with N-protein RBD2/dimerization sequence, protein amount or both.

### RBD1 prefers TRS-like sequences in an RNA structure-dependent manner

RBD1 preferentially crosslinked adjacent to TRS and TRS-loop-like sequences (YYAAAY motifs) in the first 1000nt of the SARS-CoV-2 genome (Iserman et al., 2020). Improperly structured TRS loop sequences resulting from point mutation or deletion of the TRS stem-loop completely abrogated N-protein LLPS with 5′end 1-1000nt RNA. This is consistent with RBD1 mutant N-protein no longer undergoing LLPS with 5′end RNA (Iserman et al., 2020) and previous findings in MHV suggesting TRS is specifically recognized by N-protein RBD1 not RBD2 (Grossoehme et al., 2009). Unique to our findings of the TRS motif is the discovery that the secondary structure, in addition to the primary sequence, is important for SARS-CoV-2 N-protein LLPS and presumably binding. Our results suggest weak RBD1-mediated binding can occur at regions that are not the TRS stem-loop but have similar sequence and are structured (i.e., structured YYAAAY motifs).

### RBD2 dsRNA interactions encode N-protein LCST behavior

We postulate that the native, more structured stem-loops of the genome ends (i.e. SL5, 13 and 14 in the 5′end) are the most efficient binding sites for RBD2 (as evidenced by RBD1 independent crosslinking adjacent to these stem-loops (**Fig. S3A and B**) (Iserman et al., 2020) and this binding promotes LLPS at human body temperature (37°C). We predict that the binding to RBD2 in combination with physiological temperature (37°C) allows for the dissolution of the dimerization domain adjacent to RBD2 (**Fig. 4J**). Temperature is likely to facilitate the “unfolding” of the dimerization domain (the location of the fluorescent amino acids) as purified RBD2-dimerization domain undergoes a structural change at ∼50°C by differential scanning fluorometry (Zinzula et al., 2021). Of note, the temperature of dimerization unfolding determined by Zinzula et al for purified RBD2 dimerization domain alone is very close to our observed full-length N-protein only turbidity temperature (46°C **Fig. 3A**) suggesting temperature-dependent unfolding of this domain is critical to LCST behavior. The exposure of the hydrophobic core of the dimerization domain to the solution following temperature and RNA engagement may facilitate LLPS as hydrophobic regions tend to be insoluble.

The temperature at which dimerization occurs can be lowered via addition of dsRNA, potentially due to increasing the overall affinity of wildtype N-protein’s two RBD2 domains or by offering additional sites of interaction (at a greater distance apart) on the same stem-loop. In support of the latter possibility, the two RNA binding domains are arranged diagonal to each other, and dsRNA binding may force the dimer apart (**Fig. 3J**). Cryo-EM data of purified RBD-2/dimerization domain with ssRNA seems to agree with this hypothesis (Zinzula et al., 2021) with 7 base pairs of ssRNA spanning between two RBD2 motifs and a marked separation in the dimer region.

Our results also may explain why not all labs reporting N-protein LLPS have observed LCST behavior in N-protein RNA interactions as these results show that LCST behavior is specifically encoded in N-protein dsRNA interaction. N-protein dsRNA interaction is unlikely to be observed in reconstitution experiments conducted with less physiological, unstructured RNA (poly U for example). RBD2 also seems to regulate the dimerization domain of N-protein (Zhao et al., 2021a). RBD2 dimerization is highly dependent on the salt concentration with only physiological salt concentrations (150 +/-∼30) allowing for LCST behavior (Iserman et al., 2020; Lu et al., 2021; Zhao et al., 2021a). Lower salt results in an increase in N-protein dimerization domain interactions (Jack et al., 2020) which might increase the required total solution concentration of N-protein for phase separation, thus also increasing the temperature boundary of the LCST behavior. Others have not observed LCST behavior using nearly identical RNA sequences and N-protein preparation methods, but they were using much lower salt (80) (Lu et al., 2021). We conclude that because physiological levels of salt are more likely to be present in cells, LCST behavior of N-protein is relevant.

As RBD2 may recognize RNA through a complex interaction involving charge, disorder, and transient protein structure. It is highly likely that post-translational modifications (PTMs) play a huge role not only in RBD-2 binding RNA but also dimerization and LCST behavior. This may begin to explain why those labs which purify N-protein from mammalian cells did not observe LCST while those which purify N-protein from bacterial sources did (Iserman et al., 2020; Jack et al., 2020; Zhao et al., 2021a). As packaged N-protein is specifically free of post-translational modifications (Fung and Liu, 2018; Wu et al., 2009), we hold that the LCST behavior is likely still relevant for packaging with other N-protein compartments such as those regulating viral RNA transcription and translation being far more likely candidates to be regulated by PTMs (particularly those droplets that form outside double membrane vesicles associated with packaging). Future directions will explore how PTMs tune LLPS temperature to potentially sustain viral replication and viral RNA N-protein interactions during late-stage infection/fever temperatures.

Additionally, our results suggest the primary sequence of RBD2 dimerization region is critical for RNA binding and LCST behavior we would postulate that any mutation that arises and is selected for in these regions (in patient isolates or across species) would be particularly informative. Therefore, future directions will carefully characterize the sequence variation in this region.

### RNA sequence/structure features encode N-protein interactions to regulate RNA condensation

Distinct N-protein “dsRNA stickers” are distributed throughout the genome (**Fig. 5B**). This prompted us to hypothesize dsRNA sticker patterning could be relevant for packaging. Although N-protein clearly has tendencies to form macroscopic condensates in vitro and in cells, the packaged genome is instead packed into regularly-spaced RNPs which may be arrested in coarsening. Reconstituted N-protein mixed with 1000nt RNA fragments in physiological salt and pH was able to form clusters that were roughly 1.75X-2.5X the diameter of the RNP, the unit of packaging of the virion. This size difference suggests that either 1) the RNA content of the RNP is ∼500-1000nt (to give a 14-20nm RNP diameter) or 2) further, compaction occurs in the cells. We suggest the former possibility is more likely as there is a number range of RNPs (30-35 by cryo-EM suggesting each RNP must contain less then 1000nt (∼30kb genome/ ∼35 RNPs) and there is likely a flexible linker region composed of RNA depleted in N-protein (less electron dense) between each RNP to facilitate compaction. These data suggest that the information needed to condense the RNA genome is contained within the genomic RNA sequence.

Future directions will involve modeling of the SARS-CoV-2 dsRNA and structured TRS-loop-like sequences patterning across the genome to examine if indeed sequence element patterning is sufficient for RNP patterning (Choi et al., 2019; Seim et al., 2021). Of note, the length of viral RNA fragments tested in this work (0.5 and 1kB) is highly relevant for this consideration as each RNP/egg is likely to contain <1kB. We and others have observed that longer RNAs (Jack et al., 2020) including RNA purified from infected cells containing SARS-CoV-2 genome (Iserman et al., 2020) results in a “string of pearls” type droplets rather than rounded droplets further suggesting that the formation of RNPs/eggs is recapitulated cell free. Thus, the fragments tested here are short enough to encode single RNP/egg like features but long enough to have sequence and structural complexity to allow for observable regional differences in LLPS.

### Considerations for other RNA-based LLPS

Notably, increasing RNA order, rather than disorder, through additional RNA structure drives N-protein LLPS. These results contrast with those observed by Mayr lab where increased RNA disordered single stranded regions promoted intramolecular association and LLPS (Ma et al., 2021). This discrepancy is likely due to differences in the proteins, specifically the preference of both of N-protein’s RNA binding domains for the highly-structured RNA genome of SARS-CoV-2 (dsRNA stickers).

Our work suggests that reconstitution experiments of phase separating proteins with similar dsRNA preferences must be carried out with physiological RNA targets to capture biological behavior. DsRNA protein interactions are not captured with poly A or U. In short, RNA sequence and structure profoundly influence LLPS behavior. Finally, this work shows the complexity of the RNA-protein code in determining the kinetics, emergent properties and temperatures at which biomolecular condensates can form. We predict that this is the tip of the iceberg in terms of unraveling the information provided by RNA sequence to specify the form and function of condensates.

## Acknowledgements

We thank Rick Young, Phil Sharp, Alex Holehouse, Kathleen Hall, Andrea Sorrano, Ahmet Yildez and their lab members for sharing data and discussions. Alain Laederach for initial discussion on genomic sequence, Chase Weidmann for initial discussions planning RNP-MaP experiments. We thank Benjamin Stormo for providing purified Whi3 protein. A.S.G., C.I. and C. A. R. were supported by NIH R01GM081506 and an HHMI faculty Scholar Award, C.A.R. was supported by NIH T32 CA 9156-43, F32GM136164 and L’OREAL USA for Women in Science Fellowship. This work was supported by a FastGrant. I. S. and A. S. G. were supported by Air Force Office of Scientific Research (grant FA9550-20-1-0241). M.A.B. was supported by a Ruth L. Kirschstein Postdoctoral Fellowship (F32 GM128330). Work by M.L. was supported by the Intramural Research Program of the National Institute of Diabetes and Digestive and Kidney Diseases, National Institutes of Health. K. M. W. was supported by NIH R35 GM122532.

## Contributions

C.A.R. and A.S.G. conceptualized the project, designed experiments, prepared figures, drafted and edited the manuscript. C.A.R. and S.A.W. performed experiments and analyzed data; Y.D. and M.L. performed experiments, prepared figure and analyzed data M.A.B. designed and performed experiments and computational analyses, analyzed data, prepared figures, and drafted the manuscript. K.M.W. designed experiments, analyzed data, and drafted the manuscript. G.A.M. and I.S. designed and performed experiments, analyzed data, edited the manuscript and performed computational analyses. R.S. analyzed data. O.G., L.Y., A.C., J.E, and C.I. designed experiments and drafted the manuscript.

## Limitations

This study addresses mechanisms of phase separation of components of the SARS-CoV-2 virus. However, because the work involved reconstitution experiments from purified components and expression of viral proteins in mammalian cells rather than in an actual infection it is still unclear what step(s) in the viral replication cycle may utilize the mechanisms described. The work is also is missing some factors such as M-protein and lipid membranes that may change the physical chemistry of the phase separation in an infection.

## Supplemental Figures/Figure Legends

**Fig. S1.**
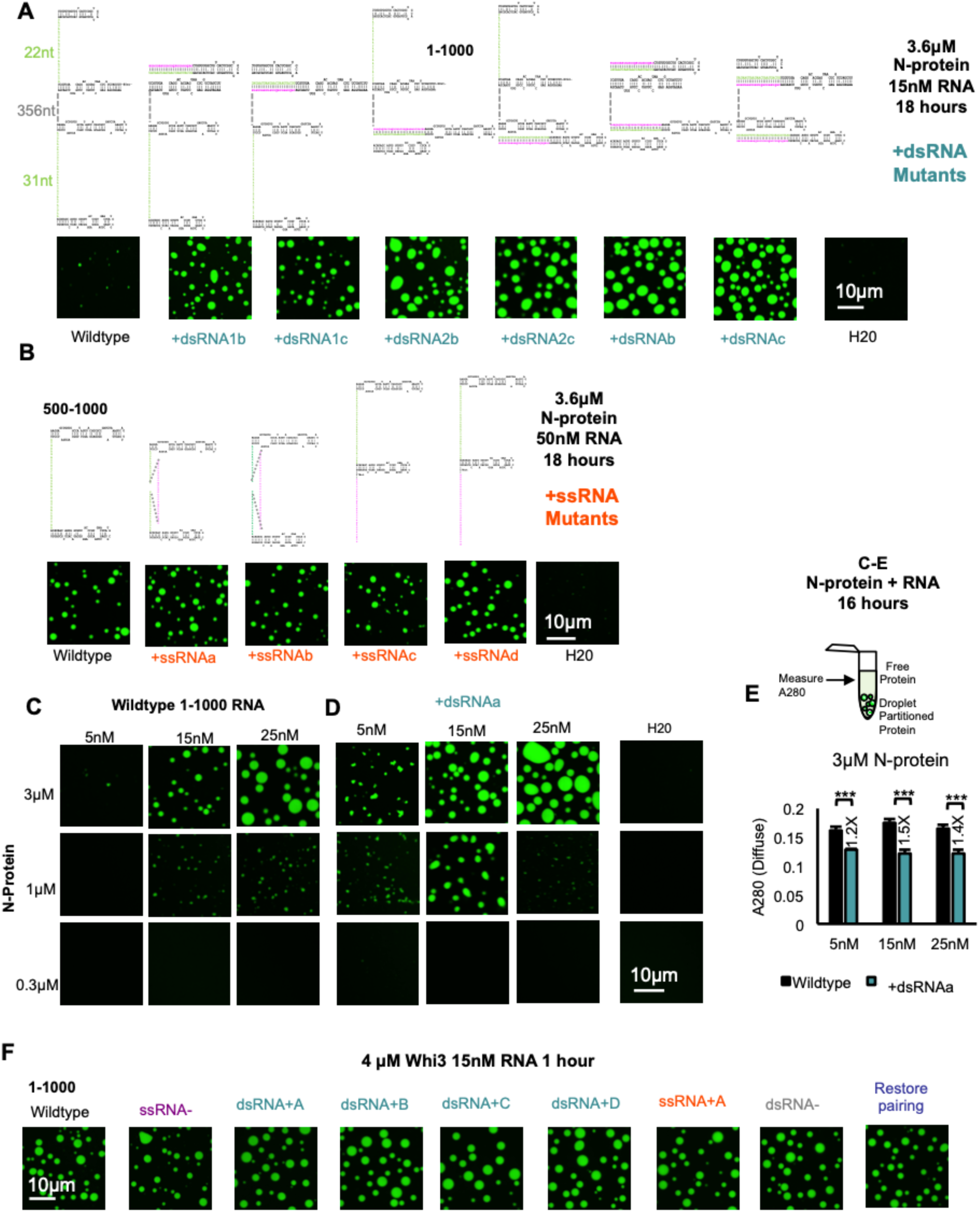
dsRNA-driven LLPS is independent of RBD1. **(A)** At 3.6μM N-protein (green) and 15nM RNA following 18 hours of incubation, mutations in the context of 1-1000nt which increase the dsRNA (+dsRNA teal) of principal site 1 only, 2 only, or 1, and 2 lead to enhanced LLPS in comparison to wildtype regardless of whether the anti-sense sequence to the principal site was inserted on the 5′ or 3′ side. Enhancement of LLPS may be length dependent. H20 in indicates equivalent volume of added water only control. **(B)** Location or sequence of ssRNA (+ssRNA orange) insertion leads to equally negligible levels of LLPS enhancement. 3.6μM N-protein, 50nM RNA 18hours of incubation in the 500-1000 sequence context. (**C-E)** Phase diagrams for equivalent RNA and N-protein concentrations (3μM, 1μM, 0.3μM) for 1-1000 wildtype, +dsRNAa for 5nM, 15nM, or 25nM RNA. At 3μM, +dsRNAa leads to more LLPS relative to Wild-type 1-1000 for all tested RNA concentrations. At 1μM +dsRNAa shifts the phase boundary to the left relative to wildtype. 0.3μM N-protein does not drive LLPS for any sequence at any tested RNA concentration. **(E)** A280 absorbance of for 3μM N-protein concentration for panels **S1C, D.** For all tested RNA concentrations relative to wildtype, +dsRNAa has less protein in solution following 16 hours incubation as measured by A280 nanodrop. Error bars mark standard deviation for the three replicates and * indicate significance students T test (*** p<0.001, ** p<0.01, ns not significant) with brackets showing comparison for the indicated statistical test. **(G)** 1-1000 RNA and its mutants (purple -ssRNA, teal +dsRNA, oranges -ssRNA, -dsRNA gray, Restore pairing blue) lead to equivalent LLPS of Whi3 protein (green). For all images scale bar indicates 10μm all experiments show representative images from at least 3 replicates.

**Fig. S2.**
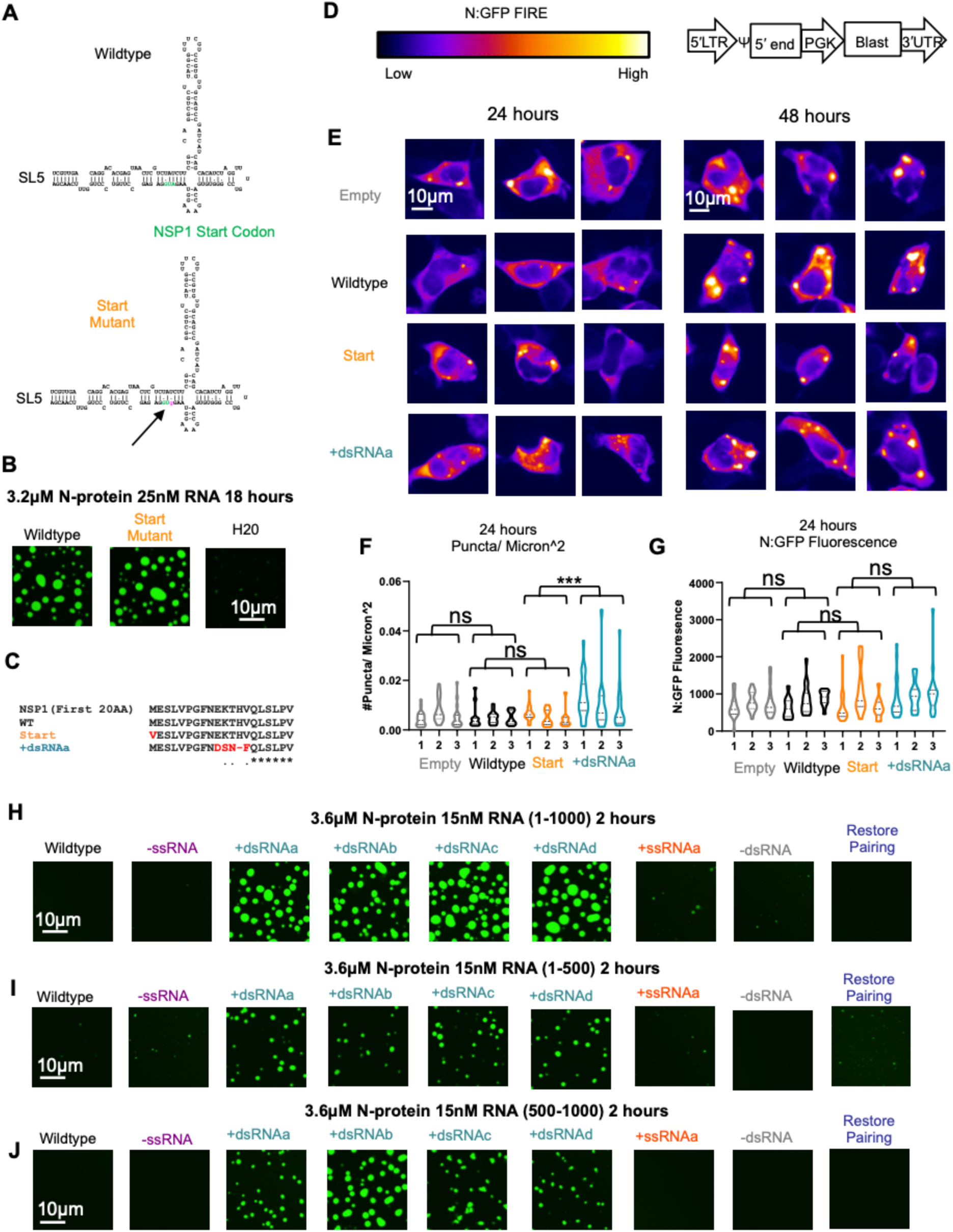
RNA structure mutants accelerate droplet formation cell free and in cells. **(A)** Structure model of SL5 for SARS-CoV-2 with the location of the start codon of non-structural protein 1 (NSP1) in orange text. AU base pair of the start codon is replaced with a GU wobble pair (ATG → gTG) to eliminate NSP1 translation while sparing RNA structure. **(B)** Wildtype and Start Mutant RNA result in similar levels of LLPS cell free. 3.2μM N-protein (green) and 25nM RNA following 2.5 hours of incubation. **(C)** NSP1 protein sequence of mutants tested in **S2E-G**. destroy NSP1 production. **(D)** N: GFP protein signal (FIRE blue low signal white high signal) key and 5′end overexpression plasmids design. **(E)** Representative HEK293T cells co-transfected with N: GFP and the indicated 5′end fragment at 24 hours (left 3 panels) or 48 hours (right 3 panels). +dsRNAa mutant produces more puncta at 24 hours (4-5 per cell) compared to wildtype, start, or empty (2-3 per cell). Difference is reduced at 48 hours (4-5 puncta in all three 5′end containing cells). **(F)** Quantification of the number of droplets per Micron^2 at 24 hours. +dsRNAa produces significantly more puncta per unit area then the Start mutant. * indicate significance students T test (*** p<0.001, ns not significant) with brackets showing comparison for the indicated statistical test. **(G)** Quantification of the mean intensity of N: GFP signal at 24 hours. Analyzed cells have similar GFP signal distribution. No comparisons are significant (ns). **(H)** Addition of dsRNA (teal) (+dsRNAa-d) enhances N-protein LLPS. Representative images from the two-hour incubation timepoint for panels **1F**. N-protein signal is shown in green. **(I and J)** Representative images from the two-hour incubation timepoint for panels Figure 1D (1-500) and **1E** (500-1000). For all images scale bar indicates 10μm.

**Fig. S3.**
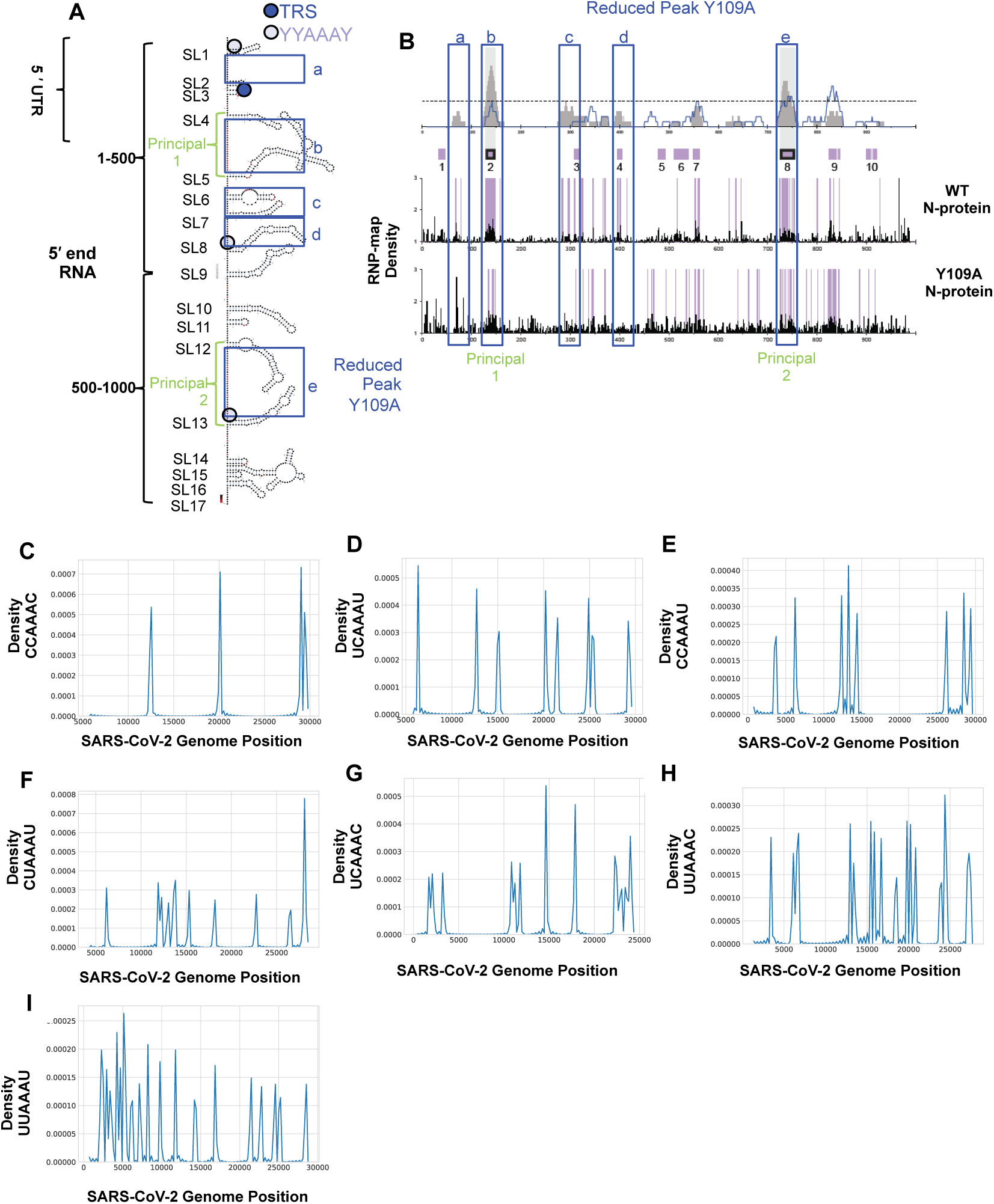
RBD1 dependent crosslinking adjacent to YYAAAY motif; distributed throughout the genome. **(A)** Structure of the 5′end 1-1000 RNA adapted from Iserman et al. (Iserman et al., 2020). Blue squares indicate the location of reduced N-protein crosslinking following Y109A mutation which destroys RBD1 (see **B**). Dark Blue and light blue circles indicate locations of the perfect TRS loop sequence (CUAAAC dark blue) verses (YYAAAY light blue) showing that peak reduction is often adjacent to TRS or TRS-like motifs. **(B)** RNP-map signal for wildtype N-protein and Y109A mutant N-protein (which is predicted to destroy RBD1. Blue boxes (lower case letters) are equivalent between **S3A** and **B**. Note that crosslinking of N-protein to RNA appears to occur more readily at single-stranded than double-stranded nts, despites N-protein’s ability to bind both double- and single-stranded nts. (see Figure 1). Green text indicates location of principal sites. **(C-I)** Density of TRS-Loop-like motif (YYAAAY) across the genome other than CUAAAC (Fig 2C **and D).** Y axis is the relative density of the indicated sequence across the SARS-CoV-2 genome.

**Fig. S4.**
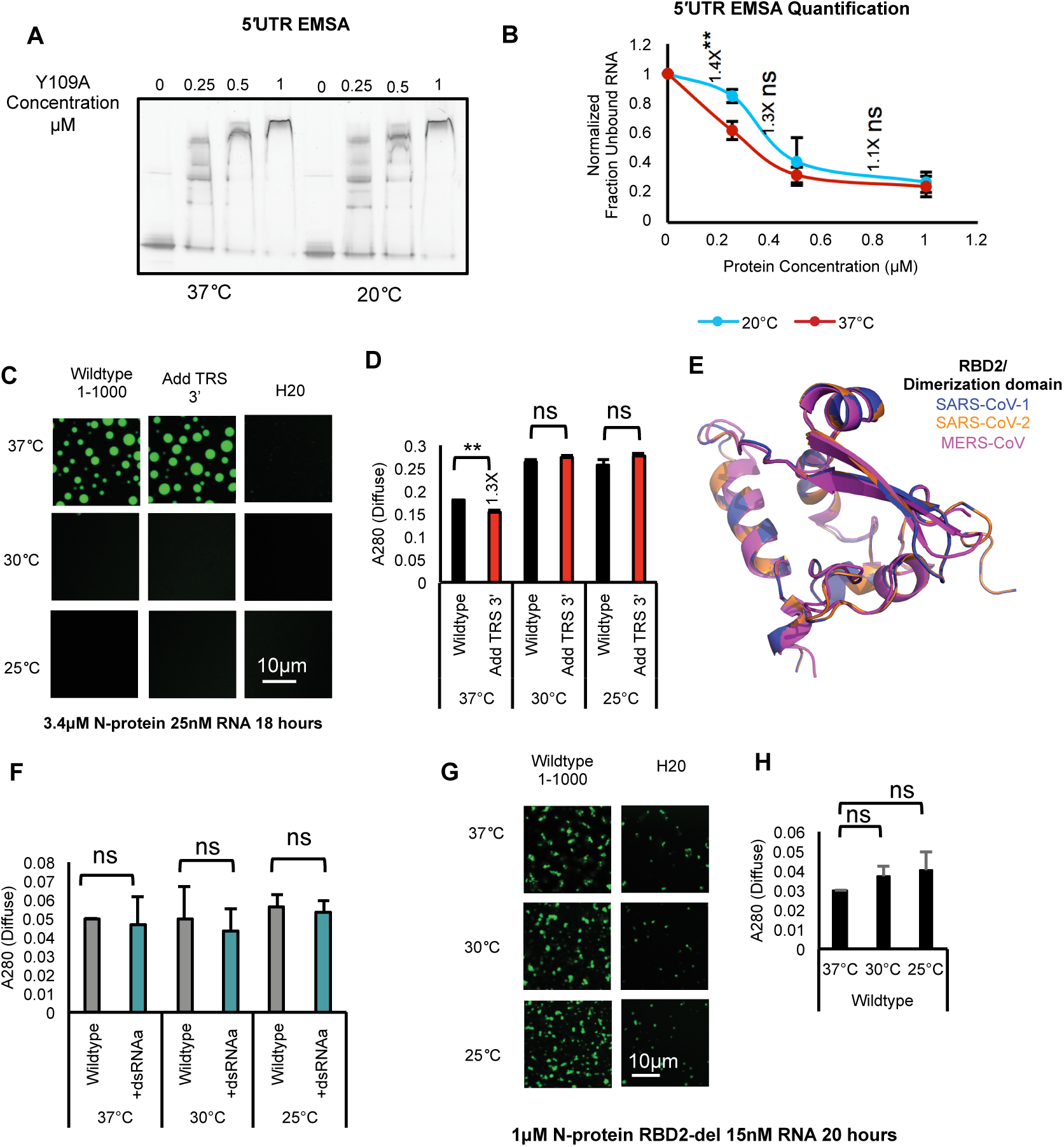
RNA sequence and structure encodes N-protein LCST Behavior via RBD2. **(A)** Representative EMSA for Y109A mutant N-protein and 5′UTR RNA at 37 and 25°C. **(B)** EMSA Quantification; 3 replicates. Less unbound 5′UTR RNA at 37°C incubation then 25°C incubation. Y axis is the normalized fraction unbound protein (0μM protein condition set to 1). X axis tested Y109A protein concentration. 37°C red line, 25°C, blue line error bars reflect the standard deviation. **(C)** Addition of a second TRS sequence structure motif (RBD1 binding site) does not alter the LCST behavior of N-protein. Wildtype 1-1000 5′end RNA or Add TRS 3′ mutant was incubated at 37°C, 30°C or 25°C for a period of 18 hours (N-protein signal shown in green). **(D)** A280 measurement of remaining N-protein in the solution for **Fig. S4A**. Only 37°C shows a difference in signal consistent with previous results (Fig 3B). (**E**) Alignment of the predicted structure for SARS-CoV-2 (orange ribbon), the crystal structure of MERS-CoV (magenta ribbon), to the crystal structure of SARS-CoV-1 (blue ribbon) RBD2 dimerization domain. Proteins may adopt similar folds. **(F)** A280 measurement of remaining N-protein in the solution for Fig. 3G. **(G)** Repeat of LCST experiment with wildtype RNA at lower N-protein (1μM) and RNA (15nM) concentration. H20 alone results in less LLPS indicative for some RNA dependence for N-protein RBD-2 Del. **(H**) A280 measurements for (**Fig. S4G**) At 37°C, 30°C and 25°C wildtype RNA has not significantly (ns) different A280 measurements indicative of similar amounts of protein in solution and identical amounts of LLPS. For all A280 measurements, error bars mark standard deviation for the three replicates and brackets indicate the comparison for students T test (** p<0.01,ns not significant). For all images scale bar indicates 10μm. All experiments show representative images from at least 3 replicates.

**Fig. S5.**
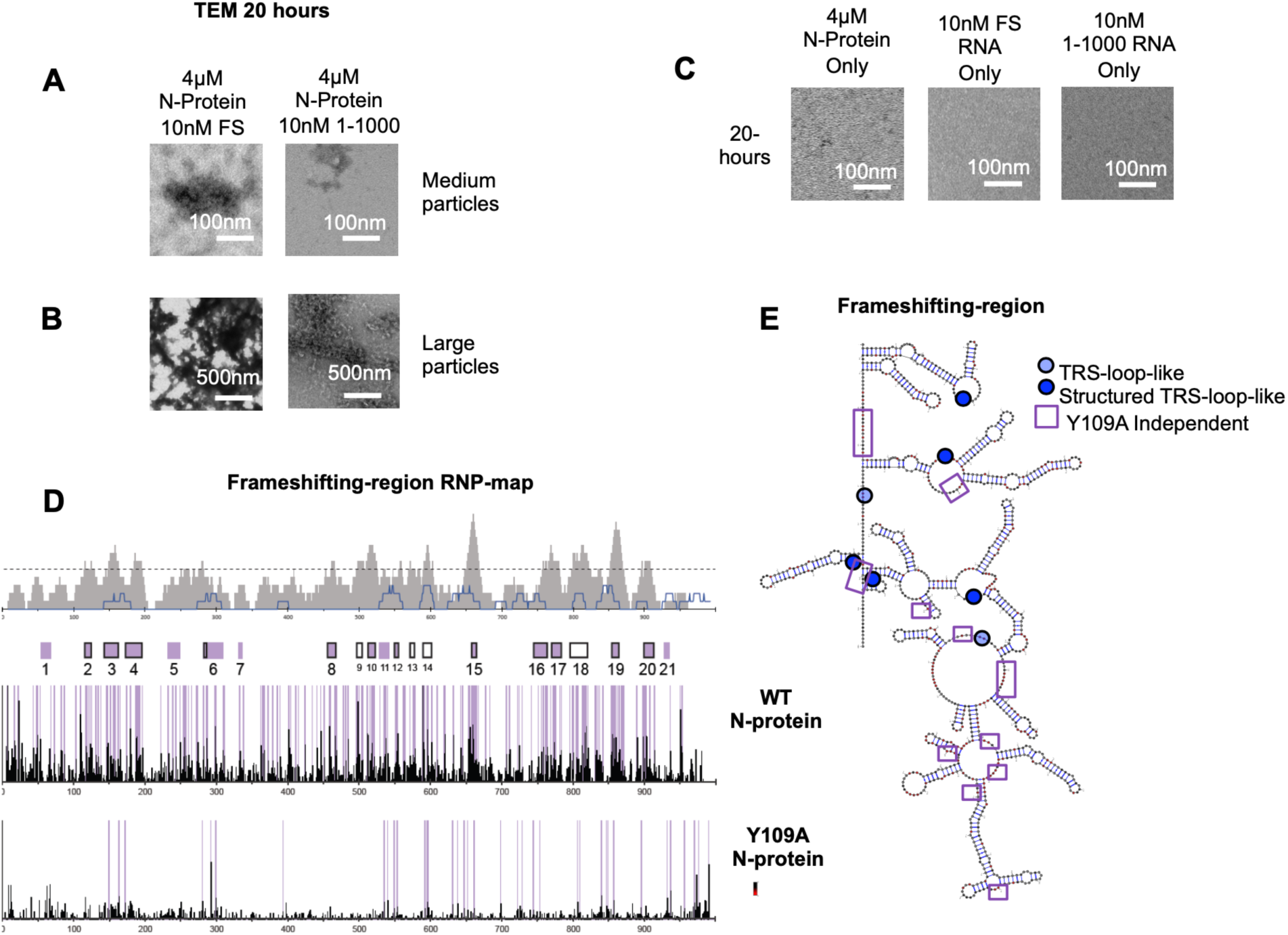
RNA sequence/ structure encodes N-protein genome interactions. **(A and B)** Representative TEM images from the 20-hour timepoint depicting medium sized **(A)** and large sized **(B)** clusters of N-protein and either 10nM frameshift left panels or 1-1000 5′end RNA. Larger clusters were not detected at the 20-minute timepoint. **(A)** Scale bar is 100nm. **(B)** Scale bar is 500nm. **(C)** RNA or N-protein alone does not form particles following 20-hours of incubation at room temperature. Representative TEM images of 4μM N-protein, 10nM FS RNA, or 10nM 1-1000 5′end incubated for 20hours at room temperature. Scale bar is 100nM. **(D and E)** RNP-map data for FS RNA with either wildtype or Y109A mutant N-protein. Majority of crosslinking is lost following RBD1 mutation. FS contains structured (dark blue circle) and unstructured (light blue circle)/ TRS-loop-like sequences. Peaks retained following Y109A mutation (purple squares) are in the vicinity of structured RNA.

## Methods

### Protein Production

#### Recombinant Protein Expression and Purification

For protein purification, full-length N-protein was tagged with an N-terminal 6-Histidine tag (pET30b-6xHis-TEV-Nucleocapsid, N-Y109A, and N-RBD2-Del) were expressed in BL21 E. coli (New England Biolabs). All steps of the purification after growth of bacteria were performed at 4°C. Cells were lysed in lysis buffer (1.5M NaCl, 20 mM Phosphate buffer pH 7.5, 20 mM Imidazole, 10mg/mL lysozyme, 1 tablet of Roche EDTA-free protease inhibitor cocktail Millipore Sigma 11873580001) and via sonication. The lysate was then clarified via centrifugation (SS34 rotor, 20,000 rpm 30 minutes) and the supernatant was incubated and passed over a HisPurTM Cobalt Resin (ThermoFisher Scientific 89965) in gravity columns. The resin was then washed with 4X 10 CV wash buffer (1.5M NaCl, 20 mM Phosphate buffer pH 7.5, 20 mM Imidazole) and protein was eluted with 4 CV Elution buffer (0.25 M NaCl, 20 mM Phosphate buffer pH 7.5, 200 mM Imidazole). The eluate was then dialyzed into fresh storage buffer (0.25 M NaCl, 20 mM Phosphate buffer) and aliquots of protein were flash frozen and stored at -80 °C. Protein was checked for purity by running an SDS-PAGE gel followed by Coomassie staining as well as checking the level of RNA contamination via Nanodrop and through running of a native agarose RNA gel. All experiments were performed with His-tagged N-protein. Whi3 was purified according to our established protocols (Langdon et al., 2018; Zhang et al., 2015).

### DNA sequences of N-protein constructs are as follows

### Wildtype

ATGCACCATCATCATCATCATTCTTCTGGTGAAAACCTGTATTTTCAGGGCGTCGACATGTCTGATAATGGACCCCAAAATCAGCGAAATGCACCCCGCATTACGTTTGGTGGACCCTCAGATTCAACTGGCAGTAACCAGAATGGAGAACGCAGTGGGGCGCGATCAAAACAACGTCGGCCCCAAGGTTTACCCAATAATACTGCGTCTTGGTTCACCGCTCTCACTCAACATGGCAAGGAAGACCTTAAATTCCCTCGAGGACAAGGCGTTCCAATTAACACCAATAGCAGTCCAGATGACCAAATTGGCTACTACCGAAGAGCTACCAGACGAATTCGTGGTGGTGACGGTAAAATGAAAGATCTCAGTCCAAGATGGTATTTCTACTACCTAGGAACTGGGCCAGAAGCTGGACTTCCCTATGGTGCTAACAAAGACGGCATCATATGGGTTGCAACTGAGGGAGCCTTGAATACACCAAAAGATCACATTGGCACCCGCAATCCTGCTAACAATGCTGCAATCGTGCTACAACTTCCTCAAGGAACAACATTGCCAAAAGGCTTCTACGCAGAAGGGAGCAGAGGCGGCAGTCAAGCCTCTTCTCGTTCCTCATCACGTAGTCGCAACAGTTCAAGAAATTCAACTCCAGGCAGCAGTAGGGGAACTTCTCCTGCTAGAATGGCTGGCAATGGCGGTGATGCTGCTCTTGCTTTGCTGCTGCTTGACAGATTGAACCAGCTTGAGAGCAAAATGTCTGGTAAAGGCCAACAACAACAAGGCCAAACTGTCACTAAGAAATCTGCTGCTGAGGCTTCTAAGAAGCCTCGGCAAAAACGTACTGCCACTAAAGCATACAATGTAACACAAGCTTTCGGCAGACGTGGTCCAGAACAAACCCAAGGAAATTTTGGGGACCAGGAACTAATCAGACAAGGAACTGATTACAAACATTGGCCGCAAATTGCACAATTTGCCCCCAGCGCTTCAGCGTTCTTCGGAATGTCGCGCATTGGCATGGAAGTCACACCTTCGGGAACGTGGTTGACCTACACAGGTGCCATCAAATTGGATGACAAAGATCCAAATTTCAAAGATCAAGTCATTTTGCTGAATAAGCATATTGACGCATACAAAACATTCCCACCAACAGAGCCTAAAAAGGACAAAAAGAAGAAGGCTGATGAAACTCAAGCCTTACCGCAGAGACAGAAGAAACAGCAAACTGTGACTCTTCTTCCTGCTGCAGATTTGGATGATTTCTCCAAACAATTGCAACAATCCATGAGCAGTGCTGACTCAACTCAGGCCTAA

### Y109A

ATGCACCATCATCATCATCATTCTTCTGGTGAAAACCTGTATTTTCAGGGCGTCGACATGTCTGATAATGGACCCCAAAATCAGCGAAATGCACCCCGCATTACGTTTGGTGGACCCTCAGATTCAACTGGCAGTAACCAGAATGGAGAACGCAGTGGGGCGCGATCAAAACAACGTCGGCCCCAAGGTTTACCCAATAATACTGCGTCTTGGTTCACCGCTCTCACTCAACATGGCAAGGAAGACCTTAAATTCCCTCGAGGACAAGGCGTTCCAATTAACACCAATAGCAGTCCAGATGACCAAATTGGCTACTACCGAAGAGCTACCAGACGAATTCGTGGTGGTGACGGTAAAATGAAAGATCTCAGTCCAAGATGGGCATTCTACTACCTAGGAACTGGGCCAGAAGCTGGACTTCCCTATGGTGCTAACAAAGACGGCATCATATGGGTTGCAACTGAGGGAGCCTTGAATACACCAAAAGATCACATTGGCACCCGCAATCCTGCTAACAATGCTGCAATCGTGCTACAACTTCCTCAAGGAACAACATTGCCAAAAGGCTTCTACGCAGAAGGGAGCAGAGGCGGCAGTCAAGCCTCTTCTCGTTCCTCATCACGTAGTCGCAACAGTTCAAGAAATTCAACTCCAGGCAGCAGTAGGGGAACTTCTCCTGCTAGAATGGCTGGCAATGGCGGTGATGCTGCTCTTGCTTTGCTGCTGCTTGACAGATTGAACCAGCTTGAGAGCAAAATGTCTGGTAAAGGCCAACAACAACAAGGCCAAACTGTCACTAAGAAATCTGCTGCTGAGGCTTCTAAGAAGCCTCGGCAAAAACGTACTGCCACTAAAGCATACAATGTAACACAAGCTTTCGGCAGACGTGGTCCAGAACAAACCCAAGGAAATTTTGGGGACCAGGAACTAATCAGACAAGGAACTGATTACAAACATTGGCCGCAAATTGCACAATTTGCCCCCAGCGCTTCAGCGTTCTTCGGAATGTCGCGCATTGGCATGGAAGTCACACCTTCGGGAACGTGGTTGACCTACACAGGTGCCATCAAATTGGATGACAAAGATCCAAATTTCAAAGATCAAGTCATTTTGCTGAATAAGCATATTGACGCATACAAAACATTCCCACCAACAGAGCCTAAAAAGGACAAAAAGAAGAAGGCTGATGAAACTCAAGCCTTACCGCAGAGACAGAAGAAACAGCAAACTGTGACTCTTCTTCCTGCTGCAGATTTGGATGATTTCTCCAAACAATTGCAACAATCCATGAGCAGTGCTGACTCAACTCAGGCCTAA

### RBD2-Del

ATGCACCATCATCATCATCATTCTTCTGGTGAAAACCTGTATTTTCAGGGCGTCGACATGTCTGATAATGGACCCCAAAATCAGCGAAATGCACCCCGCATTACGTTTGGTGGACCCTCAGATTCAACTGGCAGTAACCAGAATGGAGAACGCAGTGGGGCGCGATCAAAACAACGTCGGCCCCAAGGTTTACCCAATAATACTGCGTCTTGGTTCACCGCTCTCACTCAACATGGCAAGGAAGACCTTAAATTCCCTCGAGGACAAGGCGTTCCAATTAACACCAATAGCAGTCCAGATGACCAAATTGGCTACTACCGAAGAGCTACCAGACGAATTCGTGGTGGTGACGGTAAAATGAAAGATCTCAGTCCAAGATGGTATTTCTACTACCTAGGAACTGGGCCAGAAGCTGGACTTCCCTATGGTGCTAACAAAGACGGCATCATATGGGTTGCAACTGAGGGAGCCTTGAATACACCAAAAGATCACATTGGCACCCGCAATCCTGCTAACAATGCTGCAATCGTGCTACAACTTCCTCAAGGAACAACATTGCCAAAAGGCTTCTACGCAGAAGGGAGCAGAGGCGGCAGTCAAGCCTCTTCTCGTTCCTCATCACGTAGTCGCAACAGTTCAAGAAATTCAACTCCAGGCAGCAGTAGGGGAACTTCTCCTGCTAGAATGGCTGGCAATGGCGGTGATGCTGCTCTTGCTTTGCTGCTGCTTGACAGATTGAACCAGCTTGAGAGCAAAATGTCTGGTAAAGGCCAACAACAACAAGGCCAAACTGTCACTGGTCCAGAACAAACCCAAGGAAATTTTGGGGACCAGGAACTAATCAGACAAGGAACTGATTACAAACATTGGCCGCAAATTGCACAATTTGCCCCCAGCGCTTCAGCGTTCTTCGGAATGTCGCGCATTGGCATGGAAGTCACACCTTCGGGAACGTGGTTGACCTACACAGGTGCCATCAAATTGGATGACAAAGATCCAAATTTCAAAGATCAAGTCATTTTGCTGAATAAGCATATTGACGCATACAAAACATTCCCACCAACAGAGCCTAAAAAGGACAAAAAGAAGAAGGCTGATGAAACTCAAGCCTTACCGCAGAGACAGAAGAAACAGCAAACTGTGACTCTTCTTCCTGCTGCAGATTTGGATGATTTCTCCAAACAATTGCAACAATCCATGAGCAGTGCTGACTCAACTCAGGCCTAA

#### Dyeing of N-protein

N-protein was dyed by adding (3:1) Atto 488 NHS ester (Millipore Sigma 41698) to purified protein and incubating mix at 4°C for 1 hour with rocking. Unbound dye was removed by overnight dialysis into protein storage buffer. For LLPS percent of dyed protein was adjusted to 10% of total by dilution with undyed protein.

#### RNA Template Design/Production

Template predicted structure was designed using Vienna fold (http://rna.tbi.univie.ac.at). Sequences were generated via site directed mutagenesis using overlapping oligos (IDT). DNA sequences of tested RNA fragments are as follows.

### 1-500 Wildtype

GGGTTAAAGGTTTATACCTTCCCAGGTAACAAACCAACCAACTTTCGATCTCTTGTAGATCTGTTCTCTAAACGAACTTTAAAATCTGTGTGGCTGTCACTCGGCTGCATGCTTAGTGCACTCACGCAGTATAATTAATAACTAATTACTGTCGTTGACAGGACACGAGTAACTCGTCTATCTTCTGCAGGCTGCTTACGGTTTCGTCCGTGTTGCAGCCGATCATCAGCACATCTAGGTTTCGTCCGGGTGTGACCGAAAGGTAAGATGGAGAGCCTTGTCCCTGGTTTCAACGAGAAAACACACGTCCAACTCAGTTTGCCTGTTTTACAGGTTCGCGACGTGCTCGTACGTGGCTTTGGAGACTCCGTGGAGGAGGTCTTATCAGAGGCACGTCAACATCTTAAAGATGGCACTTGTGGCTTAGTAGAAGTTGAAAAAGGCGTTTTGCCTCAACTTGAACAGCCCTATGTGTTCATCAAACGTTCGGATGCTCGAACTGC

### 1-500 -ssRNA

GGGTTAAAGGTTTATACCTTCCCAGGTAACAAACCAACCAACTTTCGATCTCTTGTAGATCTGTTCTCTAAACGAACTTTAAAATCTGTGTGGCTGTCACTCGGCTGCATGCTTAGTGCACTCACGCAGTCGTTGACAGGACACGAGTAACTCGTCTATCTTCTGCAGGCTGCTTACGGTTTCGTCCGTGTTGCAGCCGATCATCAGCACATCTAGGTTTCGTCCGGGTGTGACCGAAAGGTAAGATGGAGAGCCTTGTCCCTGGTTTCAACGAGAAAACACACGTCCAACTCAGTTTGCCTGTTTTACAGGTTCGCGACGTGCTCGTACGTGGCTTTGGAGACTCCGTGGAGGAGGTCTTATCAGAGGCACGTCAACATCTTAAAGATGGCACTTGTGGCTTAGTAGAAGTTGAAAAAGGCGTTTTGCCTCAACTTGAACAGCCCTATGTGTTCATCAAACGTTCGGATGCTCGAACTGC

### 1-500 +dsRNAa

GGGTTAAAGGTTTATACCTTCCCAGGTAACAAACCAACCAACTTTCGATCTCTTGTAGATCTGTTCTCTAAACGTATTAATTATACTGTGTGGCTGTCACTCGGCTGCATGCTTAGTGCACTCACGCAGTATAATTAATAACTAATTACTGTCGTTGACAGGACACGAGTAACTCGTCTATCTTCTGCAGGCTGCTTACGGTTTCGTCCGTGTTGCAGCCGATCATCAGCACATCTAGGTTTCGTCCGGGTGTGACCGAAAGGTAAGATGGAGAGCCTTGTCCCTGGTTTCAACGACAGTAATTAGTTCCAACTCAGTTTGCCTGTTTTACAGGTTCGCGACGTGCTCGTACGTGGCTTTGGAGACTCCGTGGAGGAGGTCTTATCAGAGGCACGTCAACATCTTAAAGATGGCACTTGTGGCTTAGTAGAAGTTGAAAAAGGCGTTTTGCCTCAACTTGAACAGCCCTATGTGTTCATCAAACGTTCGGATGCTCGAACTGC

### 1-500 +dsRNAb

GGGTTAAAGGTTTATACCTTCCCAGGTAACAAACCAACCAACTTTCGATCTCTTGTAGATCTGTTCTCTAAACGAACTTTAAAATCAGTAATTAGTTATTAATTATACTGTGTGGCTGTCACTCGGCTGCATGCTTAGTGCACTCACGCAGTATAATTAATAACTAATTACTGTCGTTGACAGGACACGAGTAACTCGTCTATCTTCTGCAGGCTGCTTACGGTTTCGTCCGTGTTGCAGCCGATCATCAGCACATCTAGGTTTCGTCCGGGTGTGACCGAAAGGTAAGATGGAGAGCCTTGTCCCTGGTTTCAACGAGAAAACACACGTCCAACTCAGTTTGCCTGTTTTACAGGTTCGCGACGTGCTCGTACGTGGCTTTGGAGACTCCGTGGAGGAGGTCTTATCAGAGGCACGTCAACATCTTAAAGATGGCACTTGTGGCTTAGTAGAAGTTGAAAAAGGCGTTTTGCCTCAACTTGAACAGCCCTATGTGTTCATCAAACGTTCGGATGCTCGAACTGC

### 1-500 +dsRNAc

GGGTTAAAGGTTTATACCTTCCCAGGTAACAAACCAACCAACTTTCGATCTCTTGTAGATCTGTTCTCTAAACGAACTTTAAAATCTGTGTGGCTGTCACTCGGCTGCATGCTTAGTGCACTCACGCAGTATAATTAATAACTAATTACTGTCGTTGACAGGACACGAGTAACTCGTCTATCTTCTGCAGGCTGCTTACGGTTTCGTCCGTGTTGCAGCCGATCATCAGCACATCTAGGTTTCGTCCGGGTGTGACCGAAAGGTAAGATGGAGAGCCTTGTCCCTGGTTTCAACGACAGTAATTAGTTATTAATTATAGAAAACACACGTCCAACTCAGTTTGCCTGTTTTACAGGTTCGCGACGTGCTCGTACGTGGCTTTGGAGACTCCGTGGAGGAGGTCTTATCAGAGGCACGTCAACATCTTAAAGATGGCACTTGTGGCTTAGTAGAAGTTGAAAAAGGCGTTTTGCCTCAACTTGAACAGCCCTATGTGTTCATCAAACGTTCGGATGCTCGAACTGC

### 1-500 +dsRNAd

GGGTTAAAGGTTTATACCTTCCCAGGTAACAAACCAACCAACTTTCGATCTCTTGTAGATCTGTTCTCTAAACGAACTTTAAAATCGATTGCATCCTGTGTGGCTGTCACTCGGCTGCATGCTTAGTGCACTCACGCAGGATGCAATCGTATAATTAATAACTAATTACTGCGATTGCATCTCGTTGACAGGACACGAGTAACTCGTCTATCTTCTGCAGGCTGCTTACGGTTTCGTCCGTGTTGCAGCCGATCATCAGCACATCTAGGTTTCGTCCGGGTGTGACCGAAAGGTAAGATGGAGAGCCTTGTCCCTGGTTTCAACGAGATGCAATCGGAAAACACACGTCCAACTCAGTTTGCCTGTTTTACAGGTTCGCGACGTGCTCGTACGTGGCTTTGGAGACTCCGTGGAGGAGGTCTTATCAGAGGCACGTCAACATCTTAAAGATGGCACTTGTGGCTTAGTAGAAGTTGAAAAAGGCGTTTTGCCTCAACTTGAACAGCCCTATGTGTTCATCAAACGTTCGGATGCTCGAACTGC

### 1-500 +ssRNAa

GGGTTAAAGGTTTATACCTTCCCAGGTAACAAACCAACCAACTTTCGATCTCTTGTAGATCTGTTCTCTAAACGAACTTTAAAATCTGTGTGGCTGTCACTCGGCTGCATGCTTAGTGCACTCACGCAGTATAATTAATATACCCATACGATGTTCCAGATTACGCTACTAATTACTGTCGTTGACAGGACACGAGTAACTCGTCTATCTTCTGCAGGCTGCTTACGGTTTCGTCCGTGTTGCAGCCGATCATCAGCACATCTAGGTTTCGTCCGGGTGTGACCGAAAGGTAAGATGGAGAGCCTTGTCCCTGGTTTCAACGAGAAAACACACGTCCAACTCAGTTTGCCTGTTTTACAGGTTCGCGACGTGCTCGTACGTGGCTTTGGAGACTCCGTGGAGGAGGTCTTATCAGAGGCACGTCAACATCTTAAAGATGGCACTTGTGGCTTAGTAGAAGTTGAAAAAGGCGTTTTGCCTCAACTTGAACAGCCCTATGTGTTCATCAAACGTTCGGATGCTCGAACTGC

### 1-500 -dsRNA

GGGTTAAAGGTTTATACCTTCCCAGGTAACAAACCAACCAACTTTCGATCTCTTGTAGATCTGTTCTCTAAACGAACTTTAAAATGGTGTGTTCTGTCCAGCGTCTGCATGCTTAGTGCACTCACGCAGTATAATTAATAACTAATTACTGTATGGTGGCTTACGATCATAAAGTGCAGCGAGCCTGCAGGCTGCTTACGGTTTCGTCCGTGTTGCAGCCGATCATCAGCACATCTAGGTTTCGTCCGGGTGTGACCGAAAGGTAAGATGGAGAGCCTTGTCCCTGGTTTCAACGAGAAAACACACGTCCAACTCAGTTTGCCTGTTTTACAGGTTCGCGACGTGCTCGTACGTGGCTTTGGAGACTCCGTGGAGGAGGTCTTATCAGAGGCACGTCAACATCTTAAAGATGGCACTTGTGGCTTAGTAGAAGTTGAAAAAGGCGTTTTGCCTCAACTTGAACAGCCCTATGTGTTCATCAAACGTTCGGATGCTCGAACTGC

### 1-500 Restore pairing

GGGTTAAAGGTTTATACCTTCCCAGGTAACAAACCAACCAACTTTCGATCTCTTGTAGATCTGTTCTCTAAACGAACTTTAAAATGGTGTGTTCTGTCCAGCGTCTGCATGATTCTGGCACGACATACCTATAATTAATAACTAATTACTGTATGGTGGCTTACGATCATAAAGTGCAGCGAGCCTGCAGGCTGCTTACGGTTTCGTCCGTGTTGCAGCCGATCATCAGCACATCTAGGTTTCGTCCGGGTGTGACCGAAAGGTCCTCGCTGACTCTGGTCCAAGCGTTCACCATAGAAAACACACGTCCAACTCAGTTTGCCTGTTTTACAGGTTCGCGACGTGCTCGTACGTGGCTTTGGAGACTCCGTGGAGGAGGTCTTATCAGAGGCACGTCAACATCTTAAAGATGGCACTTGTGGCTTAGTAGAAGTTGAAAAAGGCGTTTTGCCTCAACTTGAACAGCCCTATGTGTTCATCAAACGTTCGGATGCTCGAACTGC

### 500-1000 Wildtype

GGGCACCTCATGGTCATGTTATGGTTGAGCTGGTAGCAGAACTCGAAGGCATTCAGTACGGTCGTAGTGGTGAGACACTTGGTGTCCTTGTCCCTCATGTGGGCGAAATACCAGTGGCTTACCGCAAGGTTCTTCTTCGTAAGAACGGTAATAAAGGAGCTGGTGGCCATAGTTACGGCGCCGATCTAAAGTCATTTGACTTAGGCGACGAGCTTGGCACTGATCCTTATGAAGATTTTCAAGAAAACTGGAACACTAAACATAGCAGTGGTGTTACCCGTGAACTCATGCGTGAGCTTAACGGAGGGGCATACACTCGCTATGTCGATAACAACTTCTGTGGCCCTGATGGCTACCCTCTTGAGTGCATTAAAGACCTTCTAGCACGTGCTGGTAAAGCTTCATGCACTTTGTCCGAACAACTGGACTTTATTGACACTAAGAGGGGTGTATACTGCTGCCGTGAACATGAGCATGAAATTGCTTGGTACACGGAACGTTCTGGGCCCTCGA

### 500-1000 -ssRNA

GGGCACCTCATGGTCATGTTATGGTTGAGCTGGTAGCAGAACTCGAAGGCATTCAGTACGGTCGTAGTGGTGAGACACTTGGTGTCCTTGTCCCTCATGTGGGCGAAATACCAGTGGCTTACCGCAAGGTTCTTCTTCGTAAGAACGGTAATAAAGGAGCTGGTGGCCATAGTTACGGCGCCGATCTAAAGTCATTTGACTTAGGCGACGAGCTTGGCACTGATCCTTACATAGCAGTGGTGTTACCCGTGAACTCATGCGTGAGCTTAACGGAGGGGCATACACTCGCTATGTCGATAACAACTTCTGTGGCCCTGATGGCTACCCTCTTGAGTGCATTAAAGACCTTCTAGCACGTGCTGGTAAAGCTTCATGCACTTTGTCCGAACAACTGGACTTTATTGACACTAAGAGGGGTGTATACTGCTGCCGTGAACATGAGCATGAAATTGCTTGGTACACGGAACGTTCTGGGCCCTCGA

### 500-1000 +dsRNAa

GGGCACCTCATGGTCATGTTATGGTTGAGCTGGTAGCAGAACTCGAAGGCATTCAGTACGGTCGTAGTGGTGAGACACTTGGTGTCCTTGTCCCTCATGTGGGCGAAATACCAGTGGCTTACCGCAAGGTTCTTCTTCCTTGAAAATCTTCATAAGGAGCTGGTGGCCATAGTTACGGCGCCGATCTAAAGTCATTTGACTTAGGCGACGAGCTTGGCACTGATCCTTATGAAGATTTTCAAGAAAACTGGAACACTAAACATAGCAGTGGTGTTACCCGTGAACTCATGCGTGAGCTTAACGGAGGGGCATACACTCGCTATGTTTAGTGTTCCAGTTTTGGCCCTGATGGCTACCCTCTTGAGTGCATTAAAGACCTTCTAGCACGTGCTGGTAAAGCTTCATGCACTTTGTCCGAACAACTGGACTTTATTGACACTAAGAGGGGTGTATACTGCTGCCGTGAACATGAGCATGAAATTGCTTGGTACACGGAACGTTCTGGGCCCTCGA

### 500-1000 +dsRNAb

GGGCACCTCATGGTCATGTTATGGTTGAGCTGGTAGCAGAACTCGAAGGCATTCAGTACGGTCGTAGTGGTGAGACACTTGGTGTCCTTGTCCCTCATGTGGGCGAAATACCAGTGGCTTACCGCAAGGTTCTTCTTCGTAAGAACGGTAATATTAGTGTTCCAGTTTTCTTGAAAATCTTCATAAGGAGCTGGTGGCCATAGTTACGGCGCCGATCTAAAGTCATTTGACTTAGGCGACGAGCTTGGCACTGATCCTTATGAAGATTTTCAAGAAAACTGGAACACTAAACATAGCAGTGGTGTTACCCGTGAACTCATGCGTGAGCTTAACGGAGGGGCATACACTCGCTATGTCGATAACAACTTCTGTGGCCCTGATGGCTACCCTCTTGAGTGCATTAAAGACCTTCTAGCACGTGCTGGTAAAGCTTCATGCACTTTGTCCGAACAACTGGACTTTATTGACACTAAGAGGGGTGTATACTGCTGCCGTGAACATGAGCATGAAATTGCTTGGTACACGGAACGTTCTGGGCCCTCGA

### 500-1000 +dsRNAc

GGGCACCTCATGGTCATGTTATGGTTGAGCTGGTAGCAGAACTCGAAGGCATTCAGTACGGTCGTAGTGGTGAGACACTTGGTGTCCTTGTCCCTCATGTGGGCGAAATACCAGTGGCTTACCGCAAGGTTCTTCTTCGTAAGAACGGTAATAAAGGAGCTGGTGGCCATAGTTACGGCGCCGATCTAAAGTCATTTGACTTAGGCGACGAGCTTGGCACTGATCCTTATGAAGATTTTCAAGAAAACTGGAACACTAAACATAGCAGTGGTGTTACCCGTGAACTCATGCGTGAGCTTAACGGAGGGGCATACACTCGCTATGTTTAGTGTTCCAGTTTTCTTGAAAATCTTCATCGATAACAACTTCTGTGGCCCTGATGGCTACCCTCTTGAGTGCATTAAAGACCTTCTAGCACGTGCTGGTAAAGCTTCATGCACTTTGTCCGAACAACTGGACTTTATTGACACTAAGAGGGGTGTATACTGCTGCCGTGAACATGAGCATGAAATTGCTTGGTACACGGAACGTTCTGGGCCCTCGA

### 500-1000 +dsRNAd

CACCTCATGGTCATGTTATGGTTGAGCTGGTAGCAGAACTCGAAGGCATTCAGTACGGTCGTAGTGGTGAGACACTTGGTGTCCTTGTCCCTCATGTGGGCGAAATACCAGTGGCTTACCGCAAGGTTCTTCTTCGTAAGAACGGTAATAAGATTGCATCAAGGAGCTGGTGGCCATAGTTACGGCGCCGATCTAAAGTCATTTGACTTAGGCGACGAGCTTGGCACTGATCCTTGATGCAATCTATGAAGATTTTCAAGAAAACTGGAACACTAACGATTGCATCACATAGCAGTGGTGTTACCCGTGAACTCATGCGTGAGCTTAACGGAGGGGCATACACTCGCTATGTGATGCAATCGCGATAACAACTTCTGTGGCCCTGATGGCTACCCTCTTGAGTGCATTAAAGACCTTCTAGCACGTGCTGGTAAAGCTTCATGCACTTTGTCCGAACAACTGGACTTTATTGACACTAAGAGGGGTGTATACTGCTGCCGTGAACATGAGCATGAAATTGCTTGGTACACGGAACGTTCTGGGCCCTCGA

### 500-1000 +ssRNAa

GGGCACCTCATGGTCATGTTATGGTTGAGCTGGTAGCAGAACTCGAAGGCATTCAGTACGGTCGTAGTGGTGAGACACTTGGTGTCCTTGTCCCTCATGTGGGCGAAATACCAGTGGCTTACCGCAAGGTTCTTCTTCGTAAGAACGGTAATAAAGGAGCTGGTGGCCATAGTTACGGCGCCGATCTAAAGTCATTTGACTTAGGCGACGAGCTTGGCACTGATCCTTATGAAGATTTTCAATACCCATACGATGTTCCAGATTACGCTGAAAACTGGAACACTAAACATAGCAGTGGTGTTACCCGTGAACTCATGCGTGAGCTTAACGGAGGGGCATACACTCGCTATGTCGATAACAACTTCTGTGGCCCTGATGGCTACCCTCTTGAGTGCATTAAAGACCTTCTAGCACGTGCTGGTAAAGCTTCATGCACTTTGTCCGAACAACTGGACTTTATTGACACTAAGAGGGGTGTATACTGCTGCCGTGAACATGAGCATGAAATTGCTTGGTACACGGAACGTTCTGGGCCCTCGA

### 500-1000 -dsRNA

GGGCACCTCATGGTCATGTTATGGTTGAGCTGGTAGCAGAACTCGAAGGCATTCAGTACGGTCGTAGTGGTGAGACACTTGGTGTCCTTGTCCCTCATGTGGGCGAAATACCAGTGGCTTACCGCAAGGTTCTTCTTCGTAAGAACGGTAATAGGAAGGCTGGTGCTGGTCTGCAACGGTAGGATCTAAAGTCATTTGACTTAGGCGACGAGCTTGGCACTGATCCTTATGAAGATTTTCAAGAAAACTGGAACACTAACACGCTACTGCTGTGGACGATGGAACTCATGCGTGAGCTTAACGGAGGGGCATACACTCGCTATGTCGATAACAACTTCTGTGGCCCTGATGGCTACCCTCTTGAGTGCATTAAAGACCTTCTAGCACGTGCTGGTAAAGCTTCATGCACTTTGTCCGAACAACTGGACTTTATTGACACTAAGAGGGGTGTATACTGCTGCCGTGAACATGAGCATGAAATTGCTTGGTACACGGAACGTTCTGGGCCCTCGA

### 500-1000 Restore pairing

GGGCACCTCATGGTCATGTTATGGTTGAGCTGGTAGCAGAACTCGAAGGCATTCAGTACGGTCGTAGTGGTGAGACACTTGGTGTCCTTGTCCCTCATGTGGGCGAAATACCAGTGGCTTACCGCAAGGTTCTTCTTCGTAAGAACGGTAATAGGAAGGCTGGTGCTGGTCTGCAACGGTAGGATCTAAAGTCATTTGACTTACTACAGTGTAGCCAGACTGACTTCCATGAAGATTTTCAAGAAAACTGGAACACTAACACGCTACTGCTGTGGACGATGGAACTCATGCGTGAGCTTACATCAGGTTACGAGCAGCTAGCGTGCGATAACAACTTCTGTGGCCCTGATGGCTACCCTCTTGAGTGCATTAAAGACCTTCTAGCACGTGCTGGTAAAGCTTCATGCACTTTGTCCGAACAACTGGACTTTATTGACACTAAGAGGGGTGTATACTGCTGCCGTGAACATGAGCATGAAATTGCTTGGTACACGGAACGTTCTGGGCCCTCGA

### 1-1000 Wildtype

GGGTTAAAGGTTTATACCTTCCCAGGTAACAAACCAACCAACTTTCGATCTCTTGTAGATCTGTTCTCTAAACGAACTTTAAAATCTGTGTGGCTGTCACTCGGCTGCATGCTTAGTGCACTCACGCAGTATAATTAATAACTAATTACTGTCGTTGACAGGACACGAGTAACTCGTCTATCTTCTGCAGGCTGCTTACGGTTTCGTCCGTGTTGCAGCCGATCATCAGCACATCTAGGTTTCGTCCGGGTGTGACCGAAAGGTAAGATGGAGAGCCTTGTCCCTGGTTTCAACGAGAAAACACACGTCCAACTCAGTTTGCCTGTTTTACAGGTTCGCGACGTGCTCGTACGTGGCTTTGGAGACTCCGTGGAGGAGGTCTTATCAGAGGCACGTCAACATCTTAAAGATGGCACTTGTGGCTTAGTAGAAGTTGAAAAAGGCGTTTTGCCTCAACTTGAACAGCCCTATGTGTTCATCAAACGTTCGGATGCTCGAACTGCACCTCATGGTCATGTTATGGTTGAGCTGGTAGCAGAACTCGAAGGCATTCAGTACGGTCGTAGTGGTGAGACACTTGGTGTCCTTGTCCCTCATGTGGGCGAAATACCAGTGGCTTACCGCAAGGTTCTTCTTCGTAAGAACGGTAATAAAGGAGCTGGTGGCCATAGTTACGGCGCCGATCTAAAGTCATTTGACTTAGGCGACGAGCTTGGCACTGATCCTTATGAAGATTTTCAAGAAAACTGGAACACTAAACATAGCAGTGGTGTTACCCGTGAACTCATGCGTGAGCTTAACGGAGGGGCATACACTCGCTATGTCGATAACAACTTCTGTGGCCCTGATGGCTACCCTCTTGAGTGCATTAAAGACCTTCTAGCACGTGCTGGTAAAGCTTCATGCACTTTGTCCGAACAACTGGACTTTATTGACACTAAGAGGGGTGTATACTGCTGCCGTGAACATGAGCATGAAATTGCTTGGTACACGGAACGTTCTGGGCCCTCGA

### 1-1000 -ssRNA

GGGTTAAAGGTTTATACCTTCCCAGGTAACAAACCAACCAACTTTCGATCTCTTGTAGATCTGTTCTCTAAACGAACTTTAAAATCTGTGTGGCTGTCACTCGGCTGCATGCTTAGTGCACTCACGCAGTCGTTGACAGGACACGAGTAACTCGTCTATCTTCTGCAGGCTGCTTACGGTTTCGTCCGTGTTGCAGCCGATCATCAGCACATCTAGGTTTCGTCCGGGTGTGACCGAAAGGTAAGATGGAGAGCCTTGTCCCTGGTTTCAACGAGAAAACACACGTCCAACTCAGTTTGCCTGTTTTACAGGTTCGCGACGTGCTCGTACGTGGCTTTGGAGACTCCGTGGAGGAGGTCTTATCAGAGGCACGTCAACATCTTAAAGATGGCACTTGTGGCTTAGTAGAAGTTGAAAAAGGCGTTTTGCCTCAACTTGAACAGCCCTATGTGTTCATCAAACGTTCGGATGCTCGAACTGCACCTCATGGTCATGTTATGGTTGAGCTGGTAGCAGAACTCGAAGGCATTCAGTACGGTCGTAGTGGTGAGACACTTGGTGTCCTTGTCCCTCATGTGGGCGAAATACCAGTGGCTTACCGCAAGGTTCTTCTTCGTAAGAACGGTAATAAAGGAGCTGGTGGCCATAGTTACGGCGCCGATCTAAAGTCATTTGACTTAGGCGACGAGCTTGGCACTGATCCTTACATAGCAGTGGTGTTACCCGTGAACTCATGCGTGAGCTTAACGGAGGGGCATACACTCGCTATGTCGATAACAACTTCTGTGGCCCTGATGGCTACCCTCTTGAGTGCATTAAAGACCTTCTAGCACGTGCTGGTAAAGCTTCATGCACTTTGTCCGAACAACTGGACTTTATTGACACTAAGAGGGGTGTATACTGCTGCCGTGAACATGAGCATGAAATTGCTTGGTACACGGAACGTTCTGGGCCCTCGA

### 1-1000 +dsRNAa

GGGTTAAAGGTTTATACCTTCCCAGGTAACAAACCAACCAACTTTCGATCTCTTGTAGATCTGTTCTCTAAACGTATTAATTATACTGTGTGGCTGTCACTCGGCTGCATGCTTAGTGCACTCACGCAGTATAATTAATAACTAATTACTGTCGTTGACAGGACACGAGTAACTCGTCTATCTTCTGCAGGCTGCTTACGGTTTCGTCCGTGTTGCAGCCGATCATCAGCACATCTAGGTTTCGTCCGGGTGTGACCGAAAGGTAAGATGGAGAGCCTTGTCCCTGGTTTCAACGACAGTAATTAGTTCCAACTCAGTTTGCCTGTTTTACAGGTTCGCGACGTGCTCGTACGTGGCTTTGGAGACTCCGTGGAGGAGGTCTTATCAGAGGCACGTCAACATCTTAAAGATGGCACTTGTGGCTTAGTAGAAGTTGAAAAAGGCGTTTTGCCTCAACTTGAACAGCCCTATGTGTTCATCAAACGTTCGGATGCTCGAACTGCACCTCATGGTCATGTTATGGTTGAGCTGGTAGCAGAACTCGAAGGCATTCAGTACGGTCGTAGTGGTGAGACACTTGGTGTCCTTGTCCCTCATGTGGGCGAAATACCAGTGGCTTACCGCAAGGTTCTTCTTCCTTGAAAATCTTCATAAGGAGCTGGTGGCCATAGTTACGGCGCCGATCTAAAGTCATTTGACTTAGGCGACGAGCTTGGCACTGATCCTTATGAAGATTTTCAAGAAAACTGGAACACTAAACATAGCAGTGGTGTTACCCGTGAACTCATGCGTGAGCTTAACGGAGGGGCATACACTCGCTATGTTTAGTGTTCCAGTTTTGGCCCTGATGGCTACCCTCTTGAGTGCATTAAAGACCTTCTAGCACGTGCTGGTAAAGCTTCATGCACTTTGTCCGAACAACTGGACTTTATTGACACTAAGAGGGGTGTATACTGCTGCCGTGAACATGAGCATGAAATTGCTTGGTACACGGAACGTTCTGGGCCCTCGA

### 1-1000 +dsRNAb

GGGTTAAAGGTTTATACCTTCCCAGGTAACAAACCAACCAACTTTCGATCTCTTGTAGATCTGTTCTCTAAACGAACTTTAAAATCAGTAATTAGTTATTAATTATACTGTGTGGCTGTCACTCGGCTGCATGCTTAGTGCACTCACGCAGTATAATTAATAACTAATTACTGTCGTTGACAGGACACGAGTAACTCGTCTATCTTCTGCAGGCTGCTTACGGTTTCGTCCGTGTTGCAGCCGATCATCAGCACATCTAGGTTTCGTCCGGGTGTGACCGAAAGGTAAGATGGAGAGCCTTGTCCCTGGTTTCAACGAGAAAACACACGTCCAACTCAGTTTGCCTGTTTTACAGGTTCGCGACGTGCTCGTACGTGGCTTTGGAGACTCCGTGGAGGAGGTCTTATCAGAGGCACGTCAACATCTTAAAGATGGCACTTGTGGCTTAGTAGAAGTTGAAAAAGGCGTTTTGCCTCAACTTGAACAGCCCTATGTGTTCATCAAACGTTCGGATGCTCGAACTGCACCTCATGGTCATGTTATGGTTGAGCTGGTAGCAGAACTCGAAGGCATTCAGTACGGTCGTAGTGGTGAGACACTTGGTGTCCTTGTCCCTCATGTGGGCGAAATACCAGTGGCTTACCGCAAGGTTCTTCTTCGTAAGAACGGTAATATTAGTGTTCCAGTTTTCTTGAAAATCTTCATAAGGAGCTGGTGGCCATAGTTACGGCGCCGATCTAAAGTCATTTGACTTAGGCGACGAGCTTGGCACTGATCCTTATGAAGATTTTCAAGAAAACTGGAACACTAAACATAGCAGTGGTGTTACCCGTGAACTCATGCGTGAGCTTAACGGAGGGGCATACACTCGCTATGTCGATAACAACTTCTGTGGCCCTGATGGCTACCCTCTTGAGTGCATTAAAGACCTTCTAGCACGTGCTGGTAAAGCTTCATGCACTTTGTCCGAACAACTGGACTTTATTGACACTAAGAGGGGTGTATACTGCTGCCGTGAACATGAGCATGAAATTGCTTGGTACACGGAACGTTCTGGGCCCTCGA

### 1-1000 +dsRNAc

GGGTTAAAGGTTTATACCTTCCCAGGTAACAAACCAACCAACTTTCGATCTCTTGTAGATCTGTTCTCTAAACGAACTTTAAAATCTGTGTGGCTGTCACTCGGCTGCATGCTTAGTGCACTCACGCAGTATAATTAATAACTAATTACTGTCGTTGACAGGACACGAGTAACTCGTCTATCTTCTGCAGGCTGCTTACGGTTTCGTCCGTGTTGCAGCCGATCATCAGCACATCTAGGTTTCGTCCGGGTGTGACCGAAAGGTAAGATGGAGAGCCTTGTCCCTGGTTTCAACGACAGTAATTAGTTATTAATTATAGAAAACACACGTCCAACTCAGTTTGCCTGTTTTACAGGTTCGCGACGTGCTCGTACGTGGCTTTGGAGACTCCGTGGAGGAGGTCTTATCAGAGGCACGTCAACATCTTAAAGATGGCACTTGTGGCTTAGTAGAAGTTGAAAAAGGCGTTTTGCCTCAACTTGAACAGCCCTATGTGTTCATCAAACGTTCGGATGCTCGAACTGCACCTCATGGTCATGTTATGGTTGAGCTGGTAGCAGAACTCGAAGGCATTCAGTACGGTCGTAGTGGTGAGACACTTGGTGTCCTTGTCCCTCATGTGGGCGAAATACCAGTGGCTTACCGCAAGGTTCTTCTTCGTAAGAACGGTAATAAAGGAGCTGGTGGCCATAGTTACGGCGCCGATCTAAAGTCATTTGACTTAGGCGACGAGCTTGGCACTGATCCTTATGAAGATTTTCAAGAAAACTGGAACACTAAACATAGCAGTGGTGTTACCCGTGAACTCATGCGTGAGCTTAACGGAGGGGCATACACTCGCTATGTTTAGTGTTCCAGTTTTCTTGAAAATCTTCATCGATAACAACTTCTGTGGCCCTGATGGCTACCCTCTTGAGTGCATTAAAGACCTTCTAGCACGTGCTGGTAAAGCTTCATGCACTTTGTCCGAACAACTGGACTTTATTGACACTAAGAGGGGTGTATACTGCTGCCGTGAACATGAGCATGAAATTGCTTGGTACACGGAACGTTCTGGGCCCTCGA

### 1-1000 +dsRNAd

GGGTTAAAGGTTTATACCTTCCCAGGTAACAAACCAACCAACTTTCGATCTCTTGTAGATCTGTTCTCTAAACGAACTTTAAAATCGATTGCATCCTGTGTGGCTGTCACTCGGCTGCATGCTTAGTGCACTCACGCAGGATGCAATCGTATAATTAATAACTAATTACTGCGATTGCATCTCGTTGACAGGACACGAGTAACTCGTCTATCTTCTGCAGGCTGCTTACGGTTTCGTCCGTGTTGCAGCCGATCATCAGCACATCTAGGTTTCGTCCGGGTGTGACCGAAAGGTAAGATGGAGAGCCTTGTCCCTGGTTTCAACGAGATGCAATCGGAAAACACACGTCCAACTCAGTTTGCCTGTTTTACAGGTTCGCGACGTGCTCGTACGTGGCTTTGGAGACTCCGTGGAGGAGGTCTTATCAGAGGCACGTCAACATCTTAAAGATGGCACTTGTGGCTTAGTAGAAGTTGAAAAAGGCGTTTTGCCTCAACTTGAACAGCCCTATGTGTTCATCAAACGTTCGGATGCTCGAACTGCACCTCATGGTCATGTTATGGTTGAGCTGGTAGCAGAACTCGAAGGCATTCAGTACGGTCGTAGTGGTGAGACACTTGGTGTCCTTGTCCCTCATGTGGGCGAAATACCAGTGGCTTACCGCAAGGTTCTTCTTCGTAAGAACGGTAATAAGATTGCATCAAGGAGCTGGTGGCCATAGTTACGGCGCCGATCTAAAGTCATTTGACTTAGGCGACGAGCTTGGCACTGATCCTTGATGCAATCTATGAAGATTTTCAAGAAAACTGGAACACTAACGATTGCATCACATAGCAGTGGTGTTACCCGTGAACTCATGCGTGAGCTTAACGGAGGGGCATACACTCGCTATGTGATGCAATCGCGATAACAACTTCTGTGGCCCTGATGGCTACCCTCTTGAGTGCATTAAAGACCTTCTAGCACGTGCTGGTAAAGCTTCATGCACTTTGTCCGAACAACTGGACTTTATTGACACTAAGAGGGGTGTATACTGCTGCCGTGAACATGAGCATGAAATTGCTTGGTACACGGAACGTTCTGGGCCCTCGA

### 1-1000 +ssRNAa

GGGTTAAAGGTTTATACCTTCCCAGGTAACAAACCAACCAACTTTCGATCTCTTGTAGATCTGTTCTCTAAACGAACTTTAAAATCTGTGTGGCTGTCACTCGGCTGCATGCTTAGTGCACTCACGCAGTATAATTAATATACCCATACGATGTTCCAGATTACGCTACTAATTACTGTCGTTGACAGGACACGAGTAACTCGTCTATCTTCTGCAGGCTGCTTACGGTTTCGTCCGTGTTGCAGCCGATCATCAGCACATCTAGGTTTCGTCCGGGTGTGACCGAAAGGTAAGATGGAGAGCCTTGTCCCTGGTTTCAACGAGAAAACACACGTCCAACTCAGTTTGCCTGTTTTACAGGTTCGCGACGTGCTCGTACGTGGCTTTGGAGACTCCGTGGAGGAGGTCTTATCAGAGGCACGTCAACATCTTAAAGATGGCACTTGTGGCTTAGTAGAAGTTGAAAAAGGCGTTTTGCCTCAACTTGAACAGCCCTATGTGTTCATCAAACGTTCGGATGCTCGAACTGCACCTCATGGTCATGTTATGGTTGAGCTGGTAGCAGAACTCGAAGGCATTCAGTACGGTCGTAGTGGTGAGACACTTGGTGTCCTTGTCCCTCATGTGGGCGAAATACCAGTGGCTTACCGCAAGGTTCTTCTTCGTAAGAACGGTAATAAAGGAGCTGGTGGCCATAGTTACGGCGCCGATCTAAAGTCATTTGACTTAGGCGACGAGCTTGGCACTGATCCTTATGAAGATTTTCAATACCCATACGATGTTCCAGATTACGCTGAAAACTGGAACACTAAACATAGCAGTGGTGTTACCCGTGAACTCATGCGTGAGCTTAACGGAGGGGCATACACTCGCTATGTCGATAACAACTTCTGTGGCCCTGATGGCTACCCTCTTGAGTGCATTAAAGACCTTCTAGCACGTGCTGGTAAAGCTTCATGCACTTTGTCCGAACAACTGGACTTTATTGACACTAAGAGGGGTGTATACTGCTGCCGTGAACATGAGCATGAAATTGCTTGGTACACGGAACGTTCTGGGCCCTCGA

### 1-1000 -dsRNA

GGGTTAAAGGTTTATACCTTCCCAGGTAACAAACCAACCAACTTTCGATCTCTTGTAGATCTGTTCTCTAAACGAACTTTAAAATGGTGTGTTCTGTCCAGCGTCTGCATGCTTAGTGCACTCACGCAGTATAATTAATAACTAATTACTGTATGGTGGCTTACGATCATAAAGTGCAGCGAGCCTGCAGGCTGCTTACGGTTTCGTCCGTGTTGCAGCCGATCATCAGCACATCTAGGTTTCGTCCGGGTGTGACCGAAAGGTAAGATGGAGAGCCTTGTCCCTGGTTTCAACGAGAAAACACACGTCCAACTCAGTTTGCCTGTTTTACAGGTTCGCGACGTGCTCGTACGTGGCTTTGGAGACTCCGTGGAGGAGGTCTTATCAGAGGCACGTCAACATCTTAAAGATGGCACTTGTGGCTTAGTAGAAGTTGAAAAAGGCGTTTTGCCTCAACTTGAACAGCCCTATGTGTTCATCAAACGTTCGGATGCTCGAACTGCACCTCATGGTCATGTTATGGTTGAGCTGGTAGCAGAACTCGAAGGCATTCAGTACGGTCGTAGTGGTGAGACACTTGGTGTCCTTGTCCCTCATGTGGGCGAAATACCAGTGGCTTACCGCAAGGTTCTTCTTCGTAAGAACGGTAATAGGAAGGCTGGTGCTGGTCTGCAACGGTAGGATCTAAAGTCATTTGACTTAGGCGACGAGCTTGGCACTGATCCTTATGAAGATTTTCAAGAAAACTGGAACACTAACACGCTACTGCTGTGGACGATGGAACTCATGCGTGAGCTTAACGGAGGGGCATACACTCGCTATGTCGATAACAACTTCTGTGGCCCTGATGGCTACCCTCTTGAGTGCATTAAAGACCTTCTAGCACGTGCTGGTAAAGCTTCATGCACTTTGTCCGAACAACTGGACTTTATTGACACTAAGAGGGGTGTATACTGCTGCCGTGAACATGAGCATGAAATTGCTTGGTACACGGAACGTTCTGGGCCCTCGA

### 1-1000 Restore pairing

GGGTTAAAGGTTTATACCTTCCCAGGTAACAAACCAACCAACTTTCGATCTCTTGTAGATCTGTTCTCTAAACGAACTTTAAAATGGTGTGTTCTGTCCAGCGTCTGCATGATTCTGGCACGACATACCTATAATTAATAACTAATTACTGTATGGTGGCTTACGATCATAAAGTGCAGCGAGCCTGCAGGCTGCTTACGGTTTCGTCCGTGTTGCAGCCGATCATCAGCACATCTAGGTTTCGTCCGGGTGTGACCGAAAGGTCCTCGCTGACTCTGGTCCAAGCGTTCACCATAGAAAACACACGTCCAACTCAGTTTGCCTGTTTTACAGGTTCGCGACGTGCTCGTACGTGGCTTTGGAGACTCCGTGGAGGAGGTCTTATCAGAGGCACGTCAACATCTTAAAGATGGCACTTGTGGCTTAGTAGAAGTTGAAAAAGGCGTTTTGCCTCAACTTGAACAGCCCTATGTGTTCATCAAACGTTCGGATGCTCGAACTGCACCTCATGGTCATGTTATGGTTGAGCTGGTAGCAGAACTCGAAGGCATTCAGTACGGTCGTAGTGGTGAGACACTTGGTGTCCTTGTCCCTCATGTGGGCGAAATACCAGTGGCTTACCGCAAGGTTCTTCTTCGTAAGAACGGTAATAGGAAGGCTGGTGCTGGTCTGCAACGGTAGGATCTAAAGTCATTTGACTTACTACAGTGTAGCCAGACTGACTTCCATGAAGATTTTCAAGAAAACTGGAACACTAACACGCTACTGCTGTGGACGATGGAACTCATGCGTGAGCTTACATCAGGTTACGAGCAGCTAGCGTGCGATAACAACTTCTGTGGCCCTGATGGCTACCCTCTTGAGTGCATTAAAGACCTTCTAGCACGTGCTGGTAAAGCTTCATGCACTTTGTCCGAACAACTGGACTTTATTGACACTAAGAGGGGTGTATACTGCTGCCGTGAACATGAGCATGAAATTGCTTGGTACACGGAACGTTCTGGGCCCTCGA

### 1-1000 +dsRNA1b

GGGTTAAAGGTTTATACCTTCCCAGGTAACAAACCAACCAACTTTCGATCTCTTGTAGATCTGTTCTCTAAACGAACTTTAAAATCAGTAATTAGTTATTAATTATACTGTGTGGCTGTCACTCGGCTGCATGCTTAGTGCACTCACGCAGTATAATTAATAACTAATTACTGTCGTTGACAGGACACGAGTAACTCGTCTATCTTCTGCAGGCTGCTTACGGTTTCGTCCGTGTTGCAGCCGATCATCAGCACATCTAGGTTTCGTCCGGGTGTGACCGAAAGGTAAGATGGAGAGCCTTGTCCCTGGTTTCAACGAGAAAACACACGTCCAACTCAGTTTGCCTGTTTTACAGGTTCGCGACGTGCTCGTACGTGGCTTTGGAGACTCCGTGGAGGAGGTCTTATCAGAGGCACGTCAACATCTTAAAGATGGCACTTGTGGCTTAGTAGAAGTTGAAAAAGGCGTTTTGCCTCAACTTGAACAGCCCTATGTGTTCATCAAACGTTCGGATGCTCGAACTGCACCTCATGGTCATGTTATGGTTGAGCTGGTAGCAGAACTCGAAGGCATTCAGTACGGTCGTAGTGGTGAGACACTTGGTGTCCTTGTCCCTCATGTGGGCGAAATACCAGTGGCTTACCGCAAGGTTCTTCTTCGTAAGAACGGTAATAAAGGAGCTGGTGGCCATAGTTACGGCGCCGATCTAAAGTCATTTGACTTAGGCGACGAGCTTGGCACTGATCCTTATGAAGATTTTCAAGAAAACTGGAACACTAAACATAGCAGTGGTGTTACCCGTGAACTCATGCGTGAGCTTAACGGAGGGGCATACACTCGCTATGTCGATAACAACTTCTGTGGCCCTGATGGCTACCCTCTTGAGTGCATTAAAGACCTTCTAGCACGTGCTGGTAAAGCTTCATGCACTTTGTCCGAACAACTGGACTTTATTGACACTAAGAGGGGTGTATACTGCTGCCGTGAACATGAGCATGAAATTGCTTGGTACACGGAACGTTCTGGGCCCTCGA

### 1-1000 +dsRNA1c

GGGTTAAAGGTTTATACCTTCCCAGGTAACAAACCAACCAACTTTCGATCTCTTGTAGATCTGTTCTCTAAACGAACTTTAAAATCTGTGTGGCTGTCACTCGGCTGCATGCTTAGTGCACTCACGCAGTATAATTAATAACTAATTACTGTCGTTGACAGGACACGAGTAACTCGTCTATCTTCTGCAGGCTGCTTACGGTTTCGTCCGTGTTGCAGCCGATCATCAGCACATCTAGGTTTCGTCCGGGTGTGACCGAAAGGTAAGATGGAGAGCCTTGTCCCTGGTTTCAACGACAGTAATTAGTTATTAATTATAGAAAACACACGTCCAACTCAGTTTGCCTGTTTTACAGGTTCGCGACGTGCTCGTACGTGGCTTTGGAGACTCCGTGGAGGAGGTCTTATCAGAGGCACGTCAACATCTTAAAGATGGCACTTGTGGCTTAGTAGAAGTTGAAAAAGGCGTTTTGCCTCAACTTGAACAGCCCTATGTGTTCATCAAACGTTCGGATGCTCGAACTGCACCTCATGGTCATGTTATGGTTGAGCTGGTAGCAGAACTCGAAGGCATTCAGTACGGTCGTAGTGGTGAGACACTTGGTGTCCTTGTCCCTCATGTGGGCGAAATACCAGTGGCTTACCGCAAGGTTCTTCTTCGTAAGAACGGTAATAAAGGAGCTGGTGGCCATAGTTACGGCGCCGATCTAAAGTCATTTGACTTAGGCGACGAGCTTGGCACTGATCCTTATGAAGATTTTCAAGAAAACTGGAACACTAAACATAGCAGTGGTGTTACCCGTGAACTCATGCGTGAGCTTAACGGAGGGGCATACACTCGCTATGTCGATAACAACTTCTGTGGCCCTGATGGCTACCCTCTTGAGTGCATTAAAGACCTTCTAGCACGTGCTGGTAAAGCTTCATGCACTTTGTCCGAACAACTGGACTTTATTGACACTAAGAGGGGTGTATACTGCTGCCGTGAACATGAGCATGAAATTGCTTGGTACACGGAACGTTCTGGGCCCTCGA

### 1-1000 +dsRNA2b

GGGTTAAAGGTTTATACCTTCCCAGGTAACAAACCAACCAACTTTCGATCTCTTGTAGATCTGTTCTCTAAACGAACTTTAAAATCTGTGTGGCTGTCACTCGGCTGCATGCTTAGTGCACTCACGCAGTATAATTAATAACTAATTACTGTCGTTGACAGGACACGAGTAACTCGTCTATCTTCTGCAGGCTGCTTACGGTTTCGTCCGTGTTGCAGCCGATCATCAGCACATCTAGGTTTCGTCCGGGTGTGACCGAAAGGTAAGATGGAGAGCCTTGTCCCTGGTTTCAACGAGAAAACACACGTCCAACTCAGTTTGCCTGTTTTACAGGTTCGCGACGTGCTCGTACGTGGCTTTGGAGACTCCGTGGAGGAGGTCTTATCAGAGGCACGTCAACATCTTAAAGATGGCACTTGTGGCTTAGTAGAAGTTGAAAAAGGCGTTTTGCCTCAACTTGAACAGCCCTATGTGTTCATCAAACGTTCGGATGCTCGAACTGCACCTCATGGTCATGTTATGGTTGAGCTGGTAGCAGAACTCGAAGGCATTCAGTACGGTCGTAGTGGTGAGACACTTGGTGTCCTTGTCCCTCATGTGGGCGAAATACCAGTGGCTTACCGCAAGGTTCTTCTTCGTAAGAACGGTAATATTAGTGTTCCAGTTTTCTTGAAAATCTTCATAAGGAGCTGGTGGCCATAGTTACGGCGCCGATCTAAAGTCATTTGACTTAGGCGACGAGCTTGGCACTGATCCTTATGAAGATTTTCAAGAAAACTGGAACACTAAACATAGCAGTGGTGTTACCCGTGAACTCATGCGTGAGCTTAACGGAGGGGCATACACTCGCTATGTCGATAACAACTTCTGTGGCCCTGATGGCTACCCTCTTGAGTGCATTAAAGACCTTCTAGCACGTGCTGGTAAAGCTTCATGCACTTTGTCCGAACAACTGGACTTTATTGACACTAAGAGGGGTGTATACTGCTGCCGTGAACATGAGCATGAAATTGCTTGGTACACGGAACGTTCTGGGCCCTCGA

### 1-1000 +dsRNA2c

GGGTTAAAGGTTTATACCTTCCCAGGTAACAAACCAACCAACTTTCGATCTCTTGTAGATCTGTTCTCTAAACGAACTTTAAAATCTGTGTGGCTGTCACTCGGCTGCATGCTTAGTGCACTCACGCAGTATAATTAATAACTAATTACTGTCGTTGACAGGACACGAGTAACTCGTCTATCTTCTGCAGGCTGCTTACGGTTTCGTCCGTGTTGCAGCCGATCATCAGCACATCTAGGTTTCGTCCGGGTGTGACCGAAAGGTAAGATGGAGAGCCTTGTCCCTGGTTTCAACGAGAAAACACACGTCCAACTCAGTTTGCCTGTTTTACAGGTTCGCGACGTGCTCGTACGTGGCTTTGGAGACTCCGTGGAGGAGGTCTTATCAGAGGCACGTCAACATCTTAAAGATGGCACTTGTGGCTTAGTAGAAGTTGAAAAAGGCGTTTTGCCTCAACTTGAACAGCCCTATGTGTTCATCAAACGTTCGGATGCTCGAACTGCACCTCATGGTCATGTTATGGTTGAGCTGGTAGCAGAACTCGAAGGCATTCAGTACGGTCGTAGTGGTGAGACACTTGGTGTCCTTGTCCCTCATGTGGGCGAAATACCAGTGGCTTACCGCAAGGTTCTTCTTCGTAAGAACGGTAATAAAGGAGCTGGTGGCCATAGTTACGGCGCCGATCTAAAGTCATTTGACTTAGGCGACGAGCTTGGCACTGATCCTTATGAAGATTTTCAAGAAAACTGGAACACTAAACATAGCAGTGGTGTTACCCGTGAACTCATGCGTGAGCTTAACGGAGGGGCATACACTCGCTATGTTTAGTGTTCCAGTTTTCTTGAAAATCTTCATCGATAACAACTTCTGTGGCCCTGATGGCTACCCTCTTGAGTGCATTAAAGACCTTCTAGCACGTGCTGGTAAAGCTTCATGCACTTTGTCCGAACAACTGGACTTTATTGACACTAAGAGGGGTGTATACTGCTGCCGTGAACATGAGCATGAAATTGCTTGGTACACGGAACGTTCTGGGCCCTCGA

### 500-1000 +ssRNAb

GGGCACCTCATGGTCATGTTATGGTTGAGCTGGTAGCAGAACTCGAAGGCATTCAGTACGGTCGTAGTGGTGAGACACTTGGTGTCCTTGTCCCTCATGTGGGCGAAATACCAGTGGCTTACCGCAAGGTTCTTCTTCGTAAGAACGGTAATAAAGGAGCTGGTGGCCATAGTTACGGCGCCGATCTAAAGTCATTTGACTTAGGCGACGAGCTTGGCACTGATCCTTATGAAGATTTTCAAGAACAAAAACTCATCTCAGAAGAGGATCTGGAAAACTGGAACACTAAACATAGCAGTGGTGTTACCCGTGAACTCATGCGTGAGCTTAACGGAGGGGCATACACTCGCTATGTCGATAACAACTTCTGTGGCCCTGATGGCTACCCTCTTGAGTGCATTAAAGACCTTCTAGCACGTGCTGGTAAAGCTTCATGCACTTTGTCCGAACAACTGGACTTTATTGACACTAAGAGGGGTGTATACTGCTGCCGTGAACATGAGCATGAAATTGCTTGGTACACGGAACGTTCTGGGCCCTCGA

### 500-1000 +ssRNAc

GGGCACCTCATGGTCATGTTATGGTTGAGCTGGTAGCAGAACTCGAAGGCATTCAGTACGGTCGTAGTGGTGAGACACTTGGTGTCCTTGTCCCTCATGTGGGCGAAATACCAGTGGCTTACCGCAAGGTTCTTCTTCGTAAGAACGGTAATAAAGGAGCTGGTGGCCATAGTTACGGCGCCGATCTAAAGTCATTTGACTTAGGCGACGAGCTTGGCACTGATCCTTATGAAGATTTTCAAGAAAACTGGAACACTAAACATAGCAGTGGTGTTACCCGTGAACTCATGCGTGAGCTTAACGGAGGGGCATACACTCGCTATGTCGATAACAACTTCTGTGGCCCTGATGGCTACCCTCTTGAGTGCATTAAAGACCTTCTAGCACGTGCTGGTAAAGCTTCATGCACTTTGTCCGAACAACTGGACTTTATTGACACTAAGAGGGGTGTATACTGCTGCCGTGAACATGAGCATGAAATTGCTTGGTACACGGAACGTTCTTACCCATACGATGTTCCAGATTACGCTGGGCCCTCGA

### 500-1000 +ssRNAd

GGGCACCTCATGGTCATGTTATGGTTGAGCTGGTAGCAGAACTCGAAGGCATTCAGTACGGTCGTAGTGGTGAGACACTTGGTGTCCTTGTCCCTCATGTGGGCGAAATACCAGTGGCTTACCGCAAGGTTCTTCTTCGTAAGAACGGTAATAAAGGAGCTGGTGGCCATAGTTACGGCGCCGATCTAAAGTCATTTGACTTAGGCGACGAGCTTGGCACTGATCCTTATGAAGATTTTCAAGAAAACTGGAACACTAAACATAGCAGTGGTGTTACCCGTGAACTCATGCGTGAGCTTAACGGAGGGGCATACACTCGCTATGTCGATAACAACTTCTGTGGCCCTGATGGCTACCCTCTTGAGTGCATTAAAGACCTTCTAGCACGTGCTGGTAAAGCTTCATGCACTTTGTCCGAACAACTGGACTTTATTGACACTAAGAGGGGTGTATACTGCTGCCGTGAACATGAGCATGAAATTGCTTGGTACACGGAACGTTCTGAACAAAAACTCATCTCAGAAGAGGATCTGGGGCCCTCGA

### Start Mutant

GGGTTAAAGGTTTATACCTTCCCAGGTAACAAACCAACCAACTTTCGATCTCTTGTAGATCTGTTCTCTAAACGAACTTTAAAATCTGTGTGGCTGTCACTCGGCTGCATGCTTAGTGCACTCACGCAGTATAATTAATAACTAATTACTGTCGTTGACAGGACACGAGTAACTCGTCTATCTTCTGCAGGCTGCTTACGGTTTCGTCCGTGTTGCAGCCGATCATCAGCACATCTAGGTTTCGTCCGGGTGTGACCGAAAGGTAAGGTGGAGAGCCTTGTCCCTGGTTTCAACGAGAAAACACACGTCCAACTCAGTTTGCCTGTTTTACAGGTTCGCGACGTGCTCGTACGTGGCTTTGGAGACTCCGTGGAGGAGGTCTTATCAGAGGCACGTCAACATCTTAAAGATGGCACTTGTGGCTTAGTAGAAGTTGAAAAAGGCGTTTTGCCTCAACTTGAACAGCCCTATGTGTTCATCAAACGTTCGGATGCTCGAACTGCACCTCATGGTCATGTTATGGTTGAGCTGGTAGCAGAACTCGAAGGCATTCAGTACGGTCGTAGTGGTGAGACACTTGGTGTCCTTGTCCCTCATGTGGGCGAAATACCAGTGGCTTACCGCAAGGTTCTTCTTCGTAAGAACGGTAATAAAGGAGCTGGTGGCCATAGTTACGGCGCCGATCTAAAGTCATTTGACTTAGGCGACGAGCTTGGCACTGATCCTTATGAAGATTTTCAAGAAAACTGGAACACTAAACATAGCAGTGGTGTTACCCGTGAACTCATGCGTGAGCTTAACGGAGGGGCATACACTCGCTATGTCGATAACAACTTCTGTGGCCCTGATGGCTACCCTCTTGAGTGCATTAAAGACCTTCTAGCACGTGCTGGTAAAGCTTCATGCACTTTGTCCGAACAACTGGACTTTATTGACACTAAGAGGGGTGTATACTGCTGCCGTGAACATGAGCATGAAATTGCTTGGTACACGGAACGTTCTGGGCCCTCGA

### 1-1000 TRS-Del

GGGTTAAAGGTTTATACCTTCCCAGGTAACAAACCAACCAACTTTCGATCTCTTGTAGATCTTTTAAAATCTGTGTGGCTGTCACTCGGCTGCATGCTTAGTGCACTCACGCAGTATAATTAATAACTAATTACTGTCGTTGACAGGACACGAGTAACTCGTCTATCTTCTGCAGGCTGCTTACGGTTTCGTCCGTGTTGCAGCCGATCATCAGCACATCTAGGTTTCGTCCGGGTGTGACCGAAAGGTAAGATGGAGAGCCTTGTCCCTGGTTTCAACGAGAAAACACACGTCCAACTCAGTTTGCCTGTTTTACAGGTTCGCGACGTGCTCGTACGTGGCTTTGGAGACTCCGTGGAGGAGGTCTTATCAGAGGCACGTCAACATCTTAAAGATGGCACTTGTGGCTTAGTAGAAGTTGAAAAAGGCGTTTTGCCTCAACTTGAACAGCCCTATGTGTTCATCAAACGTTCGGATGCTCGAACTGCACCTCATGGTCATGTTATGGTTGAGCTGGTAGCAGAACTCGAAGGCATTCAGTACGGTCGTAGTGGTGAGACACTTGGTGTCCTTGTCCCTCATGTGGGCGAAATACCAGTGGCTTACCGCAAGGTTCTTCTTCGTAAGAACGGTAATAAAGGAGCTGGTGGCCATAGTTACGGCGCCGATCTAAAGTCATTTGACTTAGGCGACGAGCTTGGCACTGATCCTTATGAAGATTTTCAAGAAAACTGGAACACTAAACATAGCAGTGGTGTTACCCGTGAACTCATGCGTGAGCTTAACGGAGGGGCATACACTCGCTATGTCGATAACAACTTCTGTGGCCCTGATGGCTACCCTCTTGAGTGCATTAAAGACCTTCTAGCACGTGCTGGTAAAGCTTCATGCACTTTGTCCGAACAACTGGACTTTATTGACACTAAGAGGGGTGTATACTGCTGCCGTGAACATGAGCATGAAATTGCTTGGTACACGGAACGTTCTGGGCCCTCGA

### 1-1000 Add TRS 3′

GGGTTAAAGGTTTATACCTTCCCAGGTAACAAACCAACCAACTTTCGATCTCTTGTAGATCTGTTCTCTAAACGAACTTTAAAATCTGTGTGGCTGTCACTCGGCTGCATGCTTAGTGCACTCACGCAGTATAATTAATAACTAATTACTGTCGTTGACAGGACACGAGTAACTCGTCTATCTTCTGCAGGCTGCTTACGGTTTCGTCCGTGTTGCAGCCGATCATCAGCACATCTAGGTTTCGTCCGGGTGTGACCGAAAGGTAAGATGGAGAGCCTTGTCCCTGGTTTCAACGAGAAAACACACGTCCAACTCAGTTTGCCTGTTTTACAGGTTCGCGACGTGCTCGTACGTGGCTTTGGAGACTCCGTGGAGGAGGTCTTATCAGAGGCACGTCAACATCTTAAAGATGGCACTTGTGGCTTAGTAGAAGTTGAAAAAGGCGTTTTGCCTCAACTTGAACAGCCCTATGTGTTCATCAAACGTTCGGATGCTCGAACTGCACCTCATGGTCATGTTATGGTTGAGCTGGTAGCAGAACTCGAAGGCATTCAGTACGGTCGTAGTGGTGAGACACTTGGTGTCCTTGTCCCTCATGTGGGCGAAATACCAGTGGCTTACCGCAAGGTTCTTCTTCGTAAGAACGGTAATAAAGGAGCTGGTGGCCATAGTTACGGCGCCGATCTAAAGTCATTTGACTTAGGCGACGAGCTTGGCACTGATCCTTATGAAGATTTTCAAGAAAACTGGAACACTAAACATAGCAGTGGTGTTACCCGTGAACTCATGCGTGAGCTTAACGGAGGGGCATACACTCGCTATGTCGATAACAACTTCTGTGGCCCTGATGGCTACCCTCTTGAGTGCATTAAAGACCTTCTAGCACGTGCTGGTAAAGCTTCATGCACTTTGTCCGAACAACTGGACTTTATTGACACTAAGAGGGGTGTATACTGCTGCCGTGAACATGAGCATGAAATTGCTTGGTACACGGAACGTTCTGTTCTCTAAACGAACGGGCCCTCGA

### 1-1000 A68U

GGGTTAAAGGTTTATACCTTCCCAGGTAACAAACCAACCAACTTTCGATCTCTTGTAGATCTGTTCTCTTAACGAACTTTAAAATCTGTGTGGCTGTCACTCGGCTGCATGCTTAGTGCACTCACGCAGTATAATTAATAACTAATTACTGTCGTTGACAGGACACGAGTAACTCGTCTATCTTCTGCAGGCTGCTTACGGTTTCGTCCGTGTTGCAGCCGATCATCAGCACATCTAGGTTTCGTCCGGGTGTGACCGAAAGGTAAGATGGAGAGCCTTGTCCCTGGTTTCAACGAGAAAACACACGTCCAACTCAGTTTGCCTGTTTTACAGGTTCGCGACGTGCTCGTACGTGGCTTTGGAGACTCCGTGGAGGAGGTCTTATCAGAGGCACGTCAACATCTTAAAGATGGCACTTGTGGCTTAGTAGAAGTTGAAAAAGGCGTTTTGCCTCAACTTGAACAGCCCTATGTGTTCATCAAACGTTCGGATGCTCGAACTGCACCTCATGGTCATGTTATGGTTGAGCTGGTAGCAGAACTCGAAGGCATTCAGTACGGTCGTAGTGGTGAGACACTTGGTGTCCTTGTCCCTCATGTGGGCGAAATACCAGTGGCTTACCGCAAGGTTCTTCTTCGTAAGAACGGTAATAAAGGAGCTGGTGGCCATAGTTACGGCGCCGATCTAAAGTCATTTGACTTAGGCGACGAGCTTGGCACTGATCCTTATGAAGATTTTCAAGAAAACTGGAACACTAAACATAGCAGTGGTGTTACCCGTGAACTCATGCGTGAGCTTAACGGAGGGGCATACACTCGCTATGTCGATAACAACTTCTGTGGCCCTGATGGCTACCCTCTTGAGTGCATTAAAGACCTTCTAGCACGTGCTGGTAAAGCTTCATGCACTTTGTCCGAACAACTGGACTTTATTGACACTAAGAGGGGTGTATACTGCTGCCGTGAACATGAGCATGAAATTGCTTGGTACACGGAACGTTCTGGGCCCTCGA

### 1-1000 A69U

GGGTTAAAGGTTTATACCTTCCCAGGTAACAAACCAACCAACTTTCGATCTCTTGTAGATCTGTTCTCTATACGAACTTTAAAATCTGTGTGGCTGTCACTCGGCTGCATGCTTAGTGCACTCACGCAGTATAATTAATAACTAATTACTGTCGTTGACAGGACACGAGTAACTCGTCTATCTTCTGCAGGCTGCTTACGGTTTCGTCCGTGTTGCAGCCGATCATCAGCACATCTAGGTTTCGTCCGGGTGTGACCGAAAGGTAAGATGGAGAGCCTTGTCCCTGGTTTCAACGAGAAAACACACGTCCAACTCAGTTTGCCTGTTTTACAGGTTCGCGACGTGCTCGTACGTGGCTTTGGAGACTCCGTGGAGGAGGTCTTATCAGAGGCACGTCAACATCTTAAAGATGGCACTTGTGGCTTAGTAGAAGTTGAAAAAGGCGTTTTGCCTCAACTTGAACAGCCCTATGTGTTCATCAAACGTTCGGATGCTCGAACTGCACCTCATGGTCATGTTATGGTTGAGCTGGTAGCAGAACTCGAAGGCATTCAGTACGGTCGTAGTGGTGAGACACTTGGTGTCCTTGTCCCTCATGTGGGCGAAATACCAGTGGCTTACCGCAAGGTTCTTCTTCGTAAGAACGGTAATAAAGGAGCTGGTGGCCATAGTTACGGCGCCGATCTAAAGTCATTTGACTTAGGCGACGAGCTTGGCACTGATCCTTATGAAGATTTTCAAGAAAACTGGAACACTAAACATAGCAGTGGTGTTACCCGTGAACTCATGCGTGAGCTTAACGGAGGGGCATACACTCGCTATGTCGATAACAACTTCTGTGGCCCTGATGGCTACCCTCTTGAGTGCATTAAAGACCTTCTAGCACGTGCTGGTAAAGCTTCATGCACTTTGTCCGAACAACTGGACTTTATTGACACTAAGAGGGGTGTATACTGCTGCCGTGAACATGAGCATGAAATTGCTTGGTACACGGAACGTTCTGGGCCCTCGA

### 1-1000 A70U

GGGTTAAAGGTTTATACCTTCCCAGGTAACAAACCAACCAACTTTCGATCTCTTGTAGATCTGTTCTCTAATCGAACTTTAAAATCTGTGTGGCTGTCACTCGGCTGCATGCTTAGTGCACTCACGCAGTATAATTAATAACTAATTACTGTCGTTGACAGGACACGAGTAACTCGTCTATCTTCTGCAGGCTGCTTACGGTTTCGTCCGTGTTGCAGCCGATCATCAGCACATCTAGGTTTCGTCCGGGTGTGACCGAAAGGTAAGATGGAGAGCCTTGTCCCTGGTTTCAACGAGAAAACACACGTCCAACTCAGTTTGCCTGTTTTACAGGTTCGCGACGTGCTCGTACGTGGCTTTGGAGACTCCGTGGAGGAGGTCTTATCAGAGGCACGTCAACATCTTAAAGATGGCACTTGTGGCTTAGTAGAAGTTGAAAAAGGCGTTTTGCCTCAACTTGAACAGCCCTATGTGTTCATCAAACGTTCGGATGCTCGAACTGCACCTCATGGTCATGTTATGGTTGAGCTGGTAGCAGAACTCGAAGGCATTCAGTACGGTCGTAGTGGTGAGACACTTGGTGTCCTTGTCCCTCATGTGGGCGAAATACCAGTGGCTTACCGCAAGGTTCTTCTTCGTAAGAACGGTAATAAAGGAGCTGGTGGCCATAGTTACGGCGCCGATCTAAAGTCATTTGACTTAGGCGACGAGCTTGGCACTGATCCTTATGAAGATTTTCAAGAAAACTGGAACACTAAACATAGCAGTGGTGTTACCCGTGAACTCATGCGTGAGCTTAACGGAGGGGCATACACTCGCTATGTCGATAACAACTTCTGTGGCCCTGATGGCTACCCTCTTGAGTGCATTAAAGACCTTCTAGCACGTGCTGGTAAAGCTTCATGCACTTTGTCCGAACAACTGGACTTTATTGACACTAAGAGGGGTGTATACTGCTGCCGTGAACATGAGCATGAAATTGCTTGGTACACGGAACGTTCTGGGCCCTCGA

### 1-1000 Rare Y

GGGTTAAAGGTTTATACCTTCCCAGGTAACAAACCAACCAACTTTCGATCTCTTGTAGATCTGTTCTCCAAACGAACTTTAAAATCTGTGTGGCTGTCACTCGGCTGCATGCTTAGTGCACTCACGCAGTATAATTAATAACTAATTACTGTCGTTGACAGGACACGAGTAACTCGTCTATCTTCTGCAGGCTGCTTACGGTTTCGTCCGTGTTGCAGCCGATCATCAGCACATCTAGGTTTCGTCCGGGTGTGACCGAAAGGTAAGATGGAGAGCCTTGTCCCTGGTTTCAACGAGAAAACACACGTCCAACTCAGTTTGCCTGTTTTACAGGTTCGCGACGTGCTCGTACGTGGCTTTGGAGACTCCGTGGAGGAGGTCTTATCAGAGGCACGTCAACATCTTAAAGATGGCACTTGTGGCTTAGTAGAAGTTGAAAAAGGCGTTTTGCCTCAACTTGAACAGCCCTATGTGTTCATCAAACGTTCGGATGCTCGAACTGCACCTCATGGTCATGTTATGGTTGAGCTGGTAGCAGAACTCGAAGGCATTCAGTACGGTCGTAGTGGTGAGACACTTGGTGTCCTTGTCCCTCATGTGGGCGAAATACCAGTGGCTTACCGCAAGGTTCTTCTTCGTAAGAACGGTAATAAAGGAGCTGGTGGCCATAGTTACGGCGCCGATCTAAAGTCATTTGACTTAGGCGACGAGCTTGGCACTGATCCTTATGAAGATTTTCAAGAAAACTGGAACACTAAACATAGCAGTGGTGTTACCCGTGAACTCATGCGTGAGCTTAACGGAGGGGCATACACTCGCTATGTCGATAACAACTTCTGTGGCCCTGATGGCTACCCTCTTGAGTGCATTAAAGACCTTCTAGCACGTGCTGGTAAAGCTTCATGCACTTTGTCCGAACAACTGGACTTTATTGACACTAAGAGGGGTGTATACTGCTGCCGTGAACATGAGCATGAAATTGCTTGGTACACGGAACGTTCTGGGCCCTCGA

### 1-1000 Common Y

GGGTTAAAGGTTTATACCTTCCCAGGTAACAAACCAACCAACTTTCGATCTCTTGTAGATCTGTTCTTTAAATGAACTTTAAAATCTGTGTGGCTGTCACTCGGCTGCATGCTTAGTGCACTCACGCAGTATAATTAATAACTAATTACTGTCGTTGACAGGACACGAGTAACTCGTCTATCTTCTGCAGGCTGCTTACGGTTTCGTCCGTGTTGCAGCCGATCATCAGCACATCTAGGTTTCGTCCGGGTGTGACCGAAAGGTAAGATGGAGAGCCTTGTCCCTGGTTTCAACGAGAAAACACACGTCCAACTCAGTTTGCCTGTTTTACAGGTTCGCGACGTGCTCGTACGTGGCTTTGGAGACTCCGTGGAGGAGGTCTTATCAGAGGCACGTCAACATCTTAAAGATGGCACTTGTGGCTTAGTAGAAGTTGAAAAAGGCGTTTTGCCTCAACTTGAACAGCCCTATGTGTTCATCAAACGTTCGGATGCTCGAACTGCACCTCATGGTCATGTTATGGTTGAGCTGGTAGCAGAACTCGAAGGCATTCAGTACGGTCGTAGTGGTGAGACACTTGGTGTCCTTGTCCCTCATGTGGGCGAAATACCAGTGGCTTACCGCAAGGTTCTTCTTCGTAAGAACGGTAATAAAGGAGCTGGTGGCCATAGTTACGGCGCCGATCTAAAGTCATTTGACTTAGGCGACGAGCTTGGCACTGATCCTTATGAAGATTTTCAAGAAAACTGGAACACTAAACATAGCAGTGGTGTTACCCGTGAACTCATGCGTGAGCTTAACGGAGGGGCATACACTCGCTATGTCGATAACAACTTCTGTGGCCCTGATGGCTACCCTCTTGAGTGCATTAAAGACCTTCTAGCACGTGCTGGTAAAGCTTCATGCACTTTGTCCGAACAACTGGACTTTATTGACACTAAGAGGGGTGTATACTGCTGCCGTGAACATGAGCATGAAATTGCTTGGTACACGGAACGTTCTGGGCCCTCGA

### 1-1000 Rescue WT

GGGTTAAAGGTTTATACCTTCCCAGGTAACAAACCAACCAACTTTCGATCTCTTGTAGATCTGTTCTCTAAACGAACTTTAAAATCTGTGTGGCTGTCACTCGGCTGCATGCTTAGTGCACTCACGCAGTATAATTAATAACTAATTACTGTCGTTCCCAGGACACGAGTAACTCGTCTATCTTCTGCAGGCTGCTTACGGTTTCGTCCGTGTTGCAGCCGATCATCAGCACATCTAGGTTTCGTCCGGGTGTGACCGAAAGGTAAGATGGAGAGCCTTGTCCCTGGTTTCAACGAGAAAACACACGTCCAACTCAGTTTGCCTGTTTTACAGGTTCGCGACGTGCTCGTACGTGGCTTTGGAGACTCCGTGGAGGAGGTCTTATCAGAGGCACGTCAACATCTTAAAGATGGCACTTGTGGCTTAGTAGAAGTTGAAAAAGGCGTTTTGCCTCAACTTGAACAGCCCTATGTGTTCATCAAACGTTCGGATGCTCGAACTGCACCTCATGGTCATGTTATGGTTGAGCTGGTAGCAGAACTCGAAGGCATTCAGTACGGTCGTAGTGGTGAGACACTTGGTGTCCTTGTCCCTCATGTGGGCGAAATACCAGTGGCTTACCGCAAGGTTCTTCTTCGTAAGAACGGTAATAAAGGAGCTGGTGGCCATAGTTACGGCGCCGATCTAAAGTCATTTGACTTAGGCGACGAGCTTGGCACTGATCCTTATGAAGATTTTCAAGAAAACTGGAACACTAAACATAGCAGTGGTGTTACCCGTGAACTCATGCGTGAGCTTAACGGAGGGGCATACACTCGCTATGTCGATAACAACTTCTGTGGCCCTGATGGCTACCCTCTTGAGTGCATTAAAGACCTTCTAGCACGTGCTGGTAAAGCTTCATGCACTTTGTCCGAACAACTGGACTTTATTGACACTAAGAGGGGTGTATACTGCTGCCGTGAACATGAGCATGAAATTGCTTGGTACACGGAACGTTCTGGGCCCTCGA

### 1-1000 Rescue A68U

GGGTTAAAGGTTTATACCTTCCCAGGTAACAAACCAACCAACTTTCGATCTCTTGTAGATCTGTTCTCTATACGAACTTTAAAATCTGTGTGGCTGTCACTCGGCTGCATGCTTAGTGCACTCACGCAGTATAATTAATAACTAATTACTGTCGTTCCCAGGACACGAGTAACTCGTCTATCTTCTGCAGGCTGCTTACGGTTTCGTCCGTGTTGCAGCCGATCATCAGCACATCTAGGTTTCGTCCGGGTGTGACCGAAAGGTAAGATGGAGAGCCTTGTCCCTGGTTTCAACGAGAAAACACACGTCCAACTCAGTTTGCCTGTTTTACAGGTTCGCGACGTGCTCGTACGTGGCTTTGGAGACTCCGTGGAGGAGGTCTTATCAGAGGCACGTCAACATCTTAAAGATGGCACTTGTGGCTTAGTAGAAGTTGAAAAAGGCGTTTTGCCTCAACTTGAACAGCCCTATGTGTTCATCAAACGTTCGGATGCTCGAACTGCACCTCATGGTCATGTTATGGTTGAGCTGGTAGCAGAACTCGAAGGCATTCAGTACGGTCGTAGTGGTGAGACACTTGGTGTCCTTGTCCCTCATGTGGGCGAAATACCAGTGGCTTACCGCAAGGTTCTTCTTCGTAAGAACGGTAATAAAGGAGCTGGTGGCCATAGTTACGGCGCCGATCTAAAGTCATTTGACTTAGGCGACGAGCTTGGCACTGATCCTTATGAAGATTTTCAAGAAAACTGGAACACTAAACATAGCAGTGGTGTTACCCGTGAACTCATGCGTGAGCTTAACGGAGGGGCATACACTCGCTATGTCGATAACAACTTCTGTGGCCCTGATGGCTACCCTCTTGAGTGCATTAAAGACCTTCTAGCACGTGCTGGTAAAGCTTCATGCACTTTGTCCGAACAACTGGACTTTATTGACACTAAGAGGGGTGTATACTGCTGCCGTGAACATGAGCATGAAATTGCTTGGTACACGGAACGTTCTGGGCCCTCGA

### 1-1000 Rescue A69U

GGGTTAAAGGTTTATACCTTCCCAGGTAACAAACCAACCAACTTTCGATCTCTTGTAGATCTGTTCTCTATACGAACTTTAAAATCTGTGTGGCTGTCACTCGGCTGCATGCTTAGTGCACTCACGCAGTATAATTAATAACTAATTACTGTCGTTCCCAGGACACGAGTAACTCGTCTATCTTCTGCAGGCTGCTTACGGTTTCGTCCGTGTTGCAGCCGATCATCAGCACATCTAGGTTTCGTCCGGGTGTGACCGAAAGGTAAGATGGAGAGCCTTGTCCCTGGTTTCAACGAGAAAACACACGTCCAACTCAGTTTGCCTGTTTTACAGGTTCGCGACGTGCTCGTACGTGGCTTTGGAGACTCCGTGGAGGAGGTCTTATCAGAGGCACGTCAACATCTTAAAGATGGCACTTGTGGCTTAGTAGAAGTTGAAAAAGGCGTTTTGCCTCAACTTGAACAGCCCTATGTGTTCATCAAACGTTCGGATGCTCGAACTGCACCTCATGGTCATGTTATGGTTGAGCTGGTAGCAGAACTCGAAGGCATTCAGTACGGTCGTAGTGGTGAGACACTTGGTGTCCTTGTCCCTCATGTGGGCGAAATACCAGTGGCTTACCGCAAGGTTCTTCTTCGTAAGAACGGTAATAAAGGAGCTGGTGGCCATAGTTACGGCGCCGATCTAAAGTCATTTGACTTAGGCGACGAGCTTGGCACTGATCCTTATGAAGATTTTCAAGAAAACTGGAACACTAAACATAGCAGTGGTGTTACCCGTGAACTCATGCGTGAGCTTAACGGAGGGGCATACACTCGCTATGTCGATAACAACTTCTGTGGCCCTGATGGCTACCCTCTTGAGTGCATTAAAGACCTTCTAGCACGTGCTGGTAAAGCTTCATGCACTTTGTCCGAACAACTGGACTTTATTGACACTAAGAGGGGTGTATACTGCTGCCGTGAACATGAGCATGAAATTGCTTGGTACACGGAACGTTCTGGGCCCTCGA

### Frameshifting-region (FS) RNA

GGGAGAGCGGCCGCCAGATCTTCCGGATGGCTCGAGTTTTTCAGCAAGATTGGCTGTAGTTGTGATCAACTCCGCGAACCCATGCTTCAGTCAGCTGATGCACAATCGTTTTTAAACGGGTTTGCGGTGTAAGTGCAGCCCGTCTTACACCGTGCGGCACAGGCACTAGTACTGATGTCGTATACAGGGCTTTTGACATCTACAATGATAAAGTAGCTGGTTTTGCTAAATTCCTAAAAACTAATTGTTGTCGCTTCCAAGAAAAGGACGAAGATGACAATTTAATTGATTCTTACTTTGTAGTTAAGAGACACACTTTCTCTAACTACCAACATGAAGAAACAATTTATAATTTACTTAAGGATTGTCCAGCTGTTGCTAAACATGACTTCTTTAAGTTTAGAATAGACGGTGACATGGTACCACATATATCACGTCAACGTCTTACTAAATACACAATGGCAGACCTCGTCTATGCTTTAAGGCATTTTGATGAAGGTAATTGTGACACATTAAAAGAAATACTTGTCACATACAATTGTTGTGATGATGATTATTTCAATAAAAAGGACTGGTATGATTTTGTAGAAAACCCAGATATATTACGCGTATACGCCAACTTAGGTGAACGTGTACGCCAAGCTTTGTTAAAAACAGTACAATTCTGTGATGCCATGCGAAATGCTGGTATTGTTGGTGTACTGACATTAGATAATCAAGATCTCAATGGTAACTGGTATGATTTCGGTGATTTCATACAAACCACGCCAGGTAGTGGAGTTCCTGTTGTAGATTCTTATTATTCATTGTTAATGCCTATATTAACCTTGACCAGGGCTTTAACTGCAGAGTCACATGTTGACACTGACTTAACAAAGCCTTACATTAAGTGGGATTTGTTAAAATATGACTTCACGGAAGAGAGGTTAAAACTCTTTGACCGTTATTTTAAATATTGGGATCAGACATACCACCCAAATTGTGTTAACTGTTTGGATGACAGATGCATTCTGCATTGTGCAAACTTTAATGTTTTATTCTCTACAGTGTATCTTT

### Nucleocapsid RNA

GATTAAAGGTTTATACCTTCCCAGGTAACAAACCAACCAACTTTCGATCTCTTGTAGATCTGTTCTCTAAACGAACAAACTAAAATGTCTGATAATGGACCCCAAAATCAGCGAAATGCACCCCGCATTACGTTTGGTGGACCCTCAGATTCAACTGGCAGTAACCAGAATGGAGAACGCAGTGGGGCGCGATCAAAACAACGTCGGCCCCAAGGTTTACCCAATAATACTGCGTCTTGGTTCACCGCTCTCACTCAACATGGCAAGGAAGACCTTAAATTCCCTCGAGGACAAGGCGTTCCAATTAACACCAATAGCAGTCCAGATGACCAAATTGGCTACTACCGAAGAGCTACCAGACGAATTCGTGGTGGTGACGGTAAAATGAAAGATCTCAGTCCAAGATGGTATTTCTACTACCTAGGAACTGGGCCAGAAGCTGGACTTCCCTATGGTGCTAACAAAGACGGCATCATATGGGTTGCAACTGAGGGAGCCTTGAATACACCAAAAGATCACATTGGCACCCGCAATCCTGCTAACAATGCTGCAATCGTGCTACAACTTCCTCAAGGAACAACATTGCCAAAAGGCTTCTACGCAGAAGGGAGCAGAGGCGGCAGTCAAGCCTCTTCTCGTTCCTCATCACGTAGTCGCAACAGTTCAAGAAATTCAACTCCAGGCAGCAGTAGGGGAACTTCTCCTGCTAGAATGGCTGGCAATGGCGGTGATGCTGCTCTTGCTTTGCTGCTGCTTGACAGATTGAACCAGCTTGAGAGCAAAATGTCTGGTAAAGGCCAACAACAACAAGGCCAAACTGTCACTAAGAAATCTGCTGCTGAGGCTTCTAAGAAGCCTCGGCAAAAACGTACTGCCACTAAAGCATACAATGTAACACAAGCTTTCGGCAGACGTGGTCCAGAACAAACCCAAGGAAATTTTGGGGACCAGGAACTAATCAGACAAGGAACTGATTACAAACATTGGCCGCAAATTGCACAATTTGCCCCCAGCGCTTCAGCGTTCTTCGGAATGTCGCGCATTGGCATGGAAGTCACACCTTCGGGAACGTGGTTGACCTACACAGGTGCCATCAAATTGGATGACAAAGATCCAAATTTCAAAGATCAAGTCATTTTGCTGAATAAGCATATTGACGCATACAAAACATTCCCACCAACAGAGCCTAAAAAGGACAAAAAGAAGAAGGCTGATGAAACTCAAGCCTTACCGCAGAGACAGAAGAAACAGCAAACTGTGACTCTTCTTCCTGCTGCAGATTTGGATGATTTCTCCAAACAATTGCAACAATCCATGAGCAGTGCTGACTCAACTCAGG

### 5′UTR Wildtype

GGGTTAAAGGTTTATACCTTCCCAGGTAACAAACCAACCAACTTTCGATCTCTTGTAGATCTGTTCTCTAAACGAACTTTAAAATCTGTGTGGCTGTCACTCGGCTGCATGCTTAGTGCACTCACGCAGTATAATTAATAACTAATTACTGTCGTTGACAGGACACGAGTAACTCGTCTATCTTCTGCAGGCTGCTTACGGTTTCGTCCGTGTTGCAGCCGATCATCAGCACATCTAGGTTTCGTCCGGGTGTGACCGAAAGGTAAG

### 5′UTR +dsRNAb

GGGTTAAAGGTTTATACCTTCCCAGGTAACAAACCAACCAACTTTCGATCTCTTGTAGATCTGTTCTCTAAACGAACTTTAAAATCAGTAATTAGTTATTAATTATACTGTGTGGCTGTCACTCGGCTGCATGCTTAGTGCACTCACGCAGTATAATTAATAACTAATTACTGTCGTTGACAGGACACGAGTAACTCGTCTATCTTCTGCAGGCTGCTTACGGTTTCGTCCGTGTTGCAGCCGATCATCAGCACATCTAGGTTTCGTCCGGGTGTGACCGAAAGGTAAG

### 5′UTR +dsRNAd

GGGTTAAAGGTTTATACCTTCCCAGGTAACAAACCAACCAACTTTCGATCTCTTGTAGATCTGTTCTCTAAACGAACTTTAAAATCGATTGCATCCTGTGTGGCTGTCACTCGGCTGCATGCTTAGTGCACTCACGCAGGATGCAATCGTATAATTAATAACTAATTACTGCGATTGCATCTCGTTGACAGGACACGAGTAACTCGTCTATCTTCTGCAGGCTGCTTACGGTTTCGTCCGTGTTGCAGCCGATCATCAGCACATCTAGGTTTCGTCCGGGTGTGACCGAAAGGTAAG

### 5′UTR +ssRNAa

GGGTTAAAGGTTTATACCTTCCCAGGTAACAAACCAACCAACTTTCGATCTCTTGTAGATCTGTTCTCTAAACGAACTTTAAAATCTGTGTGGCTGTCACTCGGCTGCATGCTTAGTGCACTCACGCAGTATAATTAATATACCCATACGATGTTCCAGATTACGCTACTAATTACTGTCGTTGACAGGACACGAGTAACTCGTCTATCTTCTGCAGGCTGCTTACGGTTTCGTCCGTGTTGCAGCCGATCATCAGCACATCTAGGTTTCGTCCGGGTGTGACCGAAAGGTAAG

### 5′UTR -dsRNA

GGGTTAAAGGTTTATACCTTCCCAGGTAACAAACCAACCAACTTTCGATCTCTTGTAGATCTGTTCTCTAAACGAACTTTAAAATGGTGTGTTCTGTCCAGCGTCTGCATGCTTAGTGCACTCACGCAGTATAATTAATAACTAATTACTGTATGGTGGCTTACGATCATAAAGTGCAGCGAGCCTGCAGGCTGCTTACGGTTTCGTCCGTGTTGCAGCCGATCATCAGCACATCTAGGTTTCGTCCGGGTGTGACCGAAAGGTAAG

### 5′UTR Restore pairing

GGGTTAAAGGTTTATACCTTCCCAGGTAACAAACCAACCAACTTTCGATCTCTTGTAGATCTGTTCTCTAAACGAACTTTAAAATGGTGTGTTCTGTCCAGCGTCTGCATGATTCTGGCACGACATACCTATAATTAATAACTAATTACTGTATGGTGGCTTACGATCATAAAGTGCAGCGAGCCTGCAGGCTGCTTACGGTTTCGTCCGTGTTGCAGCCGATCATCAGCACATCTAGGTTTCGTCCGGGTGTGACCGAAAGGTAAG

### Wildtype 5′UTR: Nano Luciferase

GGGTTAAAGGTTTATACCTTCCCAGGTAACAAACCAACCAACTTTCGATCTCTTGTAGATCTGTTCTCTAAACGAACTTTAAAATCTGTGTGGCTGTCACTCGGCTGCATGCTTAGTGCACTCACGCAGTATAATTAATAACTAATTACTGTCGTTGACAGGACACGAGTAACTCGTCTATCTTCTGCAGGCTGCTTACGGTTTCGTCCGTGTTGCAGCCGATCATCAGCACATCTAGGTTTCGTCCGGGTGTGACCGAAAGGTAAGATGGAGAGCCTTGTCCCTGGTTTCAACGACTCGAGTGGCTCGGGCTCGACCTCGGGCTCGGGCAAAACCGGTGTCTTCACACTCGAAGATTTCGTTGGGGACTGGCGACAGACAGCCGGCTACAACCTGGACCAAGTCCTTGAACAGGGAGGTGTGTCCAGTTTGTTTCAGAATCTCGGGGTGTCCGTAACTCCGATCCAAAGGATTGTCCTGAGCGGTGAAAATGGGCTGAAGATCGACATCCATGTCATCATCCCGTATGAAGGTCTGAGCGGCGACCAAATGGGCCAGATCGAAAAAATTTTTAAGGTGGTGTACCCTGTGGATGATCATCACTTTAAGGTGATCCTGCACTATGGCACACTGGTAATCGACGGGGTTACGCCGAACATGATCGACTATTTCGGACGGCCGTATGAAGGCATCGCCGTGTTCGACGGCAAAAAGATCACTGTAACAGGGACCCTGTGGAACGGCAACAAAATTATCGACGAGCGCCTGATCAACCCCGACGGCTCCCTGCTGTTCCGAGTAACCATCAACGGAGTGACCGGCTGGCGGCTGTGCGAACGCATTCTGGCGTAAGGGCCCGCGGTTCGAAGGTAAGCCTATCCC

### 5′UTR +dsRNAb Nano Luciferase

GGGTTAAAGGTTTATACCTTCCCAGGTAACAAACCAACCAACTTTCGATCTCTTGTAGATCTGTTCTCTAAACGAACTTTAAAATCAGTAATTAGTTATTAATTATACTGTGTGGCTGTCACTCGGCTGCATGCTTAGTGCACTCACGCAGTATAATTAATAACTAATTACTGTCGTTGACAGGACACGAGTAACTCGTCTATCTTCTGCAGGCTGCTTACGGTTTCGTCCGTGTTGCAGCCGATCATCAGCACATCTAGGTTTCGTCCGGGTGTGACCGAAAGGTAAGATGGAGAGCCTTGTCCCTGGTTTCAACGACTCGAGTGGCTCGGGCTCGACCTCGGGCTCGGGCAAAACCGGTGTCTTCACACTCGAAGATTTCGTTGGGGACTGGCGACAGACAGCCGGCTACAACCTGGACCAAGTCCTTGAACAGGGAGGTGTGTCCAGTTTGTTTCAGAATCTCGGGGTGTCCGTAACTCCGATCCAAAGGATTGTCCTGAGCGGTGAAAATGGGCTGAAGATCGACATCCATGTCATCATCCCGTATGAAGGTCTGAGCGGCGACCAAATGGGCCAGATCGAAAAAATTTTTAAGGTGGTGTACCCTGTGGATGATCATCACTTTAAGGTGATCCTGCACTATGGCACACTGGTAATCGACGGGGTTACGCCGAACATGATCGACTATTTCGGACGGCCGTATGAAGGCATCGCCGTGTTCGACGGCAAAAAGATCACTGTAACAGGGACCCTGTGGAACGGCAACAAAATTATCGACGAGCGCCTGATCAACCCCGACGGCTCCCTGCTGTTCCGAGTAACCATCAACGGAGTGACCGGCTGGCGGCTGTGCGAACGCATTCTGGCGTAAGGGCCCGCGGTTCGAAGGTAAGCCTATCCC

#### In vitro transcription

was carried out according to our established protocols (Langdon et al., 2018). Orf1ab templates were synthesized (IDT) and cloned into pJet (ThermoFisher Scientific K1231) using blunt end cloning. Directionality and sequence were confirmed using Sanger sequencing (GENEWIZ). Plasmid were linearized using PCR (iProof Bio-Rad 1725310). 5 μl of PCR product was loaded onto an agarose gel to determine size and purity. If the PCR product was pure then the sample was PCR purified (QIAGEN 28106) if the band was impure, it was gel purified (QIAGEN 28706) (PCR impurity was most often was a problem for the ultrastructured mutants of principal site 2). 100 ng of gel or PCR purified DNA was used as a template for in vitro transcription (NEB E2040S) carried out according to the manufacturer’s instructions with the addition of 0.1μl of Cy3 (Sigma PA53026) or Cy5 (Sigma PA55026) labeled UTP to each reaction. Following incubation at 37°C for 18 hours, in vitro transcription reactions were treated with DNAseI (NEB M0303L) according to the manufacturer’s instructions. Following DNAse treatment, reactions were purified with 2.5M LiCl precipitation. Purified RNA amounts were quantified using nanodrop and verified for purity and size using a denaturing agarose gel and Riboruler RNA ladder (Thermo Scientific SM183).

#### Phase separation assays

For in vitro reconstitution LLPS experiments, 15 μl droplet buffer (20 mM Tris pH 7.5, 150 mM NaCl) was mixed with cy3 or cy5 labeled desired RNA and DEPC treated H20 (final volume 5 μl) and 5 μl protein in storage buffer was added at desired concentration. The mix was incubated in 384-well plates (Cellvis P384-1.5H-N) for 1-20 hours at 37°C unless indicated otherwise. Droplets formed after short incubations of 20 minutes or less, however, they were initially smaller and matured into larger droplets during the overnight incubation step. Time to maturation varied based on the ratio of RNA to protein, concentration of RNA and protein and RNA sequence. Multiple conditions per mutant were tested with the most optimal conditions for differences selected for comparison. Imaging of droplets was done on a spinning disc confocal microscope (Nikon CSU-W1) with VC Plan Apo 100X/1.49 NA oil (Cargille Lab 16241) immersion objective and an sCMOS 85% QE 95B camera (Photometrics). Data shown are representative of three or more independent replicates, across 2 or more RNA preparations. Whenever possible multiple mutations were designed to disrupt the same class of feature in multiple sequence contexts.

#### EMSA

65ng/μl of the indicated RNA sequence was incubated with 0, 0.75, 1.5, or 2.2 μM Y109A mutant N-protein at 25 or 37°C for 1 hour in the following buffer 10mM HEPES pH 7.5, 50μM EDTA, 10% glycerol, 1mM DTT, 5mM MgCl2, 0.1mg/ml BSA, 2.5ug Yeast TRNA, 10U RNAse inhibitor and loading dye. Samples were then loaded onto an 8% TBE gel and run at 100V for 1 hour at 4°C. Gels were then stained with SYBRgold (S11494) and imaged. Unbound RNA was quantified using ImageJ.

#### Temperature dependent turbidity tests

The LCST behaviors of different phase separation systems were investigated on a Cary 300 temperature-dependent ultraviolet-visible spectroscopy equipped with a multicell thermoelectric temperature controller. The samples (4 μM of N-protein with 15 nM of RNAs) were mixed and prepared in a droplet buffer (20 mM Tris pH 7.5, 150 mM NaCl) at 4 °C. Before the initiation of the heating process of the turbidity test, for the experiments shown in Fig. 3A, the samples were incubated for 1 hour at 4 °C; for the experiments shown in Fig. 3B**&**C, the samples were incubated for 20 min at 4 °C. A heating rate of 1°C/min was applied during the temperature ramp while the absorbance at λ = 350 nm was recorded at every 0.33 °C increment. Normalized turbidity was calculated by the absorbance at the lowest temperature point normalizes to the absorbance at the highest temperature point.

#### Comparison of droplet images to absorbance A280 reading in diffuse phase

The mix was incubated in 384-well plates (Cellvis P384-1.5H-N) at 25, 30, or 37°C. Following imaging. 2 μl of diffuse phase solution (taken from the top of the well) was nanodropped and absorbance A280 was recorded. Error bars indicate the A280 measurement from the 3 technical replicates. (Of note, concentrations below 3μM N-protein did not give high enough A280 absorbance to generate reliable measurements.)

#### Phyre Structure prediction/ Pymol structure alignment

The following SARS-CoV-2 amino acid sequence was input into Phyre

TKKSAAEASKKPRQKRTATKAYNVTQAFGRRGPEQTQGNFGDQELIRQGTDYKHWPQIAQFAPSASAFFGMSRIGMEVTPSGTWLTYTGAIKLDDKDPNFKDQVILLNKHIDAYKTFP.

This sequence best matched with the crystal structure of the RBD2-dimerization domain of SARS-CoV-1 (Chen et al., 2007). The resulting structure prediction was aligned to the crystal structure of SARS-CoV-1 or MERS-CoV (Nguyen et al., 2019) using Pymol.

#### Mass photometry of purified N-protein

Mass photometry was performed according to established protocols (Sonn-Segev et al., 2020). 10μL of protein storage buffer (250mM NaCl 20mM phosphate buffer pH 7.5) was used to focus followed by addition of 10μl of 40nM N-protein in protein storage buffer (wildtype or RBD2-del) for a final protein concentration of 20nM. Representative histograms from were generated from 2 minutes movies reflective of the raw detected particle molecular weight in kDAs.

#### In vitro translation assay

Protocol was adapted from the method described by Tsang et al. (Tsang et al., 2019) Briefly, 25nM of 5′UTR nano luciferase fusion RNA was incubated with either protein 0.3μM or 3.2μM N-protein for 20-minutes at room temperature in PCR strip tubes (8ul total volume, final buffer conditions 140mM NaCl, 4mM phosphate buffer, 12mM TRIS pH 7.5) as a control for basal luciferase RNA translation, N-protein storage buffer was added (250mM NaCl 20mM phosphate buffer pH 7.5). Following incubation, 5ul rabbit reticulocyte lysate +Met +Leu (Promega L4960), was added to the protein/RNA mixture (or RNA and buffer) and the resulting mix was incubated at 30°C for 2 hours. 2μl of in vitro translation product was then mixed with 25μl of nano luciferase assay reagents (Promega N205A). Light production was measured on a luminometer. Data depicted represents at least three replicates. Of note, similar translational repression was observed when we incubated RNA under droplet permissive conditions in plates (37°C 1-2 hours) however this was much less reproducible likely due to the difference in RNA partitioning in the well post incubation with N-protein.

#### Cell Culture

HEK293T, and Vero-E6 cells were originally obtained from ATCC. All cell lines were maintained in DMEM (Corning 10-013-CV) supplemented with 10% Fetal Bovine Serum (Gibco). No antibiotics were used.

#### Plasmid Transfection

24 hours prior to transfection, confluent cells were split 1:5. Two hours prior to transfection, 500μl of fresh media was added to 24 well plates. 500ng of plasmid DNA for each Nucleocapsid GFP Spark (Sino biological VG40588-ACGLN) and the MSCV blast 1-1000 fragments was co-transfected using FUGENE HD. Transfections were then incubated for 24-48 hours prior to imaging.

#### Cell Imaging

Cells were imaged using a 40X air objective on a spinning disk confocal microscope (Nikon Ti-Eclipse, Yokogawa CSU-X1 spinning disk). Images were taken with a ANDOR camera. Representative cells are taken from 3 biological replicates.

#### Cell Imaging Quantification

Cells with puncta were cropped using FIJI. Experimenters were then blinded to conditions, and puncta were counted for each cell. Whole cell N:GFP signal was quantified using ImageTank (O’Shaughnessy et al., 2019).

#### Transmission electron microscopy (TEM) and quantification of RNP size distribution

For negative stained TEM images used to quantify the assemblies of RNP size distribution, 5ul of 20μM protein in 250mM NaCl, 20mM phosphate buffer pH 7.5 and 5ul of 50nM RNA (FS RNA or 1000) in water were mixed in 15ul of reaction buffer (150mM NaCl, 20mM Tris, pH 7.5). The final protein and RNA concentrations in the solution were 4μM and 10nM, respectively. For control measurements, protein without RNA and RNA without protein solutions were prepared. All mixture solutions were incubated at room temperature for overnight to measure negative stained TEM images.

Negatively stained samples were prepared on carbon film-coated grids supported by lacey carbon on 300 copper mesh (Electron microscopy Sciences). Grids were glow-discharged immediately before use. 8μl aliquot of protein and RNA mixture solution was applied to the grid. After 2 min absorption to the carbon film, the solution was blotted and washed with 8μl of water for 10 s, blotted, stained with 8μl of 2 % uranyl acetate for 10 s, blotted, and dried. Negative stained TEM images were obtained on a FEI Morgagni microscope.

Images were analyzed with ImageJ software (available at http://imagej.nih.gov/ij). Since the shape of the small RNP is not a sphere, two major and minor diameters of the elliptical shape of the RNP were measured, and the averaged values from two diameters were reported. Gaussian fitting of averaged diameter histogram was performed with Igor Pro 8.0.4.2 (WaveMetrics).

#### RNP-MaP probing of N-Protein-RNA interactions

N-Protein and RNA mixtures were prepared as described in the “Phase Separation Assay” section above and incubated for 1.5 hours at 37°C. N-protein or N-protein Y109A– FS RNA mixtures were prepared in 50nM RNA, 1μM protein (diffuse state, 20x excess protein) RNA-only samples were also prepared as a control. After confirmation of phase separation by imaging mixtures were immediately subjected to RNP-MaP treatment as described (Weidmann et al., 2021), with modifications described below. Briefly, 200 μl of mixtures were added to 10.5 μl of 200 mM SDA (in DMSO) in wells of a 6-well plate and incubated in the dark for 10 minutes at 37°C. RNPs were crosslinked with 3 J/cm^2^ of 365 nm wavelength UV light. To digest unbound and crosslinked N-proteins, reactions were adjusted to 1.5% SDS, 20 mM EDTA, 200mM NaCl, and 40mM Tris-HCl (pH 8.0) and incubated at 37°C for 10 minutes, heated to 95°C for 5 minutes, cooled on ice for 2 minutes, and warmed to 37°C for 2 minutes. Proteinase K was then added to 0.5 mg/ml and incubated for 1 hour at 37°C, followed by 1 hour at 55°C. RNA was purified with 1.8 ′ Mag-Bind TotalPure NGS SPRI beads (Omega Bio-tek), purified again (RNeasy MinElute columns, Qiagen), and eluted with 14 μl of nuclease-free water.

#### MaP reverse transcription

After SHAPE and RNP-MaP RNA modification and purification, MaP cDNA synthesis was performed using a revised protocol as described (Mustoe et al., 2019). Briefly, 7 μl of purified modified RNA was mixed with 200 ng of random 9-mer primers and 20 nmol of dNTPs and incubated at 65°C for 10 min followed by 4°C for 2 min. 9 μl 2.22 ′ MaP buffer [1 ′ MaP buffer consists of 6 mM MnCl2, 1 M betaine, 50 mM Tris (pH 8.0), 75 mM KCl, 10 mM DTT] was added and the combined solution was incubated at 23°C for 2 min. 1 μl Superscript II Reverse Transcriptase (200 units, Invitrogen) was added and the reverse transcription (RT) reaction was performed according to the following temperature program: 25°C for 10 min, 42°C for 90 min, 10 ′[50°C for 2 min, 42°C for 2 min], 72°C for 10 min. RT cDNA products were then purified (Illustra G-50 microspin columns, GE Healthcare).

#### Library preparation and Sequencing

Double-stranded DNA (dsDNA) libraries for sequencing were prepared using the randomer Nextera workflow (Smola et al., 2015). Briefly, purified cDNA was added to an NEBNext second-strand synthesis reaction (NEB) at 16°C for 150 minutes. dsDNA products were purified and size-selected with SPRI beads at a 0.8 ′ ratio. Nextera XT (Illumina) was used to construct libraries according to the manufacturer’s protocol, followed by purification and size-selection with SPRI beads at a 0.65 ′ ratio. Library size distributions and purities were verified (2100 Bioanalyzer, Agilent) and sequenced using 2x300 paired-end sequencing on an Illumina MiSeq instrument (v3 chemistry).

#### Sequence alignment and mutation parsing

FASTQ files from sequencing runs were directly input into *ShapeMapper 2* software (Busan and Weeks, 2018) for read alignment, mutation counting, and SHAPE reactivity profile generation. The *--random-primer-len 9* option was used to mask RT primer sites with all other values set to defaults. For RNP-MaP library analysis, the protein:RNA mixture samples are passed as the *--modified* samples and no-protein control RNA samples as *--unmodified* samples. Median read depths of all SHAPE-MaP and RNP-MaP samples and controls were greater than 50,000 and nucleotides with a read depth of less than 5000 were excluded from analysis.

#### Secondary structure modeling

Secondary structure models were taken from our previous publication (Iserman et al., 2020).

#### RNP-MaP reactivity analysis

A custom RNP-MaP analysis script (Weidmann et al., 2021) was used to calculate RNP-MaP “reactivity” profiles from the *Shapemapper 2* “profile.txt” output. RNP-MaP “reactivity” is defined as the relative MaP mutation rate increase of the crosslinked protein-RNA sample as compared to the uncrosslinked (no protein control) sample. Nucleotides whose reactivities exceed reactivity thresholds are defined as “RNP-MaP sites”. RNP-MaP site densities were calculated over centered sliding 15-nt windows to identify RNA regions bound by N-protein. An RNP-MaP site density threshold of 5 sites per 15-nt window was used to identify “N-protein binding sites” with boundaries defined by the RNP-MaP site nucleotides.

#### Dynamic light scattering

Dynamic light scattering (DLS) measurements were performed at 25 °C using a Wyatt DynaPro temperature-controlled Plate Reader (Wyatt Technology, Santa Barbara, CA). Samples for the DLS system were prepared in the droplet buffer and filtered through 0.02 mm Whatman Anotop sterile syringe filters (GE Healthcare Life Sciences, Pittsburgh, PA) into a 96-well plate (Wyatt Technology, Santa Barbara, CA). Samples were incubated for 20 min at 25°C before testing. 10 acquisitions were taken, and the results presented represent the mean Rh of the sample.

#### Genome N-protein motif analysis

YYAAAY motifs were counted throughout the NC_045512.2 reference genome (with overlapping motifs counted separately) and the motif counts in each 1000 base pair window were plotted as a histogram. The density of double-stranded RNA was plotted using a kernel density estimation plot with smoothing parameter set to 100. RNA structure data was taken from (Lan et al., 2021).

#### Data and materials availability

All data are available upon request from C.A.R. or A.S.G.

